# A matter of salt: global assessment of the effect of salt ionic composition as a driver of aquatic bacterial diversity

**DOI:** 10.1101/2024.03.09.584225

**Authors:** Attila Szabó, Anna J. Székely, Emil Boros, Zsuzsanna Márton, Bianka Csitári, Natalie Barteneva, Dóra Anda, Péter Dobosy, Alexander Eiler, Stefan Bertilsson, Tamás Felföldi

## Abstract

While the strong general effects of salinity on microbial diversity are well-known and described for marine and freshwater habitats, the impact of the specific composition of major inorganic ions remains largely unexplored. In this study, we assess how microbial community structure in inland saline aquatic habitats is influenced by ionic composition as compared to salinity, spatial factors, and other environmental parameters. We collected and analysed 16S rRNA gene V4 and V3-V4 amplicon datasets from freshwater to hypersaline aquatic environments worldwide (in total 375 samples from 130 lakes). With an emphasis on saline inland waters characterised by highly variable ionic composition, we demonstrated that the ionic composition of the major ions explained more variability in community composition than bulk salinity and that the geographic location of the sampling sites had only an ambiguous effect. We also identified the taxa contributing the most to the observed dissimilarity between communities from sites with different ionic composition and found mostly lineages known to be characteristic for a given habitat type, such as Actinobacteria acI in freshwater, Halomonadaceae in saline, or Nitriliruptorales in soda and soda-saline habitats. Many of these habitat type-specific indicator lineages were monophyletic, underpinning ionic composition as a crucial eco-evolutionary driver of aquatic microbial diversity.

---

External osmotic stress constitutes one of the most cardinal environmental selective factors for aquatic bacteria and drives adaptive evolutionary processes and the emergence of diversity [1-3]. This makes salt content a primary classifying parameter for microbial aquatic habitats. Still, while thalassic saline systems (i.e., waters of marine origin) are uniform in their ionic composition (i.e., predominantly Na^+^ and Cl^-^) and can therefore be adequately characterised by overall salt concentration [4, 5], athalassic (i.e., non-marine) saline waters vary dramatically with respect to ionic composition and can thus be further classified based on the relative ratio of their major ions (Na^+^, K^+^, Ca^2+^, Mg^2+^, Cl^-^, SO_4_^2-^, HCO_3_^-^, CO_3_ ^2-^) [6]. The fact that salinity stress imposed by different ions requires different adaptive mechanisms [7] underscores the significance of ionic composition as a critical selective force for microorganisms [8]. Nevertheless, the influence of differences in ionic composition on microbial diversity and community composition has not been systematically explored, in contrast to the extensive body of literature on how salinity influences microbial diversity [e.g., 9-11] or how their community composition is structured along salinity gradients in thalassic systems [e.g., 12-13].

To address this knowledge gap, we analyse bacterial 16S rRNA gene amplicon datasets from freshwaters and a diverse collection of athalassic saline aquatic systems, including soda lakes known for a dominant contribution of Na^+^ and HCO_3_^-^+CO_3_^2-^ along with elevated pH, high productivity [14] and often extremely high dissolved organic carbon (DOC) content [15]. By analysing these datasets, we test the hypothesis that ionic composition plays a major role in structuring bacterial communities.

The analysed high-quality amplicon sequences were generated from 375 samples obtained from 130 sites located across ten geographic regions on four continents (Africa: 4, Asia: 84, Europe: 262, and North America: 25) (Fig. 1A, Supplementary Figure S1). Thirty-five samples were collected from freshwater lakes (PSU < 1.0), while 340 samples were from athalassic saline lakes (PSU 1.0 - 248). Based on their anionic composition, the saline sites could be further classified according to Boros and Kolpakova [6] as soda lakes (n = 246, Na^+^-HCO_3_^-^-CO_3_^2-^), soda-saline lakes (n = 28, Na^+^-Cl^-^ or Na^+^-SO_4_^2-^ > HCO_3_^-^-CO_3_ ^2-^ > 25 e%) or saline lakes (n = 66, Na^+^-Cl^-^ or Na^+^-SO_4_^2-^) (Fig. 1E, F, Supplementary Figure S2, Supplementary Table S1). Details on the study sites, applied methods, and the sequence data are provided in the Supplementary Material and Supplementary Table S1.

**Fig. 1.**
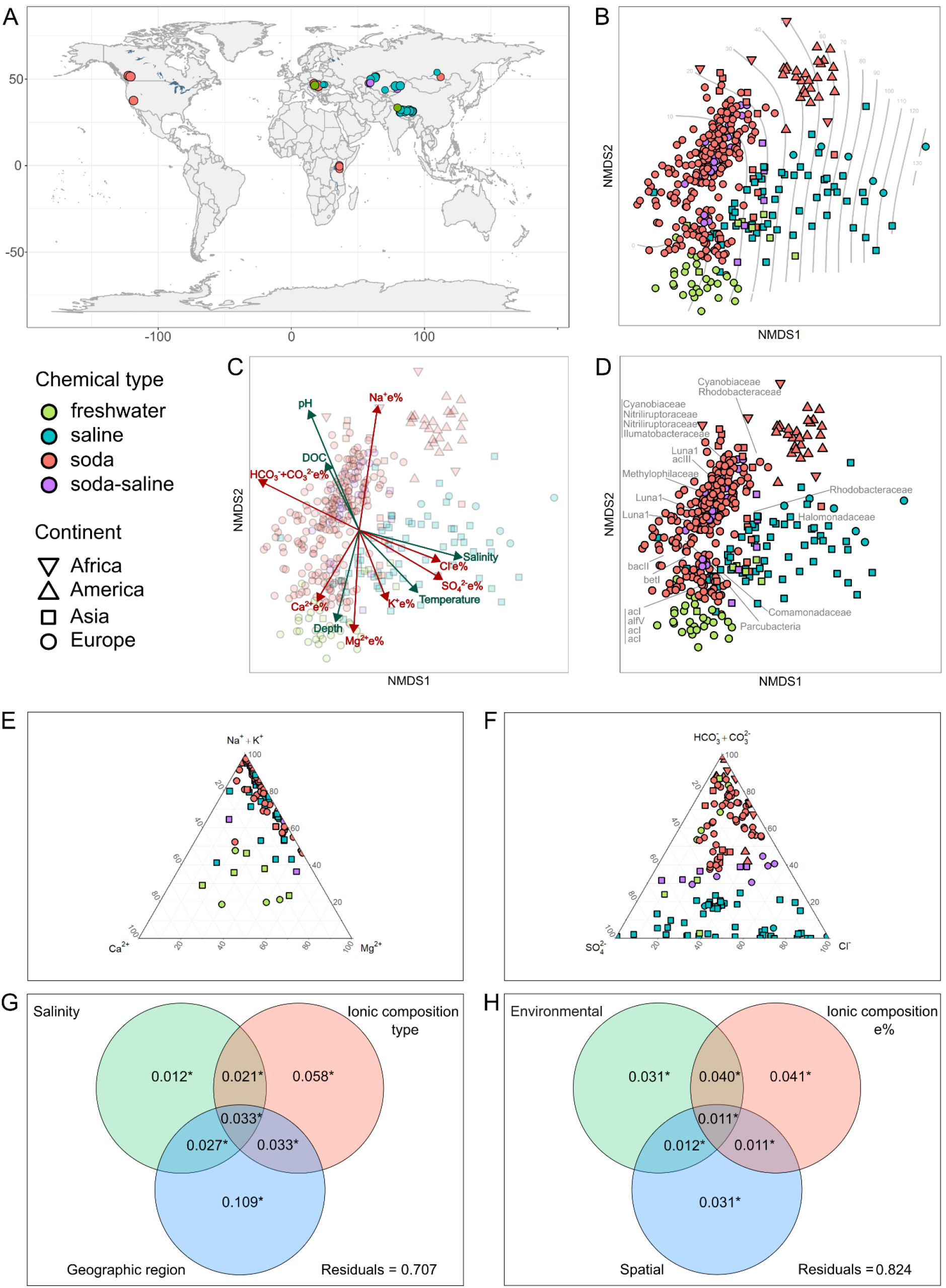
Comparison of planktonic freshwater and saline inland water bacterial communities. A: Map showing the geographic origins of the samples, B-D: NMDS ordination of bacterial OTUs (stress 0.19), defined at 99% sequence identity for the V4 region of the 16S rRNA and rotated along the salinity gradient. Samples are colour-coded according to ionic composition types. B: Salinity (in g/L) is projected as grey contours. C: Significantly fitted (p < 0.05) environmental variables are represented by green vectors, while significantly fitted ion equivalent percentages are depicted in red. D: OTUs responsible for 20% dissimilarity between ionic composition types were identified according to SIMPER analysis and shown in grey by the name of the closest affiliated taxa (family level or above). E-F: Ternary diagrams of the equivalent percentage (e%) contribution of dissolved ions to total ion content in the samples. G-H: Variance in bacterial community composition explained by ionic composition and other parameters. G: Variance significantly explained by salinity, ionic composition types, and geographic regions based on R^2^ of PERMANOVA tests (p < 0.001). H: Variance explained according to variance partitioning analysis by significant environmental parameters (salinity, pH, and sampling depth), ion equivalent percentages, and geographic distance represented by positive significant eigenvectors derived from a distance-based Moran’s eigenvector maps (dbMEM) analysis.

Despite the global geographic distribution of the sampling sites, bacterial community composition was primarily structured in accordance with the specific ionic composition of the waters (Fig. 1, Supplementary Fig. S3, S5). The communities were also structured along the salinity gradient (Fig. 1B, C) with salinity as an additional significant community structuring factor according to the PERMANOVA, but only explaining 1.2% of the variance in community composition (Fig. 1G). Other environmental factors such as pH, DOC, water temperature and depth were also significantly fitted on the NMDS plot and according to the variance partitioning analysis (VPA), the three environmental parameters reported for all samples (salinity, pH and sampling depth) explained 3.1% of the variance (Fig. 1H). The geographic region of the sampling sites explained most of the variance in the PERMANOVA analysis (10.9%, Fig. 1G) but this finding was not corroborated by the VPA, where spatial vectors explained merely 3.1% of the variance (Fig. 1H), or by the distance decay analyses (Fig. S7) where the similarity between time series samples from the same sampling location was often lower than the similarity of samples from distant sites, reflecting previously reported high seasonal turnover of bacterial communities in these habitats [16, 17]. A likely explanation for the contrasting results from the PERMANOVA could be that samples from the same region were often processed within the same project or by the same group and that regional differences were in fact reflecting variations in sampling and sample processing (e.g., sample collection, DNA extraction, PCR amplification) and not true spatial processes such as dispersal limitation.

Meanwhile, all major ions (Na^+^, K^+^, Ca^2+^, Mg^2+^, Cl^-^, SO_4_^2-^, HCO_3_^-^, CO_3_^2-^) were significantly fitted on the NMDS plot, with soda and soda-saline lake communities aligned along the Na^+^, and HCO_3_^-^+CO_3_^2-^ e% vectors, and with saline lakes aligned with the Cl^-^ and SO_4_^2-^ e% vectors (Fig. 1C). The ionic composition of the samples also explained a consistently high percentage of the community variance regardless of whether it was considered as ionic composition type (PERMANOVA: 5.8%) or as ion equivalent percentages (VPA: 4.1%).

The principal taxa underscoring the observed community differences were distinguished by acI-related actinobacteria lineages for freshwaters and Halomonadaceae for saline samples. Both of these are well-known groups characteristic for the respective habitats [18, 19]. Soda and soda-saline lakes were distinguished by other planktonic actinobacteria (e.g., acIII, Luna1, Nitriliruptoraceae) as well as Cyanobiaceae, Methylophilaceae and Rhodobacteraceae, all of which are lineages previously described from these types of habitats (16, 20, 21) (Fig. 1D, Supplementary Fig. S5). Some of the indicator OTUs of saline and freshwater habitats were monophyletic, exemplified in freshwaters by the bet1 lineage within Proteobacteria and Frankiales, and for saline waters by Flavobacteriaceae, Idiomarinaceae, and Alteromonadaceae (Fig. 2). In contrast there were no monophyletic lineages that exclusively contained taxa from soda or soda-saline lakes, but several with representatives from both soda or soda-saline types (i.e., Nitriliruptorales, Rubritaleaceae, Gemmatimonadota, and Cyclobacteriaceae) (Fig 2).

**Fig. 2.**
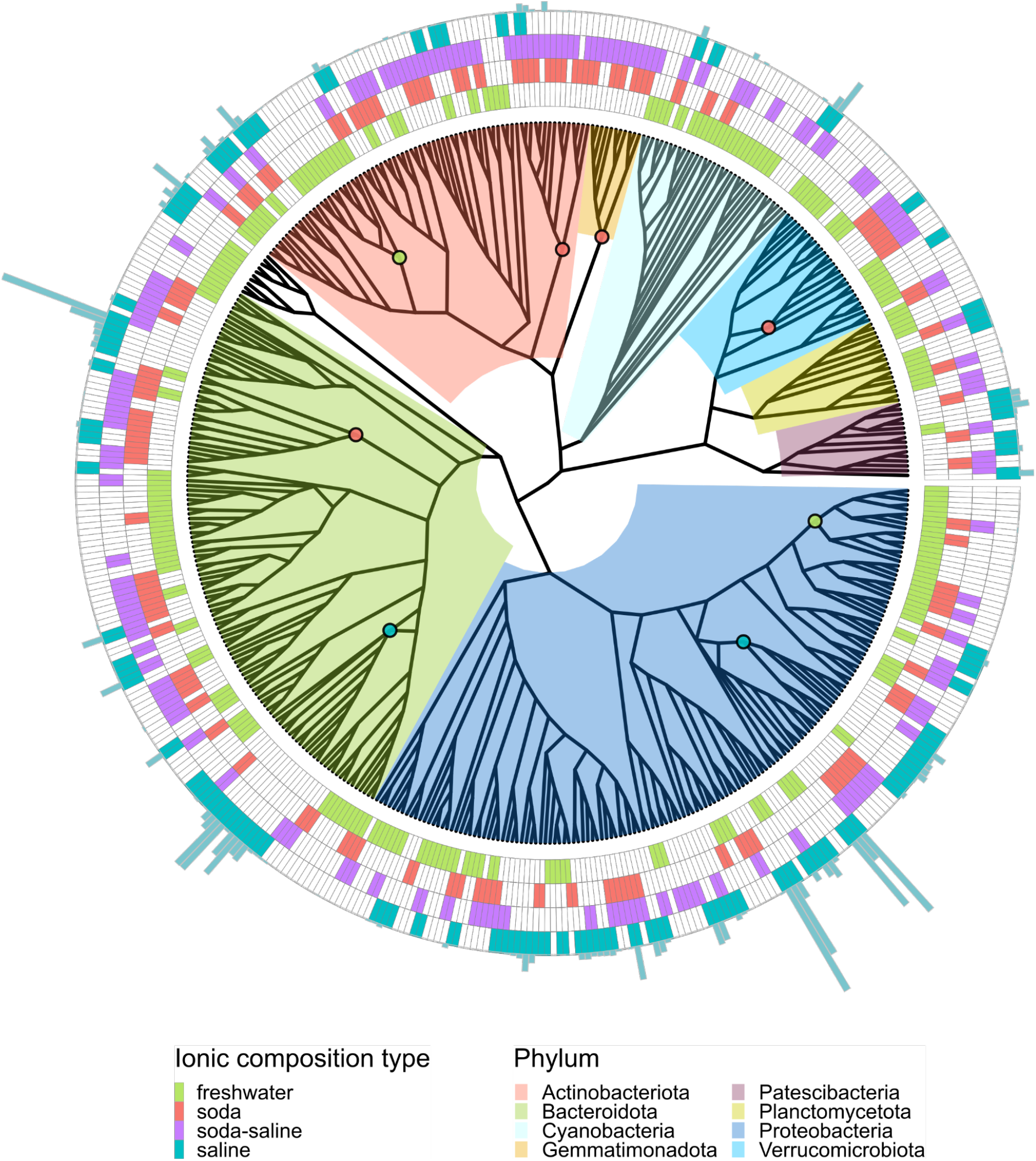
Phylogenetic tree of the OTUs indicative of different ionic composition types of inland waters. Colours in the tree and heatmap correspond to bacterial phyla and ionic composition types, respectively, as detailed in the figure legend. Coloured inner nodes represent monophyletic lineages associated with distinct ionic composition types (> 70% of indicator OTUs align with a given environment type (freshwater, soda and soda-saline, saline), where less than 20% of the indicator OTUs are affiliation with any other category. Key lineages include betI (Proteobacteria) and Frankiales (including acI, Actinobacteriota) for freshwater, Nitriliruptorales (Actinobacteriota), Cytophagales (Bacteroidota), Gemmatimonadota, and Verrucomicrobiales (Verrucomicrobiota) for soda and soda-saline, and Flavobacteriaceae (Bacteriodota) and Enterobacterales (Proteobacteria) for saline aquatic habitats. The heatmap in the outer circle illustrates the ionic composition type associated with each indicator OTU. The mean salinity of samples, weighted by the abundance of each indicator is represented by greenish-blue bars in the outermost circle.

Our comprehensive analysis of a large number of 16S rRNA gene amplicon datasets supports our hypothesis that the composition of major ions is a decisive factor that structures microbial communities and in athalassic habitats, it may even surpass the importance of overall salinity. Samples with different ionic composition were distinguished by characteristic microbial taxa that often represented monophyletic lineages. This indicates that ionic composition is a strong selective force with evolutionary implications [1, 7]. Our findings can serve as a stepping stone for further studies into the genomic and physiological adaptation to different ionic stress and underscore the importance of considering ionic composition besides salinity, particularly when studying saline systems of non-marine origin.

## Acknowledgements

The authors are thankful for the assistance in sample collections to Aidyn Abilkas, Zarina Inelova, Kristóf Korponai, Gergely Krett, István Máthé, Csaba Romsics and Tamás Sápi. Processing of the samples were aided by Marina Ádler, Balázs József Nagy, Izabel Siszler and Sára Szuróczki. Csaba Vad advised on statistical analyses. The authors express gratitude to Károly Márialigeti for providing funding for a part of the project.

## Data Availability Statement

Detailed information on sequence sets’ accession, available metadata and the read processing pipeline is given in Supplementary Table S1 and Supplementary Methods. Detailed scripts of data analysis and for reproducing the manuscript’s figures were uploaded to the GitHub repository ‘MatterOfSalt’ (https://github.com/attiszabo/MatterOfSalt).

## CRediT

Conceptualization

AS, AJS, TF

Data Curation AS, DA

Formal Analysis

AS

Funding Acquisition

AJS, EB, NB, SB, TF

Investigation

AS, AJS, EB, ZM, BC, NB, DA, PD, AE, TF

Methodology

AS

Project Administration

AS, AJS, EB, TF

Resources

AJS, EB, NB, AE, SB, TF

Software

AS

Supervision

AS, AJS, SB, TF

Validation

AS, DA

Visualisation

AS

Writing – Original Draft Preparation

AS, AJS, TF

Writing – Review & Editing

EB, ZSM, BC, NB, DA, PD, AE, SB

