## Supplementary Text and Supplementary Figures for "A matter of salt: global assessment of the effect of salt ionic composition as a driver of aquatic bacterial diversity"

The supplementary text content includes detailed sample site descriptions, material and methods of the study and code snippets for V4 and V3-V4 16S rRNA gene amplicon data analysis.

**S1 Data collection**

To ensure comprehensive data collection, we utilised a search strategy that involved querying the NCBI, EBI, and IMG/M databases with a set of relevant terms "soda lake", "saline lake", "alkaline lake", "soda pan", "soda", "saline", and "alkaline". Additionally, we manually reviewed the results and searched for peer-reviewed publications that referred to sequence data in Google Scholar and Web of Science databases. Our analysis was primarily based on sequence sets obtained by amplifying the V4 and V3-V4 regions of the bacterial 16S rRNA gene, as they were the most common. From the hits evaluated as of 15th November 2022, we excluded entries without available sequences, lacked proper metadata description, or contained too few sequences (< 2500 reads) after pipeline processing. For deeper water bodies, we included only sequences obtained above the chemo- and oxycline. To supplement the dataset, we also generated additional datasets from various aquatic environments sampled in the period 2012 to 2021 by amplifying and sequencing the V3-V4 gene region of the 16S rRNA gene.

**S2 Site information**

Detailed metadata available for the samples included in the analysis are provided in Supplementary Table 1. Several sites could be regarded as ‘true’ lakes (with distinct shoreline habitat) while others are pans or ponds in the strict limnological sense. Due to patchy or incomplete literature data, such detailed categorization was not unequivocally possible for all sites and was accordingly not applied to the studied aquatic habitats. However, where possible, the information has been added to Supplementary Table S1 (column ‘Site description’).

**Africa**: Great Rift Valley soda lakes

The most famous soda lakes in the world are probably those in the East African Rift Valley. A comprehensive overview of these lakes can be found in the book by Schagerl (2016). For detailed information on the chemical composition of these lakes, refer to the recent study conducted by Lameck et al. (2023). However, it is worth noting that the availability of bacterial amplicon data from these sites is limited. In our study, we included 16S rRNA gene data from five water samples collected from Kenyan soda lakes, namely Bogoria, Elmenteita, Magadi, and Sonachi. These data have been deposited under BioProject PRJNA187566 in the NCBI database. Due to the low number of sequence reads, the two available samples from Lake Bogoria water were merged in the analysis.

**Asia:** Kazakhstan

Kazakhstan encompasses an area approximately twice the size of Western Europe, and boasts a remarkable array of permanent and intermittent aquatic habitats and wetlands, covering an estimated 37% of the country's total land area (Boros et al., 2017a). Urivaev (1959) estimated the steppe region of Central Kazakhstan alone harboured 14 000 shallow freshwater and saline lakes. However, the impact of anthropogenic activities over the past decades has likely resulted in a decline in the abundance of these lakes. To the best of our knowledge, only very limited data is currently available on the planktonic prokaryotic communities of the saline-alkaline lakes in this region.

To bridge this knowledge gap, we recently conducted extensive sampling campaigns in various semi-arid and arid regions of Kazakhstan. The goal of these campaigns was to identify soda lakes in Kazakhstan and to reveal the microbial community composition of previously uncharacterized aquatic habitats.

*Northern Kazakhstan*

Water samples were collected during April-May 2015 to investigate the bacterial community composition of several steppe lakes. The analysis included 16S rRNA gene amplicon data from two freshwater and twelve saline lakes, encompassing a salinity range of 0.5 to 149 g/L and pH values ranging from 7.9 to 9.5. For detailed information on the sampled lakes, refer to Boros et al. (2017a).

*South East: Balkhash-Alakol depression*

Lake Balkhash is a large, shallow, and slightly saline lake, ranking as the third largest lake in Eurasia in terms of area. Stretching approximately 600 km in length and varying between 9 and 75 km in width, Lake Balkhash covers a total surface area of 18 210 $km^2$. It consists of two distinct basins, separated by the Uzunaral Strait: a freshwater basin in the West and a saline, deeper water basin in the East. Despite its size, the lake is shallow, with the Western Basin reaching a maximum depth of 11 m and a maximum depth of 26 m in the Eastern Basin. The salinity of Lake Balkhash ranges from 0.2 to 5.0 g/L (Sala et al., 2020).

Adjacent to Lake Balkhash, Lake Alakol is another important saline lake in the region. It covers an area of 2 650 $km^2$ and measures 104 km in length from northwest to southeast, with a width of 52 km. Lake Alakol surpasses Lake Balkhash in-depth, with an average depth of 22 m and a maximum depth of 54 m. Salinity varies with the lake axis and with depth, ranging between 1.1 and 10 g/L (Aladin & Plotnikov, 1993).

In our analysis, we included seven samples collected from different parts of Lake Balkhash during a monitoring campaign in September 2018.

Furthermore, an additional 27 samples were obtained during a sampling campaign in 2021, focusing on the southeastern shore of Lake Balkhash, Lake Alakol, and the surrounding saline, alkaline lakes. Based on the ionic composition of the water, soda and soda-saline lakes were identified alongside the saline sites.

*West Kazakhstan (Aktobe Region) and South Kazakhstan (Zhambyl Region)*

To identify soda lakes in the Aktobe and Zhambyl regions of Kazakhstan, a sampling campaign was conducted during 2018 and 2019. The investigation focused on saline-alkaline water bodies located near the towns of Shalkar (north of the Aral Sea), Belkopa, and Akkol in the southern region. These sampling sites are situated in the Kazakh semi-desert and Central Asian northern desert regions, characterised by semi-arid and arid climates. For our analysis, nine samples were included. The salinity of these samples ranged from 1.0 to 248 g/L, representing a diverse range of saline conditions within the investigated alkaline lakes.

**Asia**: Transbaikalia

*Barguzin Valley*

As part of our analysis, we included the 16S rRNA gene sequences obtained from a water (brine) sample collected from Lake Gudzhirganskoe. This alkaline lake is located in the valley of the Barguzin River, situated to the east of Lake Baikal, as described by Lavrentyeva et al. (2020).

*Lake Doroninskoe*

Lake Doroninskoe is a meromictic soda lake with a depth of approximately 6 meters. The lake is characterised by a pH level of around 10 and a salinity of approximately 27 g/L. The ionic composition of the water reflects the typical characteristics of soda lakes (Gorlenko et al., 2010; Matyugina et al., 2018). Six water samples from 2.5, 3.0 and 3.15 m depth were included in our analysis. These samples were collected during a diurnal campaign in September 2013, as described by Matyugina et al. (2018).

**Asia**: Tibetan Plateau

Ji et al. (2019) conducted a study to investigate the bacterial community composition of 25 lakes located on the Tibetan Plateau, situated at altitudes ranging from 4 280 to 4 856 m above sea level. The sampling took place during the summer of 2015, focusing on 6 freshwater and 19 saline lakes with salinity levels ranging from 0.1 to 118 g/L. In our analysis, we utilised the 'Replicate ID 1' samples from the dataset provided by Ji et al. (2019). Additional physicochemical data for the sites was obtained from Yue et al. (2019). References for water ion composition can be found in Supplementary Table 1. Samples from lakes Baima Nam Co, Guogen Co, Gyarab Punco, Norma Co, and Wuma Co (Darab Co) were excluded from our analysis due to inaccessible ionic composition data.

**Europe:** Carpathian Basin, Pannonian Steppe

The Carpathian (or Pannonian) Basin is a broad flat alluvial basin in Central Europe shaped by two major rivers (Danube and Tisza) and bordered by high mountain ranges (Carpathians, Alps, and Dinarides). Within this region lies the Pannonian Steppe or Puszta, one of the largest grasslands in Europe, stretching from Austria to Serbia and covering a significant portion of Hungary. The Pannonian Steppe was once a forest-steppe area with extensive wetlands, marshes, and lakes formed by regular river floods. It is characterised by soda lakes, pans (shallow lakes with a high area-to-volume ratio), and a few saline lakes. Lake Neusiedler/Fertő and Lake Velence are the largest lakes in this area, complemented by several shallow and some intermittently dry pans. These alkaline and saline lakes typically feature salinity levels in the hyposaline range (3-20 g/L) and pH above 9. The water in most soda lakes and pans is turbid and grey due to the presence of inorganic clay particles, but at some sites instead shows brown-colour due the high humic content. Detailed information about the origin, nature, and conservation of these saline aquatic habitats can be found in the works of Boros et al. (2013, 2014, 2017b).

In recent years, several sampling campaigns have been conducted in soda lakes and pans of the Pannonian Steppe to investigate the spatiotemporal changes in bacterioplankton community structure. Various studies (Márton et al., 2023; Sinclair et al., 2015; Szabó et al., 2017, 2020; Szabó et al., 2022; Szuróczki et al., 2020; reviewed by Felföldi 2020) have provided valuable insights into these ecosystems. The current study incorporated samples collected from different locations and time periods, including soda pans in the Kiskunság National Park in Hungary (Szabó et al., 2017, 2020), Seewinkel soda pans in Austria (Sinclair et al., 2015), Lake Neusiedler/Fertő in Austria (Szuróczki et al., 2020), and other soda pans in Austria and Hungary (Márton et al., 2023a, 2023b; Szabó et al., 2022). Furthermore, samples were collected from a transect spanning the Pannonian Steppe, including the Seewinkel region in Austria and Vojvodina in Serbia (2018), as well as during a monitoring program in soda and saline lakes in Hungary (2021).

Additionally, surface water samples from Lacul Ursu and Lacul 'Plus' hypersaline lakes in Romania (2015), as well as freshwater sites from the Carpathian Basin (25 samples), were included in the analysis to compare bacterial communities within the same biogeographic region.

**North America**: Cariboo Plateau

Zorz et al. (2019) conducted a comprehensive investigation of microbial mat communities in four alkaline soda lakes located on the Cariboo Plateau in British Columbia, Canada. The Plateau is characterised by a basaltic bedrock formed during volcanic activity in the Miocene and Pliocene eras, which creates favourable conditions for the development of soda lakes due to the decreased levels of soluble calcium and magnesium resulting from volcanic rock weathering. The researchers collected samples from Deer Lake, Goodenough Lake, Last

Chance Lake, and Probe Lake annually from 2014 to 2017, with the lakes exhibiting high pH levels ranging from 10.1 to 10.7 and high salinity levels ranging from 17 to 48 g/L. For our analysis, we incorporated fifteen 16S rRNA gene amplicon libraries that were published alongside their study. High-quality metadata was provided for the years 2015 and 2017, and we extrapolated the metadata from the 2015 year to the years 2014 and 2016 for our analyses (Supplementary Table S1).

**North America**: Mono Lake

Mono Lake is an athalassic hypersaline, alkaline lake situated in California, USA, east of the Sierra Nevada Mountains on the western edge of the Great Basin. It has a high salinity of approximately 90 g/L and a pH of around 9.8. Although the lake is usually monomictic, it can undergo extended periods of stratification, especially following wet winters, resulting in a seasonal oxycline and anoxic conditions in its lower layer (Edwardson & Hollibaugh, 2018). From this lake, we included samples collected from the epilimnion (above the chemocline at 12 m depth) at Station 6 in the south basin in our analysis . After quality filtering, the dataset included 16S rRNA gene amplicon sets from a sample collected at a depth of 10 meters in July 2012 (Edwardson & Hollibaugh 2018) and 9 samples collected at depths of 0, 5, 10, and 12 m from Phillips et al. (2021). From the latter study, triplicate samples from September 2017 were merged for the analysis. Based on ion chemistry data provided by Nielsen & DePaolo (2013), the lake was classified as a soda type. It is important to note that chloride concentrations in the water are nearly as high as carbonates, and the sulphate content was also remarkably high at approximately 17 equivalent percentage (e%). Williams (1998) reported a similar ionic composition, highlighting some variation in anionic composition with changing salinities, although the difference was not substantial.

### S3 Materials and methods

Environmental measurements and the determination of ionic composition type

Our study centered on examining the planktonic bacterial communities of inland waters and their relationship with the dissolved ion content. Freshwater habitats were defined as aquatic environments having salinity lower than 1 g/L (Saccó et al., 2021), while saline sites were categorised according to the framework recently suggested by Boros and Kolpakova (2018). Therefore, sodium brines were classified into the following categories, based on their anion content: (1) soda type if carbonate ions in the water are present with an equivalent percentage (e%, defined as the molar equivalent according to the separated calculation of total cation or anion pools) of more than 25 e%; (2) soda-saline type if carbonate ions are present with >25 e%, but are not the most dominant anions in the water; (3) saline type when chloride or sulphate is the dominant anion and the presence of carbonates are less than 25 e% (Supplementary Fig. S2.

Available metadata are provided in Supplementary Table S1, which provides references for environmental parameters and ionic composition of the sites. For previously unpublished samples (Carpathian Basin and Kazakhstan), temperature, pH, and conductivity parameters were measured on-site with a Hanna HI9033 portable field meter. Salinity data were obtained through on-site conductivity measurements, based on the equation by Boros et al. (2014) estimated for soda lakes (total ions (g/l) = 0.79 * EC (mS/cm)). Alternatively, the mass of dissolved inorganic solids (total dissolved solids (TDS) were employed as a salinity proxy, calculated by summing the concentrations of measured ions in g/l. Concentrations of $Na^{+}$, $K^{+}$, $Ca^{2+}$, $Mg^{2+}$, $Cl^{-}$ and $SO_4^{2-}$ were quantified applying a DIONEX 5000 ICS+ dual channel ion-chromatography (Thermo Fisher Scientific, USA), $HCO_3^{-}$ and $CO_3^{2-}$ by alkalinity titration (Eaton et al., 2005). Equations (1) and (2) (Boros and Kolpakova, 2018) were used to calculate the equivalent percentage of cations and anions. Conventional ion balance percentage calculation was used to validate ion e% calculations by applying a 10% error threshold as described in Lameck et al. (2023). Chlorophyll-a concentration was determined as described by Boros et al. (2017a). TOC concentrations of unfiltered samples and DOC following filtration through membrane having pore size of 0.45 µm were determined using high-temperature combustion with a MULTI N/C 3100 analyzer (Analytik Jena, Germany) in accordance with the international standard MSZ EN 1484:1998.

Analysis of planktonic bacterial community composition of samples collected within this study

*Sample collection*

Asia: Kazakhstan

Surface water samples gathered approximately 10 metres from the shoreline, were collected from the upper 5 cm of the water column using clean, washed beakers or glass bottles. Two replicate samples were filtered through 0.22 µm pore size syringe filters (Merck Millipore) until clogging occurred (10-60 ml), followed by on-site drying. To eliminate residual water, air was pushed through the filters using syringes. The desiccated filters were stored at -20°C until extraction of community DNA content.

Europe: Carpathian Basin

From the saline lakes Ursu and 'Plus' 100-200 ml surface water was filtered through a 0.22 µm pore filter (Sartorius). Water samples from Neusiedler See/Lake Fertő were filtered using a mixed cellulose filter with a pore size of 0.22 µm (type GSWP; Merck Millipore), as detailed in Szuroczki et al. (2020). For freshwater and soda lake samples collected in 2018 approximately 250 ml of the water was filtered through a 0.22 µm pore diameter polycarbonate filter (Merck Millipore). During the 2018 Carpathian Basin transect campaign, samples were first prefiltered through a 20 µm pore-size plankton net, and microbial material was subsequently collected by filtering through a 0.1 µm membrane filter (type GSWP; Merck

Millipore). In the 2021 soda lake monitoring campaign, water samples were prefiltered through a 30 µm pore-size plankton net on-site. From each water sample, 10 mL was filtered through 0.1 µm pore-sized mixed cellulose membrane filters (type GSWP; Merck Millipore).

*Community DNA extraction*

Filters were sectioned using sterile scissors, and total genomic DNA was extracted using the PowerSoil DNA isolation kit (MoBio Laboratories) according to the manufacturer's protocol with the exception that the cell disruption step was carried out by shaking at 30 Hz for 2 min in a Mixer Mill MM301 (Retsch, Haan, Germany). For the saline lake (Ursu and 'Plus') and for 2018 freshwater campaign samples, the Ultra Clean Soil DNA Isolation Kit (MoBio Laboratories) was used with a similar protocol.

*Library preparation and amplicon sequencing*

For sampling campaigns carried out in the Carpathian Basin and North Kazakhstan before 2017, the V3-V4 region of the bacterial 16S rRNA gene was amplified using primer pairs BAKT_341F (5'-CCTACGGGNGGCWGCAG-3') and BAKT_805R (5'-GACTACHVGGGTATCTAATCC-3') (Herlemann et al., 2011), and sequenced on a Roche GS Junior platform. DNA extraction, amplification, and the pyrosequencing approach were carried out as described in Szabó et al. (2020).

Samples collected in December 2015 or later were sequenced on the Illumina MiSeq platform, and library preparation utilised the same primer pair with a slightly different reverse primer for wider coverage, Pro805R (5'-GACTACNVGGGTATCTAATCC-3', Takahashi et al., 2014), with Fluidigm CS1 and CS2 universal sequences at their 5' ends, but amplifying the same region of the bacterial 16S rRNA gene. The PCR amplification process was conducted in triplicate in a final volume of 20 µL containing 1× Phusion HF Buffer (Thermo Fisher Scientific Inc., Waltham, MA, USA), 0.2 mM dNTPs (Fermentas Vilnius, Lithuania), 0.4

µg/µL Bovine Serum Albumin (Fermentas), 0.3 µM of each primer, and 0.4 U Phusion High-Fidelity DNA Polymerase (Thermo Fisher). The following thermal cycling conditions were applied: initial denaturation at 98 °C for 5 min, followed by 25 cycles of denaturation (95 °C for 40 s), annealing (55 °C for 2 min), and extension (72 °C for 1 min), with a final extension step at 72 °C for 10 min. The triplicate PCR products were pooled, normalised, and then subjected to additional library preparation steps and sequencing at the Genomics Core, RTSF, Michigan State University (USA). Paired-end sequencing was carried out on an Illumina MiSeq platform using a MiSeq Reagent Kit v2 (2 × 250 bp).

An exception includes samples from the seasonal soda lake campaign in the Kiskunság National Park (2017) and those from the Pannonian Steppe transect campaign (2018), processed and amplified according to the following protocol (https://www.protocols.io/view/sample-preparation-for-illumina-miseq-dual-index-a-yxmvm769ov3p/v1) (Vass et al. 2020) at Uppsala University.

*Amplicon data analysis*

This analysis encompasses two datasets:  a V4 dataset with broader representation due to its prevalence in NCBI SRA, and a V3-V4 dataset with fewer samples but enabling a finer taxonomic resolution. Since sequence data were obtained by different sequencing platforms and chemistries, the mothur tool (Schloss et al., 2009) was applied to process the raw sequences using a custom-created pipeline to obtain a unified high-quality OTU table using a 99% sequence identity threshold due to the wide variation in sequence quality among datasets originating from diverse sources and aiming to mitigate potential diversity estimation biases (Johnson et al., 2019; Schloss 2021). Sequence sets used in the comparative analysis can be obtained through the NCBI SRA database; accessions are given in Supplementary Table 1. Detailed descriptions of the different sequence processing steps are provided in S3 Supplementary scripts of the bioinformatics analysis and under the GitHub repository 'MatterOfSalt' (https://github.com/attiszabo/MatterOfSalt).

The initial steps of sequence processing included quality filtering and the removal of adapters, barcodes, and primers, according to the platform-specific best practices of each sequencing data set. After the preliminary processings, datasets generated on different sequencing platforms were merged. For the alignment of sequence reads, the ARB-SILVA SSU Ref NR 138 reference database (Quast et al., 2013) was used. Denoising was performed using mothur's pre.cluster command using the default algorithm (Huse et al., 2010) and applying the suggested 1 bp difference per 100 bp cutoff (Schloss et al., 2011). Detection and elimination of chimeric sequences were carried out using the mothur-integrated version of VSEARCH. Taxonomic assignment relied on the Silva SSU 138 reference database using a minimum bootstrap confidence score of 80. Bacterial OTUs assigned to non-primer specific taxonomic groups (e.g. Archaea, chloroplasts, mitochondria, unknown) were removed from the dataset. OTU assignment was performed using the OptiClust algorithm (Westcott et al., 2017) at 99% similarity threshold. The TaxAss software (Rohwer et al., 2018) was employed to enhance taxonomic classification, utilising FreshTrain (2020 June 15 release) database as a reference for inland aquatic habitat bacterial lineages, and the Silva SSU 138. Prior to statistical analyses, subsampling was conducted to rarefy reads with the lowest sequence count among samples (n = 2558).

<u>Ordination and statistical analyses</u>

Statistical analysis and visualisation were conducted using R 4.3.0 software (R Core Team, 2023). The 'ggplot2' package (Wickham 2016) was utilised for data visualisation. Ternary plots were made using the 'ggtern' package (Hamilton & Ferry, 2018). Nonmetric multidimensional scaling (NMDS) ordinations and 'envfit' analyses were carried out using the 'vegan' package (Oksanen et al., 2022) 'metaMDS' function. Salinity values were fitted to the ordination plot with 'ordisurf'. Procrustes test, implemented in the 'vegan' package, was employed to assess the dissimilarities in the ordination patterns between the V4 and V3-V4 datasets. Similarity percentage (SIMPER) analysis based on Bray-Curtis similarity was carried out with the PAST3 software (Hammer et al., 2001) to determine which OTUs were responsible for the dissimilarity among ionic composition types used in this study. To test for significant differences in planktonic bacterial community composition between different salinities, water chemical types and among geographic regions, three-way PERMANOVA tests were carried out with the 'adonis2' function of vegan. To investigate distance-decay relationships, Bray-Curtis dissimilarity was employed as a metric to quantify the dissimilarity between ecological communities based on their abundance profiles. The calculated dissimilarity values were then transformed into community similarity values by subtracting them from 1, enabling the examination of community similarity as a function of geographic distance. Geographical distances between sampling sites were calculated using the Haversine formula, accounting for the curvature of the Earth's surface, from the 'geosphere' library (Hijmans et al., 2017). A Mantel test was then conducted to test the significance between community similarity and geographic distance in R. The relative contributions of environmental, spatial, and methodological factors to planktonic bacterial community similarity were assessed through variance partitioning analyses. Subsampled OTU abundances were transformed using the Hellinger method via 'decostand.' Only those environmental parameters were considered (salinity, pH, sampling depth) that were available for all samples assessed in the study. The impact of chemical types was evaluated based on the equality percentages of major dissolved ions. Samples with any missing dissolved ion e% information (n = 47) were excluded from the analysis. Collinear variables ($Mg^{2+}$, $SO_4^{2-}$, $HCO_3^-$+$CO_3^{2-}$) were detected using variance inflation factors (VIF) analyses with the 'vif.cca' command and were excluded from the analyses. Variables deviating from normality based on Shapiro-Wilk tests were transformed (untransformed: $HCO_3^-$; square root: $Cl^-$; $\log_{x+1}$: salinity, pH, $Na^+$, $K^+$, $Ca^{2+}$, $CO_3^{2-}$; inverse, 1/x: sampling depth, ) and z-score standardised. Stepwise selection was used to select environmental and ion e% parameters using the 'ordistep' function (direction="both", perm.max=200, pstep=999). To test the effect of spatial scale on the community composition, Moran's Eigenvector Maps (MEM) were constructed based on the spatial distance matrix created from sampling site coordinates using the dbMEM function from the ‘adespatial’ package (Dray et al., 2023) and positive MEMs were kept in the subsequent analysis. Stepwise selection was carried out for the three positive MEMs as described for the environmental parameters. Variance partitioning was performed utilising the 'varpart' function and tested for significance with the ‘anova’ function.

Indicator species analysis was conducted using the 'indicspecies' R package (De Cáceres & Legendre, 2009) to identify abundant OTUs (present in at least three different samples and constituting < 1% relative abundance in one sample) characteristic of ionic composition types. A phylogenetic tree (Fig. 2) was constructed for indicator OTUs using ‘clearcut’ (Sheneman et al., 2006) implemented in the mothur program and visualised with the ‘ggtree’, and ‘ggtreeExtra’ packages in R. Monophyletic lineages were identified in association with specific ionic composition types, with more than 70% of indicator OTUs aligning with a particular environment type (freshwater, soda, soda-saline, saline), while indicating less than 20% association with another category. The scripts for statistical analyses and data visualisation (Fig 1., 2. and Supplementary Figures) can be found in the GitHub repository:
https://github.com/attiszabo/MatterOfSalt/.

### S4 Supplementary scripts of the bioinformatic analysis

Since sequence data were obtained by different sequencing platforms and chemistries, the mothur tool (Schloss et al., 2009) was applied to process the raw sequences using a custom-created pipeline to obtain a unified high-quality OTU table using 99% similarity cutoff (see Supplementary Methods). These mothur scripts outline the pipeline for processing V4 and V3-V4 16S rRNA gene amplicon data using mothur v1.44.3. However, the preprocessing of 454 pyrosequencing reads was carried out with mothur v1.35. This script also can be found in the GitHub repository: https://github.com/attiszabo/MatterOfSalt/.

**Sequence set SRA accesions used in this study are listed in Supplementary Table 1.**

> ⓘ Note
>
> There are a few differences from mothur version 1.47 on the handling and creations of the name, count and group files

#### Preprocessing

##### Preprocess 454 pyrosequencing reads

*Convert SFF to fasta and quality files*

```
sffinfo(sff=run_name.sff)
```

*Trim flowgrams*

```
trim.flows(flow=run_name.flow, oligos=run_name_barcodes_341F_785R.oligos,
minflows=350, maxflows=700, pdiffs=1)
  ##run_name_barcodes_515F_806R.oligos used for the V4 dataset,
run_name_barcodes_341F_805R.oligos for the V3-V4 dataset
```

*Denoise flowgrams*

```
shhh.flows(file=run_name.flow.files)
```

*Trim barcode, primers and quality filter*

```
trim.seqs(fasta=current, name=current,
oligos=run_name_barcodes_341F_785R.oligos, pdiffs=1, bdiffs=0, maxhomop=8,
flip=T)
  #run_name_barcodes_515F_806R.oligos used for the V4 dataset,
run_name_barcodes_341F_805R.oligos for the V3-V4 dataset
```

*oligos file*(https://mothur.org/wiki/oligos_file/):
barcode *SAMPLE1_BARCODE* SAMPLE1
barcode *SAMPLE2_BARCODE* SAMPLE2
barcode *SAMPLE3_BARCODE* SAMPLE3
forward CCTACGGGNGGCWGCAG 341F
reverse GACTACHVGGGTATCTAATCC 785R
forward GTGNCAGCNGCCGCGGTAA 515F

reverse GGACTACNNGGGTNTCTAAT 805R

*Summarize sequences*

```
summary.seqs(fasta=current, name=current)
```

### Using mothur v1.44.3:

*Find and list unique sequences*

```
unique.seqs(fasta=current)
```

*Create a count_table*

```
count.seqs(name=current, group=current)
```

*Summarize sequence and group (sample) information*

```
summary.seqs(fasta=current, count=current)
count.groups(count=current)
```

#### Preprocess pair-end Illumina sequences

*List samples and sequence files*

```
make.file(inputdir=., type=gz, prefix=project_name)
```

*Combine paired reads of forward and reverse direction*

```
make.contigs(file=project_name.files, deltaq=10)
```

*Summarize sequences*

```
summary.seqs(fasta=project_name.trim.contigs.fasta)
```

*Count sequences per samples*

```
count.groups(group=project_name.contigs.groups)
```

*Quality filter reads*

```
screen.seqs(fasta=current, group=current, maxambig=0, minlength=200,
maxhomop=8)
    #'minlength' adjusted 200 for the V4 dataset, 400 for the V3-V4
dataset
```

*Summarize filtered reads*

```
summary.seqs(fasta=current)
```

#### Preprocess non-paired-end Illumina sequences

*Create a fasta and quality file from fastq*

```
fastq.info(fastq=SRRxxxxxxx.fastq)
rename.file(fasta=current, qfile=current, prefix=sample1)
```

*Create a group file and count reads per sample*

```
make.group(fasta=sample1.fasta-sample2.fasta-sample3.fasta,
groups=sample1-sample2-sample3)
count.groups(group=current)
```

*Merge fasta files within the same project*

```
merge.files(input=sample1.fasta-sample2.fasta-sample3.fasta,
output=non_paired_project_name.fasta)
summary.seqs(fasta=current)
```

*Quality filter reads*

```
screen.seqs(fasta=current, group=current, maxhomop=8, maxambig=0,
minlength=200, maxlength=300)
    #'minlength' adjusted to 200 for the V4 dataset, 400 for the V3-V4
dataset
    #'maxlength' adjusted to 300 for the V4 dataset, 500 for the V3-V4
dataset
summary.seqs()
```

*Primer removal from Illumina datasets*

```
trim.seqs(fasta=current, oligos=515F_806R.oligos, pdiffs=2, checkorient=T)
    #515F_806R.oligos used for the V4 dataset, 341F_805R.oligos for the
V3-V4 dataset
summary.seqs()
# Optionally, if trim.seqs eliminated some of your sequences you need to
list and remove them from your group file
list.seqs(fasta=project_name.trim.contigs.good.scrap.fasta)
remove.seqs(group=project_name.contigs.good.groups,
accnos=project_name.trim.contigs.good.scrap)
# For cases where unidentified adapter residuals or primer sequences
remained, mothur's trim.seqs was used in combination with the MEGA version
6 or 11 (Tamura, Stecher, Peterson, Filipski, and Kumar 2013) on
previously SILVA 138 SSU aligned fasta files
```

*Find and list unique sequences*

```
unique.seqs(fasta=current)
```

*Create a count_table*

```
count.seqs(name=current, group=current)
```

*Summarize sequence and group (sample) information*

```
summary.seqs(fasta=current, count=current)
count.groups(count=current)
```

### Merging datasets

*For the V3-V4 dataset*

```
merge.files(input=V3V4.project_name1.fasta-V3V4.project_name2.fasta-
V3V4.project_name3.fasta…V3V4.project_namen.fasta, output=V3V4.fasta)
merge.count(count=V3V4.project_name1.count_table-
V3V4.project_name2.count_table-
V3V4.project_name3.count_table…V3V4.project_namen.count_table,
output=V3V4.count_table)
summary.seqs(fasta=current, count=current)
count.groups(count=current)
unique.seqs(fasta=current, count=current)
summary.seqs(fasta=current, count=current)
```

*For the V4 dataset*

Ahead of merging, the MEGA software was used to trim the 341-533 region from the pre-aligned V3-V4 dataset, ensuring compatibility with the V4 dataset.

```
merge.files(input=V3V4_533F_trim.fasta-V4.project_name1.fasta-
V4.project_name2.fasta…V4.project_namen.fasta, output=V4.fasta)
merge.count(count=V3V4.count_table-V4.project_name1.count_table-
V4.project_name2.count_table…V4.project_namen.count_table,
output=V4.count_table)
summary.seqs(fasta=current, count=current)
count.groups(count=current)
unique.seqs(fasta=current, count=current)
summary.seqs(fasta=current, count=current)
```

### Processing quality filtered and trimmed, merged datasets (V4/V3-V4)

*Sequence Alignment*

```
align.seqs(fasta=current, reference=/path/Silva_nr138/silva.nr_v138.align)
summary.seqs(fasta=current, count=current)
```

*Removing non-overlapping reads*

```
screen.seqs(fasta=current, count=current, start=13862, end=23440)
    #'start' adjusted to 13862 for the V4 dataset, 6428 for the V3-V4
dataset
summary.seqs(fasta=current, count=current)
count.groups(count=current)
```

*Removing overhanging positions*

```
filter.seqs(fasta=current, trump=., vertical=T)
unique.seqs(fasta=current, count=current)
summary.seqs(fasta=current, count=current)
```

*De-noising reads*

```
pre.cluster(fasta=current, count=current, diffs=2)
  #'diffs' adjusted to 2 for the V4 dataset, 4 for the V3-V4 dataset
summary.seqs(fasta=current, count=current)
count.grouos(count=current)
```

*Chimera check and removal*

```
chimera.vsearch(fasta=current, count=current, dereplicate=T)
remove.seqs(fasta=current, accnos=current)
summary.seqs(fasta=current, count=current)
count.groups(count=current)
```

*Singleton removal*

```
split.abund(fasta=current, count=current, cutoff=1)
summary.seqs(fasta=\*.abund.fasta, count=\*.abund.count_table)
count.groups(count=current)
```

*Taxonomic assignment and the removal of non-primer specific targets*

```
classify.seqs(fasta=current, count=current, reference=silva.nr_v138.align,
taxonomy=silva.nr_v138.tax, cutoff=80, method=wang)
remove.lineage(fasta=current, count=current, taxonomy=current,
taxon=Archaea-Chloroplast-Mitochondria-Eukaryota-unknown)
summary.seqs(fasta=current, count=current)
count.groups(count=current)
#For a more robust comparison, removing samples with less than 2500 reads
remove.groups(fasta=current, count=current, groups=sample_x-sample_y-
sample_z)
count.groups(count=current)
```

### OTU-based analyses

#### *Calculate distances and clustering sequences to OTUs*

```
dist.seqs(fasta=current, cutoff=0.1)
cluster(column=current, count=current, cutoff=0.01)
```

#### *Create OTU99 table and classify OTUs*

```
make.shared(list=current, count=current, label=0.01)
classify.otu(list=current, label=0.01, taxonomy=current, count=current,
basis=sequence)
```

#### *Get OTU representative sequences (based on abundance)*

```
get.oturep(list=current, fasta=current, count=current, method=abundance)
```

#### *Calculate alpha diversity metrics*

```
summary.single(shared=current, subsample=T, calc=nseqs-coverage-sobs-ace-
chao-simpson-shannon-invsimpson-shannoneven-simpsoneven)
```

#### *Rarefying OTU table for statistical analyses*

```
sub.sample(shared=current)
```

### More-detailed OTU taxonomy using a freshwater database with SILVA

Additional Taxonomic assignments of the OTU99 representatives for both datasets were made using TaxAss (Rohwer et al., 2018 Msphere) with the FreshTrain 2020Jun15 and Silva SSU v138 databases

- Using a text editor header of the *.0.01.rep.fasta were modified with regular expressions to contain only the OTU IDs
- file was renamed to 'otus.fasta' as an input for TaxAss

```
cd TaxAss-master/tax-scripts/scripts
./RunSteps_quickie.sh otus FreshTrain15Jun2020silva138
silva_nr_v138_taxass 98 80 80 8
```

### Supplementary figures

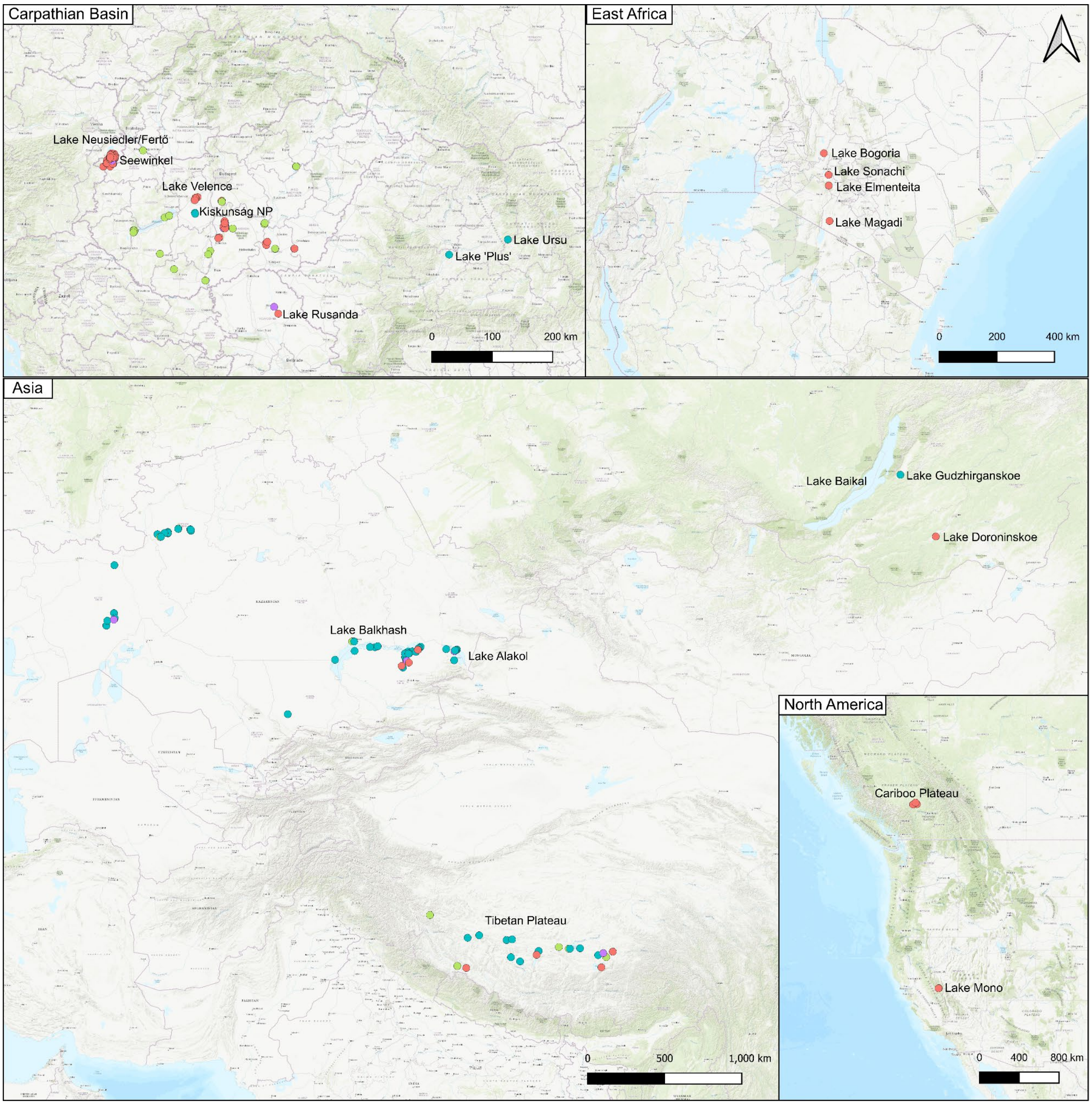


**Supplementary Fig. S1**. Detailed maps of the sampling sites. The sites are coloured according to ionic composition types. The maps were created using the QGIS software v3.28.6. The base map used is Esri.WorldTopoMap, sourced from Esri (https://server.arcgisonline.com/ArcGIS/rest/services/World_Topo_Map/MapServer/tile/{z}/{y}/{x})

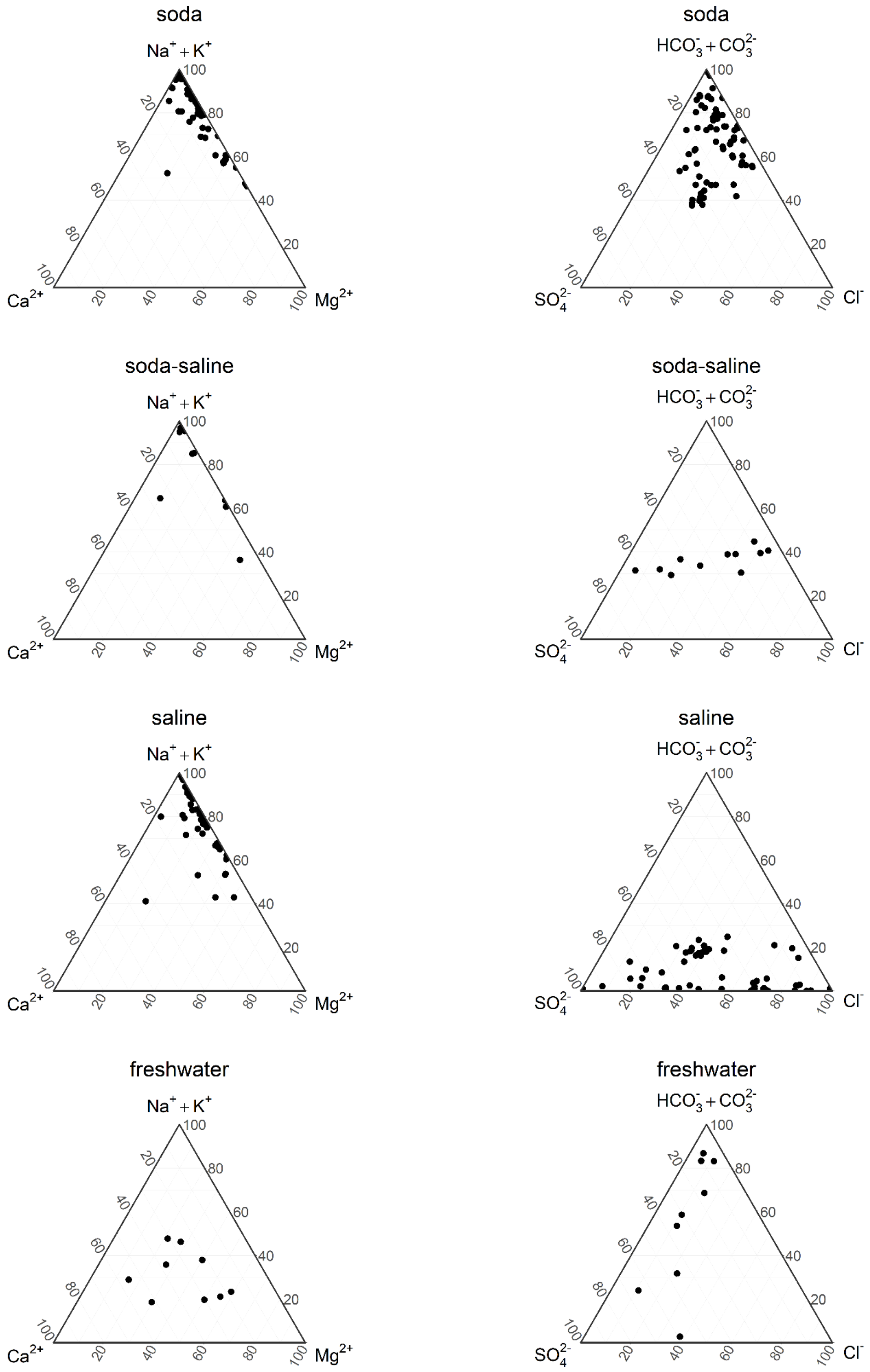


**Supplementary Fig. S2.** Ternary diagrams of the equivalent percentage (e%) contribution of dissolved ions to total ion content in the samples according to their ionic composition types.

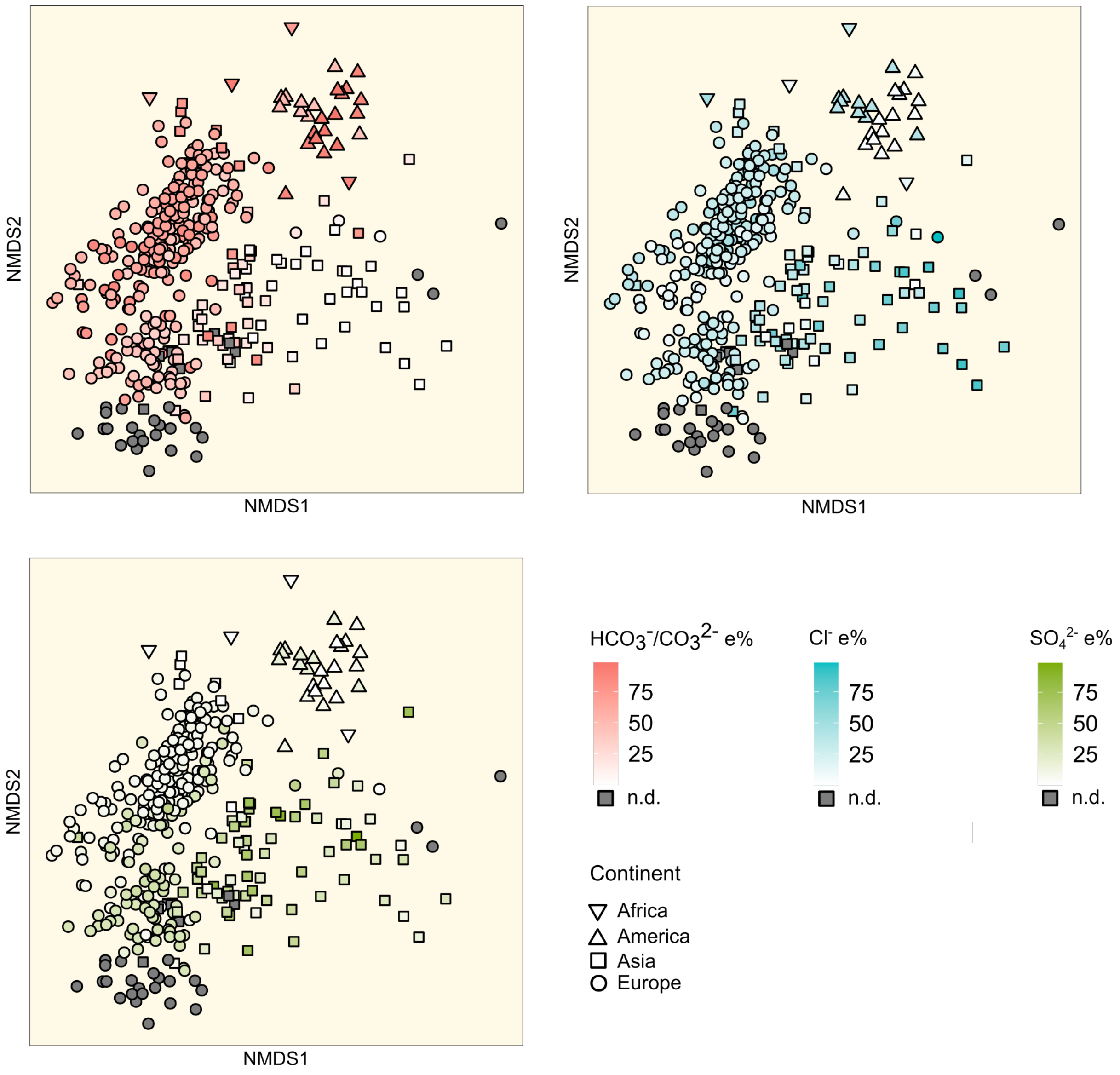


**Supplementary Fig. S3.** Comparison of planktonic freshwater and saline lake bacterial communities indicating the relative amount of different anions in the samples. NMDS ordination of bacterial OTUs defined at 99% similarity (stress 0.19) derived from the 16S rRNA V4 region sample set and rotated with salinity. Samples are coloured based on the major anion equivalent percentage (e%) of the sites, with different anions represented by distinct colours. Grey-coloured sites indicate a lack of data (n.d. - not determined) to calculate e% for every major anion.

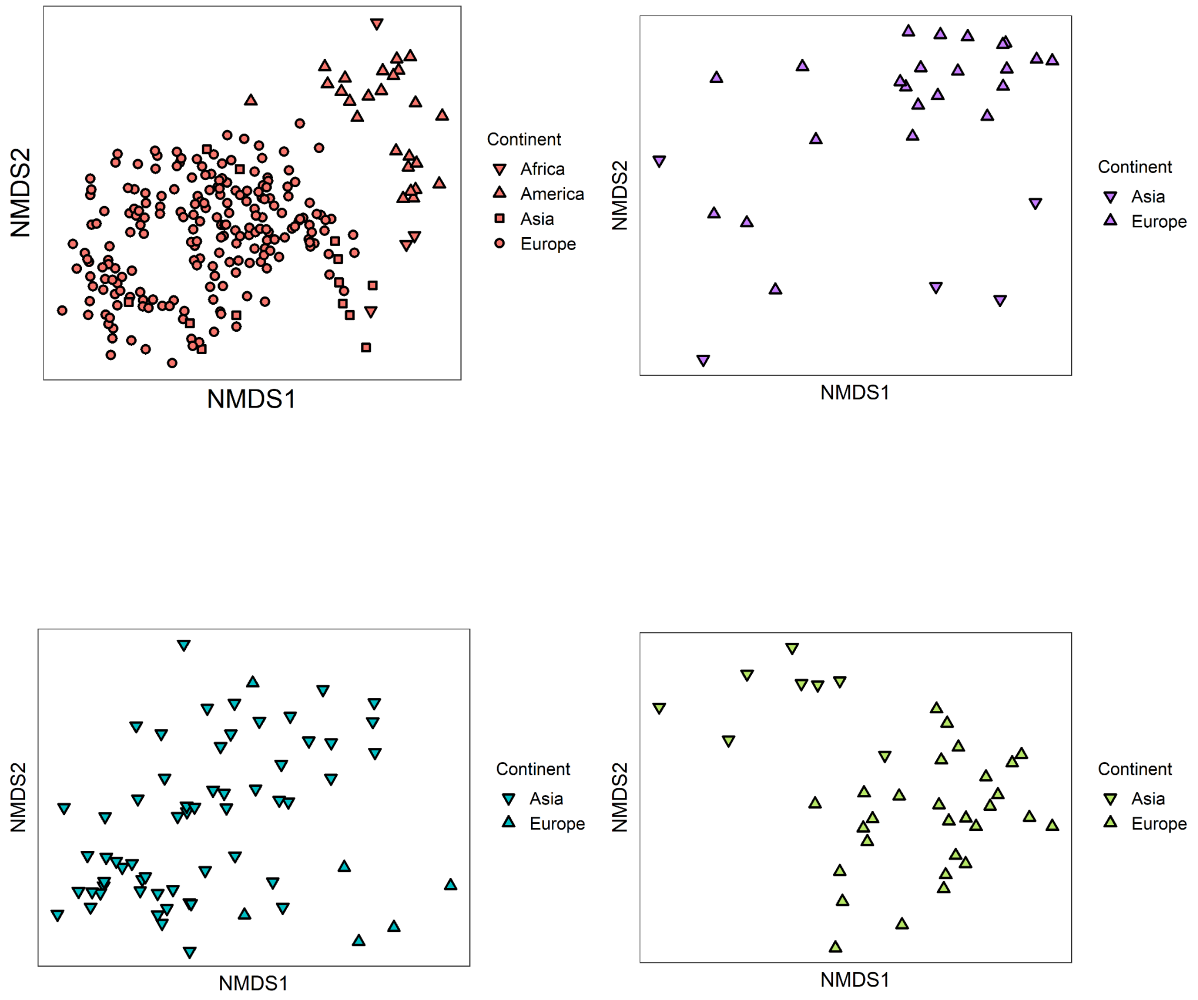


**Supplementary Fig. S4.** NMDS ordination of bacterial OTUs according to the sample's ionic composition types. Colour-coding in the figure corresponds to the colour scheme used throughout the manuscript.

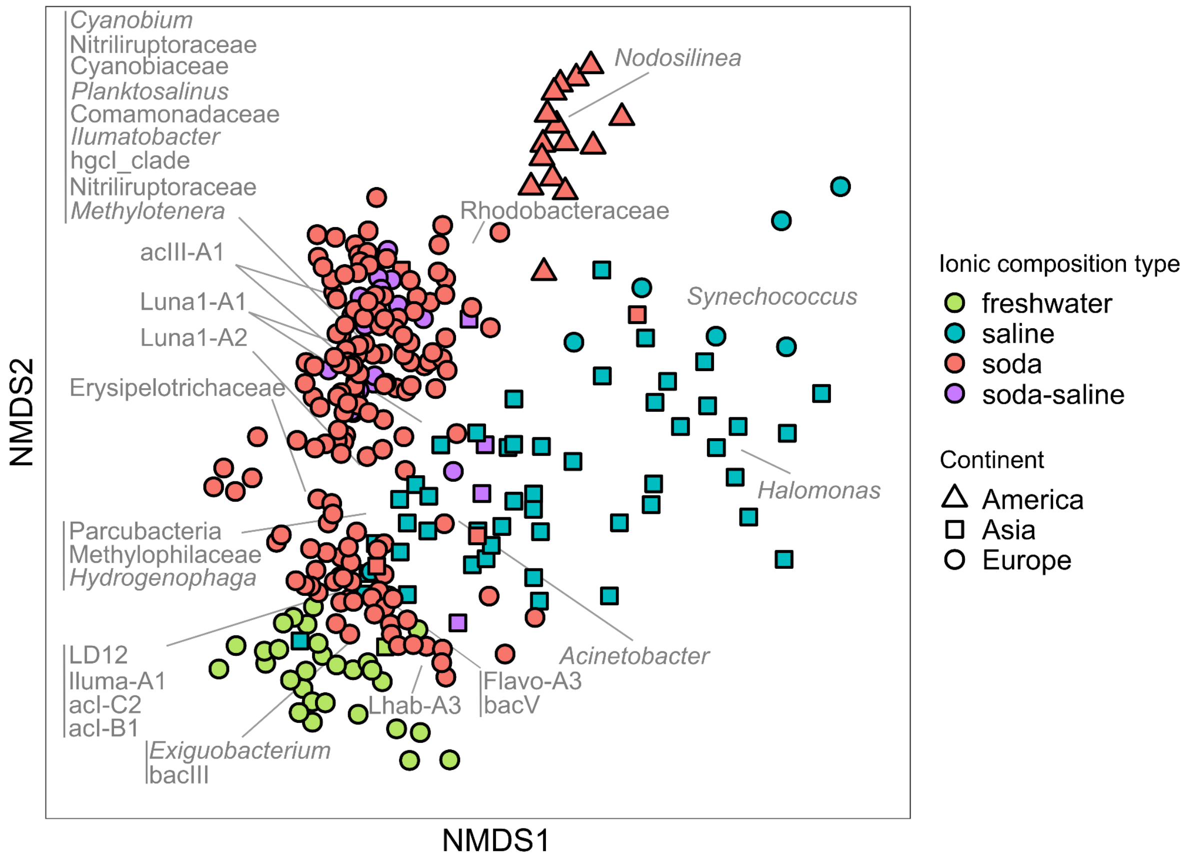


**Supplementary Fig. S5**. Comparison of planktonic freshwater and saline lake bacterial communities based on the V3-V4 region dataset. NMDS ordination of bacterial OTUs defined at 99% similarity (stress 0.17) derived from the 16S rRNA V3-V4 region sample set containing 282 samples from 78 sites and rotated with salinity. Based on the SIMPER analysis of water ion composition types (freshwater: green, saline: blue, soda: red, soda-saline: purple), the closest affiliated taxa of OTUs responsible for 25% dissimilarity among ion composition categories are shown in grey.

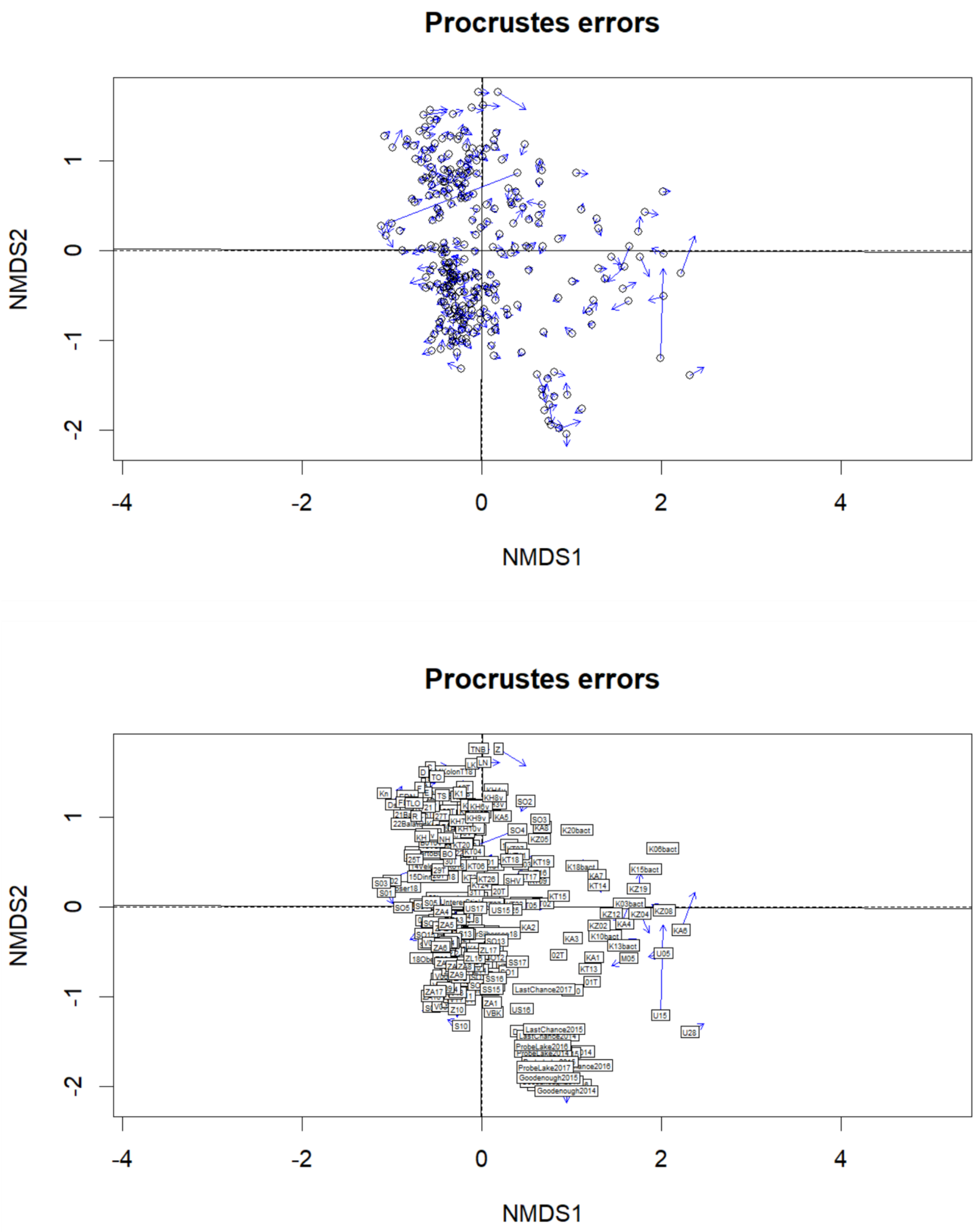


**Supplementary Fig. S6**. Dissimilarities in the ordination patterns between the V4 and V3-V4 datasets based on a procrustes analysis. To test the significance of the procrustes result 999 permutations were used with the 'protest' function. The result highlighted the similarity of the two ordinations, indicated by a good fit (Procrustes sum of squares: 0.02), and by a significant ($p < 0.01$) strong similarity ( $r = 0.989$).

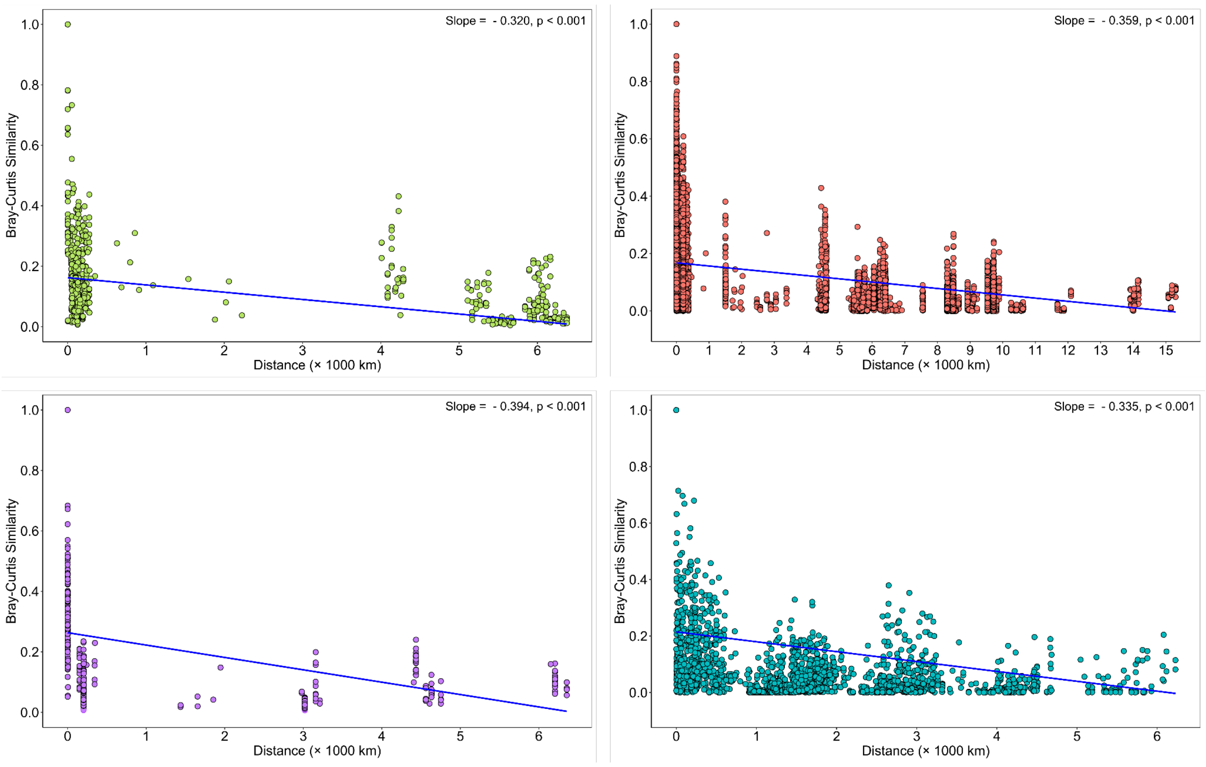


**Supplementary Fig. S7.** Distance-decay relationships between community similarity and geographic distance for inland surface water microbial communities. Blue lines represent regression calculated based on Bray-Curtis similarity values across all spatial distances. Colours represent the sampling site's ionic composition type. Green: freshwater, red: soda, purple: soda-saline, blue: saline sites. Despite accounting for all samples or individual ionic composition types, there was no significant correlation found between community similarity and geographic distance (Mantel $r_{all}$ = -0.328, $p > 0.05$).
