## Supplementary Table 1 for "A matter of salt: global assessment of the effect of salt ionic composition as a driver of aquatic bacterial diversity"

| Sample ID | Site name | Site description | Continent | Geographic region | Ionic composition type* |
|---|---|---|---|---|---|
| Bac-12S | Sonachi Crater Lake | soda lake | Africa | Great Rift Valley | soda |
| Bac-18E | Lake Elmenteita | soda lake | Africa | Great Rift Valley | soda |
| Bac-6M | Lake Magadi | soda lake | Africa | Great Rift Valley | soda |
| B1-W | Lake Bogoria | soda lake | Africa | Great Rift Valley | soda |
| B2-W | Lake Bogoria | soda lake | Africa | Great Rift Valley | soda |
| KZ02 | unknown | saline lake | Asia | North Kazakhstan | saline |
| K03bact | unknown | saline lake | Asia | North Kazakhstan | saline |
| KZ04 | Zharsor | saline lake | Asia | North Kazakhstan | saline |
| KZ05 | unknown | freshwater lake | Asia | North Kazakhstan | freshwater |
| K06bact | Kaindysor | saline lake | Asia | North Kazakhstan | saline |
| KZ08 | Asubastysor | saline lake | Asia | North Kazakhstan | saline |
| K10bact | unknown | saline lake | Asia | North Kazakhstan | saline |
| KZ12 | unknown | saline lake | Asia | North Kazakhstan | saline |
| K13bact | Big Aqsuat | saline lake | Asia | North Kazakhstan | saline |
| K15bact | unknown | saline lake | Asia | North Kazakhstan | saline |
| K16bact | Teniz | saline lake | Asia | North Kazakhstan | saline |
| K18bact | Little Aqsuat | saline lake | Asia | North Kazakhstan | saline |
| KZ19 | Kaiyndysor | saline lake | Asia | North Kazakhstan | saline |
| K20bact | Zharman | freshwater lake | Asia | North Kazakhstan | freshwater |
| KT01 | Lake Balkhash | Eastern Basin point 1 | Asia | Lake Balkhash Drainage Basin | saline |
| KT02 | Lake Balkhash | Eastern Basin point 2 | Asia | Lake Balkhash Drainage Basin | saline |
| KT03 | Lake Balkhash | Eastern Basin point 3 | Asia | Lake Balkhash Drainage Basin | saline |
| KT04 | Lake Balkhash | Eastern Basin point 4 | Asia | Lake Balkhash Drainage Basin | saline |
| KT05 | Lake Balkhash | Eastern Basin, unnamed lake, point 5 | Asia | Lake Balkhash Drainage Basin | saline |
| KT06 | Lake Balkhash | Eastern Basin point 6 | Asia | Lake Balkhash Drainage Basin | saline |
| KT07 | Lake Balkhash | Eastern Basin point 7 | Asia | Lake Balkhash Drainage Basin | saline |
| KT08 | Lake Balkhash | Eastern Basin point 8 | Asia | Lake Balkhash Drainage Basin | saline |
| KT09 | unknown | saline lake | Asia | Lake Balkhash Drainage Basin | saline |
| KT10 | unknown | saline lake | Asia | Lake Balkhash Drainage Basin | saline |
| KT11 | unknown | saline lake | Asia | Lake Balkhash Drainage Basin | saline |
| KT12 | unknown | soda-saline lake | Asia | Lake Balkhash Drainage Basin | soda-saline |
| KT13 | unknown | soda lake | Asia | Lake Balkhash Drainage Basin | soda |
| KT14 | unknown | saline lake | Asia | Lake Balkhash Drainage Basin | saline |
| KT15 | unknown | saline lake | Asia | Lake Balkhash Drainage Basin | saline |
| KT16 | Lake Balkhash | Eastern Basin point 9 | Asia | Lake Balkhash Drainage Basin | saline |
| KT17 | Lake Balkhash | Eastern Basin, unnamed lake, point 10 | Asia | Lake Balkhash Drainage Basin | saline |
| KT18 | Lake Balkhash | Eastern Basin, point 11 | Asia | Lake Balkhash Drainage Basin | soda |
| KT19 | unknown | saline lake | Asia | Lake Balkhash Drainage Basin | saline |
| KT20 | unknown | soda lake | Asia | Lake Balkhash Drainage Basin | soda |
| KT21 | Lake Sasykkol | saline lake | Asia | Lake Balkhash Drainage Basin | saline |
| KT22 | Lake Alakol | saline marshland | Asia | Lake Balkhash Drainage Basin | soda-saline |
| KT23 | Lake Alakol | saline lake | Asia | Lake Balkhash Drainage Basin | saline |
| KT24 | Lake Alakol | saline lake | Asia | Lake Balkhash Drainage Basin | saline |

| | | | | | | |
|---|---|---|---|---|---|---|
| KT25 | Lake Alakol | saline lake | | Asia | Lake Balkhash Drainage Basin | saline |
| KT26 | Lake Alakol | saline lake | | Asia | Lake Balkhash Drainage Basin | saline |
| KT27 | Lake Alakol | saline lake | | Asia | Lake Balkhash Drainage Basin | saline |
| 0S | Warm stream | saline lake | | Asia | Lake Balkhash | saline |
| 02S | Karabas | saline lake | | Asia | Lake Balkhash | saline |
| 04S | Korzhun | saline lake | | Asia | Lake Balkhash | saline |
| 06S | Kors | saline lake | | Asia | Lake Balkhash | saline |
| 08S | Municipal beach | saline lake | | Asia | Lake Balkhash | saline |
| 09S | Narine | freshwater lake | | Asia | Lake Balkhash | freshwater |
| 11S | Saline lake | saline lake | | Asia | Lake Balkhash | saline |
| SHV | Old Shalkar Lake | soda-saline lake | | Asia | West Kazakhstan, Aktobe region | soda-saline |
| KA1 | unknown | saline lake | | Asia | West Kazakhstan, Aktobe region | saline |
| KA2 | Lakalu | saline lake | | Asia | West Kazakhstan, Aktobe region | saline |
| KA3 | Lakalu | saline lake | | Asia | West Kazakhstan, Aktobe region | saline |
| KA4 | Kopasor | saline lake | | Asia | West Kazakhstan, Aktobe region | saline |
| KA5 | Shalkar Lake | soda-saline lake | | Asia | West Kazakhstan, Aktobe region | soda-saline |
| KA6 | saline lake | saline lake | | Asia | West Kazakhstan, Aktobe region | saline |
| KA7 | Belkopa | saline lake | | Asia | West Kazakhstan, Aktobe region | saline |
| KA8 | unknown | saline lake | | Asia | South Kazakhstan, Zhambyl region | saline |
| Gudzh_water | Lake Gudzhirganskoe | saline lake | | Asia | Barguzin Valley, Lake Baikal | saline |
| NB30 | Lake Doroninskoe | soda lake | | Asia | Transbaikalia | soda |
| NB31 | Lake Doroninskoe | soda lake | | Asia | Transbaikalia | soda |
| NB32 | Lake Doroninskoe | soda lake | | Asia | Transbaikalia | soda |
| NB41 | Lake Doroninskoe | soda lake | | Asia | Transbaikalia | soda |
| NB42 | Lake Doroninskoe | soda lake | | Asia | Transbaikalia | soda |
| NB43 | Lake Doroninskoe | soda lake | | Asia | Transbaikalia | soda |
| Bangong Co | Bangong Co | freshwater lake | | Asia | Tibetan Plateau | freshwater |
| Bong Co | Bong Co | freshwater lake | | Asia | Tibetan Plateau | freshwater |
| Co Ngoin (1) | Co Ngoin | freshwater lake | | Asia | Tibetan Plateau | freshwater |
| Mapam Yumco | Mapam Yumco | freshwater lake | | Asia | Tibetan Plateau | freshwater |
| Urru Co | Urru Co | freshwater lake | | Asia | Tibetan Plateau | freshwater |
| Bam Co | Bam Co | saline lake | | Asia | Tibetan Plateau | saline |
| Bero Zeco | Bero Zeco | saline lake | | Asia | Tibetan Plateau | saline |
| Co Ngoin (2) | Co Ngoin | soda lake | | Asia | Tibetan Plateau | soda |
| Dawa Co | Dawa Co | saline lake | | Asia | Tibetan Plateau | saline |
| Tangra Yumco | Tangra Yumco | soda lake | | Asia | Tibetan Plateau | soda |
| Dong Co | Dong Co | saline lake | | Asia | Tibetan Plateau | saline |
| Kunggyu | Kunggyu | soda lake | | Asia | Tibetan Plateau | soda |
| Nam Co | Nam Co | soda lake | | Asia | Tibetan Plateau | soda |
| Pung Co | Pung Co | soda-saline lake | | Asia | Tibetan Plateau | soda-saline |
| Selin Co | Selin Co | saline lake | | Asia | Tibetan Plateau | saline |
| Zhari Namco | Zhari Namco | saline lake | | Asia | Tibetan Plateau | saline |
| Zhaxi Co | Zhaxi Co | saline lake | | Asia | Tibetan Plateau | saline |
| Bangkog Co | Bangkog Co | saline lake | | Asia | Tibetan Plateau | saline |
| Dangqiong Co | Dangqiong Co | saline lake | | Asia | Tibetan Plateau | saline |

| | | | | | |
|---|---|---|---|---|---|
| Nyer Co (Nieer Co) | Nyer Co (Nieer Co) | saline lake | Asia | Tibetan Plateau | saline |
| BU1 | Budos-szek | soda pan | Europe | Carpathian Basin | soda |
| SO1 | Sos-er | soda pan | Europe | Carpathian Basin | soda |
| ZA1 | Zab-szek | soda pan | Europe | Carpathian Basin | soda |
| SO2 | Sos-er | soda pan | Europe | Carpathian Basin | soda |
| SO3 | Sos-er | soda pan | Europe | Carpathian Basin | soda |
| SO4 | Sos-er | soda pan | Europe | Carpathian Basin | soda |
| SO5 | Sos-er | soda pan | Europe | Carpathian Basin | soda |
| SO6 | Sos-er | soda pan | Europe | Carpathian Basin | soda |
| SO9 | Sos-er | soda pan | Europe | Carpathian Basin | soda |
| SO11 | Sos-er | soda pan | Europe | Carpathian Basin | soda |
| SO12 | Sos-er | soda pan | Europe | Carpathian Basin | soda |
| SO13 | Sos-er | soda pan | Europe | Carpathian Basin | soda |
| SO14 | Sos-er | soda pan | Europe | Carpathian Basin | soda |
| SO15 | Sos-er | soda pan | Europe | Carpathian Basin | soda |
| SO16 | Sos-er | soda pan | Europe | Carpathian Basin | soda |
| VBK | pan no. 60 | soda pan | Europe | Carpathian Basin | soda |
| ZA2 | Zab-szek | soda pan | Europe | Carpathian Basin | soda |
| ZA3 | Zab-szek | soda pan | Europe | Carpathian Basin | soda |
| ZA4 | Zab-szek | soda pan | Europe | Carpathian Basin | soda |
| ZA5 | Zab-szek | soda pan | Europe | Carpathian Basin | soda |
| ZA6 | Zab-szek | soda pan | Europe | Carpathian Basin | soda |
| ZA7 | Zab-szek | soda pan | Europe | Carpathian Basin | soda |
| ZA8 | Zab-szek | soda pan | Europe | Carpathian Basin | soda |
| ZA9 | Zab-szek | soda pan | Europe | Carpathian Basin | soda |
| ZA10 | Zab-szek | soda pan | Europe | Carpathian Basin | soda |
| ZA11 | Zab-szek | soda pan | Europe | Carpathian Basin | soda |
| ZA12 | Zab-szek | soda pan | Europe | Carpathian Basin | soda |
| ZA13 | Zab-szek | soda pan | Europe | Carpathian Basin | soda |
| ZA14 | Zab-szek | soda pan | Europe | Carpathian Basin | soda |
| ZA15 | Zab-szek | soda pan | Europe | Carpathian Basin | soda |
| ZA16 | Zab-szek | soda pan | Europe | Carpathian Basin | soda |
| ZA17 | Zab-szek | soda pan | Europe | Carpathian Basin | soda |
| M05 | Lacul Ursu | saline lake | Europe | Carpathian Basin | saline |
| U05 | Lacul 'Plus' | saline lake | Europe | Carpathian Basin | saline |
| U15 | Lacul 'Plus' | saline lake | Europe | Carpathian Basin | saline |
| U28 | Lacul 'Plus' | saline lake | Europe | Carpathian Basin | saline |
| B01w | Lake Ferto/Neusiedl | soda lake, open water | Europe | Carpathian Basin | soda |
| B02w | Lake Ferto/Neusiedl | soda lake, open water | Europe | Carpathian Basin | soda |
| B03w | Lake Ferto/Neusiedl | soda lake, open water | Europe | Carpathian Basin | soda |
| B04w | Lake Ferto/Neusiedl | soda lake, open water | Europe | Carpathian Basin | soda |
| B05w | Lake Ferto/Neusiedl | soda lake, open water | Europe | Carpathian Basin | soda |
| B06w | Lake Ferto/Neusiedl | soda lake, open water | Europe | Carpathian Basin | soda |
| B07w | Lake Ferto/Neusiedl | soda lake, open water | Europe | Carpathian Basin | soda |
| B08w | Lake Ferto/Neusiedl | soda lake, open water | Europe | Carpathian Basin | soda |

| | | | | | |
|---|---|---|---|---|---|
| B09w | Lake Ferto/Neusiedl | soda lake, open water | Europe | Carpathian Basin | soda |
| B010w | Lake Ferto/Neusiedl | soda lake, open water | Europe | Carpathian Basin | soda |
| B011w | Lake Ferto/Neusiedl | soda lake, open water | Europe | Carpathian Basin | soda |
| KH1w | Lake Ferto/Neusiedl | soda lake, Kis-Herlakni inner pond | Europe | Carpathian Basin | soda |
| KH2w | Lake Ferto/Neusiedl | soda lake, Kis-Herlakni inner pond | Europe | Carpathian Basin | soda |
| KH3w | Lake Ferto/Neusiedl | soda lake, Kis-Herlakni inner pond | Europe | Carpathian Basin | soda |
| KH4w | Lake Ferto/Neusiedl | soda lake, Kis-Herlakni inner pond | Europe | Carpathian Basin | soda |
| KH5w | Lake Ferto/Neusiedl | soda lake, Kis-Herlakni inner pond | Europe | Carpathian Basin | soda |
| KH6w | Lake Ferto/Neusiedl | soda lake, Kis-Herlakni inner pond | Europe | Carpathian Basin | soda |
| KH7w | Lake Ferto/Neusiedl | soda lake, Kis-Herlakni inner pond | Europe | Carpathian Basin | soda |
| KH8w | Lake Ferto/Neusiedl | soda lake, Kis-Herlakni inner pond | Europe | Carpathian Basin | soda |
| KH9w | Lake Ferto/Neusiedl | soda lake, Kis-Herlakni inner pond | Europe | Carpathian Basin | soda |
| KH10w | Lake Ferto/Neusiedl | soda lake, Kis-Herlakni inner pond | Europe | Carpathian Basin | soda |
| KH11w | Lake Ferto/Neusiedl | soda lake, Kis-Herlakni inner pond | Europe | Carpathian Basin | soda |
| B01 | Boddi-szek | soda-saline pan | Europe | Carpathian Basin | soda-saline |
| B02 | Boddi-szek | soda-saline pan | Europe | Carpathian Basin | soda-saline |
| B03 | Boddi-szek | soda-saline pan | Europe | Carpathian Basin | soda-saline |
| B04 | Boddi-szek | soda-saline pan | Europe | Carpathian Basin | soda-saline |
| B05 | Boddi-szek | soda-saline pan | Europe | Carpathian Basin | soda-saline |
| B06 | Boddi-szek | soda-saline pan | Europe | Carpathian Basin | soda-saline |
| B07 | Boddi-szek | soda-saline pan | Europe | Carpathian Basin | soda-saline |
| B08 | Boddi-szek | soda-saline pan | Europe | Carpathian Basin | soda-saline |
| B09 | Boddi-szek | soda-saline pan | Europe | Carpathian Basin | soda-saline |
| B10 | Boddi-szek | soda-saline pan | Europe | Carpathian Basin | soda-saline |
| B11 | Boddi-szek | soda-saline pan | Europe | Carpathian Basin | soda-saline |
| B12 | Boddi-szek | soda-saline pan | Europe | Carpathian Basin | soda-saline |
| B13 | Boddi-szek | soda-saline pan | Europe | Carpathian Basin | soda-saline |
| B14 | Boddi-szek | soda-saline pan | Europe | Carpathian Basin | soda-saline |
| K01 | Kelemen-szek | soda pan | Europe | Carpathian Basin | soda |
| K02 | Kelemen-szek | soda pan | Europe | Carpathian Basin | soda |
| K03 | Kelemen-szek | soda pan | Europe | Carpathian Basin | soda |
| K04 | Kelemen-szek | soda pan | Europe | Carpathian Basin | soda |
| K05 | Kelemen-szek | soda pan | Europe | Carpathian Basin | soda |
| K06 | Kelemen-szek | soda pan | Europe | Carpathian Basin | soda |
| K09 | Kelemen-szek | soda pan | Europe | Carpathian Basin | soda |
| K12 | Kelemen-szek | soda pan | Europe | Carpathian Basin | soda |
| K14 | Kelemen-szek | soda pan | Europe | Carpathian Basin | soda |
| S01 | Sos-er | soda pan | Europe | Carpathian Basin | soda |
| S02 | Sos-er | soda pan | Europe | Carpathian Basin | soda |
| S03 | Sos-er | soda pan | Europe | Carpathian Basin | soda |
| S04 | Sos-er | soda pan | Europe | Carpathian Basin | soda |
| S05 | Sos-er | soda pan | Europe | Carpathian Basin | soda |
| S06 | Sos-er | soda pan | Europe | Carpathian Basin | soda |
| S07 | Sos-er | soda pan | Europe | Carpathian Basin | soda |
| S08 | Sos-er | soda pan | Europe | Carpathian Basin | soda |

| | | | | | |
|---|---|---|---|---|---|
| S09 | Sos-er | soda pan | Europe | Carpathian Basin | soda |
| S10 | Sos-er | soda pan | Europe | Carpathian Basin | soda |
| S11 | Sos-er | soda pan | Europe | Carpathian Basin | soda |
| S12 | Sos-er | soda pan | Europe | Carpathian Basin | soda |
| S13 | Sos-er | soda pan | Europe | Carpathian Basin | soda |
| S14 | Sos-er | soda pan | Europe | Carpathian Basin | soda |
| V01 | pan no. 60 | soda pan | Europe | Carpathian Basin | soda |
| V02 | pan no. 60 | soda pan | Europe | Carpathian Basin | soda |
| V03 | pan no. 60 | soda pan | Europe | Carpathian Basin | soda |
| V04 | pan no. 60 | soda pan | Europe | Carpathian Basin | soda |
| V05 | pan no. 60 | soda pan | Europe | Carpathian Basin | soda |
| V06 | pan no. 60 | soda pan | Europe | Carpathian Basin | soda |
| V07 | pan no. 60 | soda pan | Europe | Carpathian Basin | soda |
| V08 | pan no. 60 | soda pan | Europe | Carpathian Basin | soda |
| V09 | pan no. 60 | soda pan | Europe | Carpathian Basin | soda |
| V10 | pan no. 60 | soda pan | Europe | Carpathian Basin | soda |
| V11 | pan no. 60 | soda pan | Europe | Carpathian Basin | soda |
| V12 | pan no. 60 | soda pan | Europe | Carpathian Basin | soda |
| V13 | pan no. 60 | soda pan | Europe | Carpathian Basin | soda |
| V14 | pan no. 60 | soda pan | Europe | Carpathian Basin | soda |
| Z01 | Zab-szek | soda pan | Europe | Carpathian Basin | soda |
| Z02 | Zab-szek | soda pan | Europe | Carpathian Basin | soda |
| Z03 | Zab-szek | soda pan | Europe | Carpathian Basin | soda |
| Z04 | Zab-szek | soda pan | Europe | Carpathian Basin | soda |
| Z05 | Zab-szek | soda pan | Europe | Carpathian Basin | soda |
| Z06 | Zab-szek | soda pan | Europe | Carpathian Basin | soda |
| Z07 | Zab-szek | soda pan | Europe | Carpathian Basin | soda |
| Z08 | Zab-szek | soda pan | Europe | Carpathian Basin | soda |
| Z09 | Zab-szek | soda pan | Europe | Carpathian Basin | soda |
| Z10 | Zab-szek | soda pan | Europe | Carpathian Basin | soda |
| Z12 | Zab-szek | soda pan | Europe | Carpathian Basin | soda |
| Z14 | Zab-szek | soda pan | Europe | Carpathian Basin | soda |
| BV | Lake Balaton | freshwater lake | Europe | Carpathian Basin | freshwater |
| K0 | Lake Kolon | freshwater lake | Europe | Carpathian Basin | freshwater |
| K01 | Lake Kolon | freshwater lake | Europe | Carpathian Basin | freshwater |
| A | Oxbow Tiszaalpar | freshwater lake, open water | Europe | Carpathian Basin | freshwater |
| B | Oxbow Tiszaalpar | freshwater lake, site with submerged macrophytes | Europe | Carpathian Basin | freshwater |
| B0 | Lake Ferto/Neusiedl | soda lake, open water | Europe | Carpathian Basin | soda |
| C | Oxbow Lakiteleki | freshwater lake, site with submerged macrophytes | Europe | Carpathian Basin | freshwater |
| D | Oxbow Lakiteleki | freshwater lake, open water | Europe | Carpathian Basin | freshwater |
| Ds | Reservoir Deseda | freshwater lake, open water | Europe | Carpathian Basin | freshwater |
| E | Oxbow Martely | freshwater lake, open water | Europe | Carpathian Basin | freshwater |
| F | Oxbow Martely | freshwater lake, site with submerged macrophytes | Europe | Carpathian Basin | freshwater |
| FDN | Oxbow Fadd-Dombori | freshwater lake, site with emergent macrophytes | Europe | Carpathian Basin | freshwater |
| FDO | Oxbow Fadd-Dombori | freshwater lake, open water | Europe | Carpathian Basin | freshwater |

| | | | | | |
|---|---|---|---|---|---|
| Ff | Lake Balaton | freshwater lake, open water, Siofok Basin | Europe | Carpathian Basin | freshwater |
| K | Lake Balaton | freshwater lake, open water, Keszthely Basin | Europe | Carpathian Basin | freshwater |
| KH | Lake Ferto/Neusiedl | soda lake, Kis-Herlakni inner pond, site with submerged macrophytes | Europe | Carpathian Basin | soda |
| Kn | Lake Kovacsszenaja | freshwater lake, open water | Europe | Carpathian Basin | freshwater |
| LK | Lake Morotva | freshwater lake, site with submerged macrophytes | Europe | Carpathian Basin | freshwater |
| LN | Lake Morotva | freshwater lake, site with emergent macrophytes | Europe | Carpathian Basin | freshwater |
| NH | Lake Ferto/Neusiedl | soda lake, Nagy-Herlakni inner pond, site with submerged macrophytes | Europe | Carpathian Basin | soda |
| R | Oxbow Riha | freshwater lake, open water | Europe | Carpathian Basin | freshwater |
| TLO | Oxbow Tolnai | freshwater lake, open water | Europe | Carpathian Basin | freshwater |
| TNB | Lake Tisza | freshwater lake, site with emergent macrophytes | Europe | Carpathian Basin | freshwater |
| TO | Lake Tisza | freshwater lake, open water | Europe | Carpathian Basin | freshwater |
| TS | Lake Tisza | freshwater lake, site with submerged macrophytes | Europe | Carpathian Basin | freshwater |
| Z | Lake Balaton | freshwater lake, inflow of Zala River, site with submerged macrophytes | Europe | Carpathian Basin | freshwater |
| SS15 | Silberlacke (Sudlicher Silbersee) | soda pan | Europe | Carpathian Basin | soda |
| SS16 | Silberlacke (Sudlicher Silbersee) | soda pan | Europe | Carpathian Basin | soda |
| SS17 | Silberlacke (Sudlicher Silbersee) | soda pan | Europe | Carpathian Basin | soda |
| US15 | Unterer Stinkersee | soda pan | Europe | Carpathian Basin | soda |
| US16 | Unterer Stinkersee | soda pan | Europe | Carpathian Basin | soda |
| US17 | Unterer Stinkersee | soda pan | Europe | Carpathian Basin | soda |
| ZL15 | Zicklacke | soda pan | Europe | Carpathian Basin | soda |
| ZL16 | Zicklacke | soda pan | Europe | Carpathian Basin | soda |
| ZL17 | Zicklacke | soda pan | Europe | Carpathian Basin | soda |
| Sp17_01 | Albersee | soda pan | Europe | Carpathian Basin | soda |
| Sp17_02 | Auerlacke | soda pan | Europe | Carpathian Basin | soda |
| Sp17_04 | Birnbaumlacke | soda pan | Europe | Carpathian Basin | soda |
| Sp17_05 | Borsodi-dulo | soda-saline pan | Europe | Carpathian Basin | soda-saline |
| Sp17_07 | Grosse Neubruchlacke | soda pan | Europe | Carpathian Basin | soda |
| Sp17_09 | Krautingsee | soda pan | Europe | Carpathian Basin | soda |
| Sp17_10 | Kuhbrunnlacke | soda pan | Europe | Carpathian Basin | soda |
| Sp17_11 | Lange Lacke | soda pan | Europe | Carpathian Basin | soda |
| Sp17_12 | Martenhofenlacke | soda pan | Europe | Carpathian Basin | soda |
| Sp17_13 | Mittlerer Stinkersee | soda pan | Europe | Carpathian Basin | soda |
| Sp17_14 | unknown | soda pan, beside Warmsee | Europe | Carpathian Basin | soda |
| Sp17_15 | Nyeki-szallas | soda pan | Europe | Carpathian Basin | soda |
| Sp17_16 | Obere Hollacke | soda pan | Europe | Carpathian Basin | soda |
| Sp17_17 | Oberer Stinkersee | soda pan | Europe | Carpathian Basin | soda |
| Sp17_18 | Ochsenbrunnlacke | soda pan | Europe | Carpathian Basin | soda |
| Sp17_19 | Ostliche Fuchslochlacke | soda pan | Europe | Carpathian Basin | soda |
| Sp17_20 | Ostliche Worthenlacke | soda pan | Europe | Carpathian Basin | soda |
| Sp17_21 | Runde Lacke | soda pan | Europe | Carpathian Basin | soda |
| Sp17_22 | Sechsmahdlacke | soda pan | Europe | Carpathian Basin | soda |
| Sp17_23 | Standlacke | soda pan | Europe | Carpathian Basin | soda |
| Sp17_24 | Sudlicher Silbersee | soda pan | Europe | Carpathian Basin | soda |
| Sp17_27 | Westliche Fuschslochlacke | soda pan | Europe | Carpathian Basin | soda |
| Sp17_28 | Westliche Wörthenlacke | soda pan | Europe | Carpathian Basin | soda |

| | | | | | |
|---|---|---|---|---|---|
| Sp17_29 | Zicklacke | soda pan | Europe | Carpathian Basin | soda |
| Sp17_31 | Borsodi-dulo | soda-saline pan, inflow | Europe | Carpathian Basin | soda-saline |
| Sp17_Neu2 | Neufeldlacke | soda pan | Europe | Carpathian Basin | soda |
| Sp18_01 | Albersee | soda pan | Europe | Carpathian Basin | soda |
| Sp18_04 | Birnbaumlacke | soda pan | Europe | Carpathian Basin | soda |
| Sp18_05 | Borsodi-dulo | soda-saline pan | Europe | Carpathian Basin | soda-saline |
| Sp18_07 | Grosse Neubruchlacke | soda pan | Europe | Carpathian Basin | soda |
| Sp18_08 | Kirchsee | soda pan | Europe | Carpathian Basin | soda |
| Sp18_10 | Kuhbrunnlacke | soda pan | Europe | Carpathian Basin | soda |
| Sp18_11 | Lange Lacke | soda pan | Europe | Carpathian Basin | soda |
| Sp18_12 | Martenhofenlacke | soda pan | Europe | Carpathian Basin | soda |
| Sp18_13 | Mittlerer Stinkersee | soda pan | Europe | Carpathian Basin | soda |
| Sp18_15 | Nyeki-szallas | soda pan | Europe | Carpathian Basin | soda |
| Sp18_16 | Obere Hollacke | soda pan | Europe | Carpathian Basin | soda |
| Sp18_17 | Oberer Stinkersee | soda pan | Europe | Carpathian Basin | soda |
| Sp18_18 | Ochsenbrunnlacke | soda pan | Europe | Carpathian Basin | soda |
| Sp18_19 | Ostliche Fuchslochlacke | soda pan | Europe | Carpathian Basin | soda |
| Sp18_1_d | Auerlacke | soda pan | Europe | Carpathian Basin | soda |
| Sp18_20 | Ostliche Worthenlacke | soda pan | Europe | Carpathian Basin | soda |
| Sp18_21 | Runde Lacke | soda pan | Europe | Carpathian Basin | soda |
| Sp18_22 | Sechsmahdlacke | soda pan | Europe | Carpathian Basin | soda |
| Sp18_23 | Standlacke | soda pan | Europe | Carpathian Basin | soda |
| Sp18_24 | Sudlicher Silbersee | soda pan | Europe | Carpathian Basin | soda |
| Sp18_27 | Westliche Fuschslochlacke | soda pan | Europe | Carpathian Basin | soda |
| Sp18_28 | Westliche Worthenlacke | soda pan | Europe | Carpathian Basin | soda |
| Sp18_29 | Zicklacke | soda pan | Europe | Carpathian Basin | soda |
| Sp18_30 | Apetloner Meierhoflacke | soda-saline pan | Europe | Carpathian Basin | soda-saline |
| Sp18_31 | Herrnsee | soda pan | Europe | Carpathian Basin | soda |
| Sp18_9 | Krautingsee | soda pan | Europe | Carpathian Basin | soda |
| Sp18_Neu2 | Neufeldlacke | soda pan | Europe | Carpathian Basin | soda |
| 01Boddi | Boddi-szek | soda pan | Europe | Carpathian Basin | soda-saline |
| 02Soser | Sos-er | soda pan | Europe | Carpathian Basin | soda |
| 03KolonNy | Lake Kolon | freshwater lake, open water | Europe | Carpathian Basin | freshwater |
| 04KolonT18 | Lake Kolon | freshwater lake, site with *Nymphaea* | Europe | Carpathian Basin | freshwater |
| 05Szelid18 | Lake Szelid | soda lake | Europe | Carpathian Basin | soda |
| 06Rusanda18 | Ruszanda | soda lake | Europe | Carpathian Basin | soda |
| 07SlanoKopovo18 | Soskopo | soda pan | Europe | Carpathian Basin | soda-saline |
| 08VBK18 | pan no. 60 | soda pan | Europe | Carpathian Basin | soda |
| 09Kelemen18 | Kelemen-szek | soda pan | Europe | Carpathian Basin | soda |
| 10Zab18 | Zab-szek | soda pan | Europe | Carpathian Basin | soda |
| 12KH18 | Lake Ferto/Neusiedl | soda lake, Kis-Herlakni inner pond | Europe | Carpathian Basin | soda |
| 13FertoB018 | Lake Ferto/Neusiedl | soda lake, open water | Europe | Carpathian Basin | soda |
| 14VelenceiKo18 | Lake Velence | soda lake, Eastern part | Europe | Carpathian Basin | soda |
| 15Dinnyesi18 | Dinnyesi-ferto | soda lake | Europe | Carpathian Basin | soda |
| 16VelenceiNy18 | Lake Velence | soda lake, Western part | Europe | Carpathian Basin | soda |

| | | | | | |
|---|---|---|---|---|---|
| 17Zicklacke18 | Zicklacke | soda pan | Europe | Carpathian Basin | soda |
| 18ObererStinker18 | Oberer Stinkersee | soda pan | Europe | Carpathian Basin | soda |
| 19SudlicherSilbersee18 | Sudlicher Silbersee | soda pan | Europe | Carpathian Basin | soda |
| 20LangeLacke18 | Lange Lacke | soda pan | Europe | Carpathian Basin | soda |
| 21BalatonTihany18 | Lake Balaton | freshwater lake, open water, Siofok Basin | Europe | Carpathian Basin | freshwater |
| 22BalatonKeszthely18 | Lake Balaton | freshwater lake, open water, Keszthely Basin | Europe | Carpathian Basin | freshwater |
| 23UntererStinker18 | Unterer Stinkersee | soda pan | Europe | Carpathian Basin | soda |
| 01T | Sarkany-to | saline lake | Europe | Carpathian Basin | saline |
| 02T | Sosto | saline lake | Europe | Carpathian Basin | saline |
| 03T | Budos-szek | soda pan, near Pusztaszer | Europe | Carpathian Basin | soda |
| 04T | Vesszos-szek | near Pusztaszer | Europe | Carpathian Basin | soda |
| 05T | Baks, Donger | soda lake | Europe | Carpathian Basin | soda |
| 06T | Sos-er | soda pan | Europe | Carpathian Basin | soda |
| 07T | Boddi-szek | soda-saline pan, Southern part | Europe | Carpathian Basin | soda-saline |
| 08T | Boddi-szek | soda-saline pan, Northern part | Europe | Carpathian Basin | soda-saline |
| 09T | Kelemen-szek | soda pan, Southern part | Europe | Carpathian Basin | soda |
| 10T | Kelemen-szek | soda pan, Northern part | Europe | Carpathian Basin | soda |
| 11T | Budos-szek | soda pan, near Szabadszallas | Europe | Carpathian Basin | soda |
| 12T | Zab-szek | soda pan, Northern part | Europe | Carpathian Basin | soda |
| 13T | Zab-szek | soda pan, Southern part | Europe | Carpathian Basin | soda |
| 14T | Lake Feher | soda lake, Western part | Europe | Carpathian Basin | soda |
| 15T | Lake Feher | soda lake, Eastern part | Europe | Carpathian Basin | soda |
| 16T | Also-Szunyogi ret | artificial pan, site 1 | Europe | Carpathian Basin | freshwater |
| 17T | Also-Szunyogi ret | artificial pan, site 2 | Europe | Carpathian Basin | soda |
| 18T | Also-Szunyogi ret | artificial pan, site 3 | Europe | Carpathian Basin | freshwater |
| 19T | Nyeki szallas | soda pan | Europe | Carpathian Basin | soda |
| 20T | Borsodi-dulo | soda-saline pan | Europe | Carpathian Basin | soda-saline |
| 21T | Lake Ferto/Neusiedl | soda lake, Kis-Herlakni inner pond | Europe | Carpathian Basin | soda |
| 22T | Lake Ferto/Neusiedl | soda lake, open water, Fertorakos Bay | Europe | Carpathian Basin | soda |
| 23T | Lake Ferto/Neusiedl | soda lake, open water | Europe | Carpathian Basin | soda |
| 24T | Lake Ferto/Neusiedl | soda lake, Hidegsegi inner pond | Europe | Carpathian Basin | soda |
| 25T | Lake Ferto/Neusiedl | soda lake, open water, Madarvarta Bay | Europe | Carpathian Basin | soda |
| 26T | Lake Szelid | soda lake, Katona steg | Europe | Carpathian Basin | soda |
| 27T | Lake Szelid | soda lake, To Szallo | Europe | Carpathian Basin | soda |
| 28T | Lake Velence | soda lake, Hosszu-tisztas | Europe | Carpathian Basin | soda |
| 29T | Lake Velence | soda lake, Langi-tisztas | Europe | Carpathian Basin | soda |
| 30T | Lake Velence | soda lake, Felsoeri-tisztasok | Europe | Carpathian Basin | soda |
| 31T | Dinnyesi-ferto | soda lake | Europe | Carpathian Basin | soda |
| DL-M Day 0 | Deer Lake | soda lake | North America | Cariboo Plateau | soda |
| DLM 2015 | Deer Lake | soda lake | North America | Cariboo Plateau | soda |
| DLM 2017 | Deer Lake | soda lake | North America | Cariboo Plateau | soda |
| GEL-M Day 0 | Goodenough Lake | soda lake | North America | Cariboo Plateau | soda |
| GEM 2015 | Goodenough Lake | soda lake | North America | Cariboo Plateau | soda |
| GEM 2016 | Goodenough Lake | soda lake | North America | Cariboo Plateau | soda |
| GEM 2017 | Goodenough Lake | soda lake | North America | Cariboo Plateau | soda |

| | | | | |
|---|---|---|---|---|
| LCL-M Day 0 | Last Chance Lake | soda lake | North America Cariboo Plateau | soda |
| LCM 2015 | Last Chance Lake | soda lake | North America Cariboo Plateau | soda |
| LCM 2016 | Last Chance Lake | soda lake | North America Cariboo Plateau | soda |
| LCM 2017 | Last Chance Lake | soda lake | North America Cariboo Plateau | soda |
| PL-M Day 0 | Probe Lake | soda lake | North America Cariboo Plateau | soda |
| PLM 2015 | Probe Lake | soda lake | North America Cariboo Plateau | soda |
| PLM 2016 | Probe Lake | soda lake | North America Cariboo Plateau | soda |
| PLM 2017 | Probe Lake | soda lake | North America Cariboo Plateau | soda |
| ML16S_10m | Mono Lake | soda lake | North America California | soda |
| MLW_0517_00 | Mono Lake | soda lake | North America California | soda |
| MLW_0517_10 | Mono Lake | soda lake | North America California | soda |
| MLW_0617_00 | Mono Lake | soda lake | North America California | soda |
| MLW_0917_00_1;MLW_0917_00_2;MLW_0917_00_3 | Mono Lake | soda lake | North America California | soda |
| MLW_0917_05_1;MLW_0917_05_2;MLW_0917_05_3 | Mono Lake | soda lake | North America California | soda |
| MLW_0917_10_1;MLW_0917_10_2;MLW_0917_10_3 | Mono Lake | soda lake | North America California | soda |
| MLW_1018_00 | Mono Lake | soda lake | North America California | soda |
| MLW_1018_05 | Mono Lake | soda lake | North America California | soda |
| MLW_1018_12 | Mono Lake | soda lake | North America California | soda |

n.d. - not determined

*defined according to Boros & Kolpakova 2018

| Sample ID | $Na^+$ (mg/L) | $K^+$ (mg/L) | $Ca^{2+}$ (mg/L) | $Mg^{2+}$ (mg/L) | $Cl^-$ (mg/L) | $SO_4^{2-}$ (mg/L) | $HCO_3^-$ (mg/L) | $CO_3^{2-}$ (mg/L) |
|---|---|---|---|---|---|---|---|---|
| Bac-12S | 2423 | 309 | 3 | 2 | 268 | 100 | 4033 | 1044 |
| Bac-18E | 3793 | 274 | 0 | 0 | 1985 | 144 | 6544 | 440 |
| Bac-6M | 12835 | 228 | 4 | 12 | 5244 | 100 | 6093 | 9589 |
| B1-W | 25860 | 414 | 4 | 0 | 5240 | 148 | 34950 | 12700 |
| B2-W | 25860 | 414 | 4 | 0 | 5240 | 148 | 34950 | 12700 |
| KZ02 | 4320 | 44 | 392 | 309 | 3550 | 7560 | 153 | 24 |
| K03bact | 26930 | 830 | 11 | 444 | 40825 | 8200 | 1830 | 120 |
| KZ04 | 25910 | 200 | 127 | 1308 | 37275 | 4720 | 171 | 0 |
| KZ05 | 66 | 50 | 87 | 38 | 142 | 300 | 18 | 0 |
| K06bact | 43410 | 450 | 646 | 4950 | 79875 | 18960 | 390 | 0 |
| KZ08 | 25980 | 680 | 498 | 2533 | 35500 | 22870 | 171 | 12 |
| K10bact | 4050 | 101 | 101 | 535 | 7455 | 1370 | 329 | 48 |
| KZ12 | 5790 | 68 | 385 | 836 | 4260 | 11429 | 354 | 0 |
| K13bact | 4490 | 100 | 650 | 566 | 4970 | 2530 | 98 | 0 |
| K15bact | 12150 | 59 | 1049 | 5043 | 26625 | 4090 | 61 | 12 |
| K16bact | 1510 | 49 | 122 | 630 | 3195 | 680 | 165 | 0 |
| K18bact | 238 | 32 | 233 | 53 | 604 | 630 | 122 | 0 |
| KZ19 | 39070 | 420 | 479 | 4438 | 58575 | 27490 | 317 | 0 |
| K20bact | 44 | 16 | 90 | 15 | 60 | 166 | 146 | 0 |
| KT01 | 1270 | 109 | 28 | 343 | 1189 | 1879 | 769 | 78 |
| KT02 | 25960 | 380 | 28 | 2658 | 33725 | 18819 | 2885 | 522 |
| KT03 | 1326 | 52 | 28 | 372 | 1350 | 1744 | 830 | 120 |
| KT04 | 939 | 49 | 30 | 237 | 888 | 1301 | 537 | 72 |
| KT05 | 1944 | 35 | 34 | 660 | 2556 | 2452 | 1269 | 216 |
| KT06 | 1000 | 78 | 22 | 260 | 1030 | 1407 | 537 | 120 |
| KT07 | 1060 | 121 | 26 | 294 | 1030 | 1551 | 634 | 72 |
| KT08 | 1128 | 127 | 20 | 292 | 1065 | 1771 | 647 | 72 |
| KT09 | 8933 | 396 | 1278 | 5670 | 22543 | 11286 | 537 | 84 |
| KT10 | 17000 | 259 | 32 | 297 | 7313 | 18531 | 6124 | 1584 |
| KT11 | 675 | 17 | 40 | 116 | 604 | 522 | 464 | 48 |
| KT12 | 6340 | 103 | 9 | 159 | 2095 | 5603 | 3880 | 1128 |
| KT13 | 113000 | 1750 | 88 | 238 | 10206 | 51913 | 166123 | 24840 |

| | | | | | | | | |
|---|---|---|---|---|---|---|---|---|
| KT14 | 124000 | 2530 | 100 | 5411 | 145550 | 86687 | 5173 | 648 |
| KT15 | 90000 | 2310 | 130 | 6123 | 70290 | 108497 | 1440 | 336 |
| KT16 | 1785 | 153 | 356 | 34 | 1420 | 2151 | 708 | 144 |
| KT17 | 143000 | 5090 | 687 | 21471 | 94075 | 255800 | 4807 | 852 |
| KT18 | 2458 | 133 | 20 | 341 | 1243 | 1677 | 537 | 1680 |
| KT19 | 595 | 39 | 22 | 54 | 405 | 647 | 268 | 96 |
| KT20 | 363 | 46 | 30 | 116 | 293 | 429 | 561 | 84 |
| KT21 | 380 | 7 | 36 | 32 | 568 | 64 | 256 | 0 |
| KT22 | 1750 | 98 | 32 | 601 | 284 | 4101 | 2135 | 240 |
| KT23 | 2056 | 50 | 55 | 305 | 1420 | 2583 | 915 | 240 |
| KT24 | 1830 | 46 | 34 | 308 | 1420 | 2023 | 1068 | 120 |
| KT25 | 2590 | 52 | 12 | 376 | 1775 | 3247 | 1159 | 240 |
| KT26 | 2008 | 39 | 28 | 263 | 1207 | 2254 | 854 | 120 |
| KT27 | 2490 | 72 | 14 | 393 | 2059 | 2897 | 1251 | 270 |
| 0S | n.d. | n.d. | n.d. | n.d. | n.d. | n.d. | n.d. | n.d. |
| 02S | n.d. | n.d. | n.d. | n.d. | n.d. | n.d. | n.d. | n.d. |
| 04S | n.d. | n.d. | n.d. | n.d. | n.d. | n.d. | n.d. | n.d. |
| 06S | n.d. | n.d. | n.d. | n.d. | n.d. | n.d. | n.d. | n.d. |
| 08S | n.d. | n.d. | n.d. | n.d. | n.d. | n.d. | n.d. | n.d. |
| 09S | n.d. | n.d. | n.d. | n.d. | n.d. | n.d. | n.d. | n.d. |
| 11S | n.d. | n.d. | n.d. | n.d. | n.d. | n.d. | n.d. | n.d. |
| SHV | 695 | 24 | 242 | 59 | 489 | 299 | 720 | 30 |
| KA1 | 19560 | 46 | 116 | 622 | 18193 | 19334 | 67 | 228 |
| KA2 | 3238 | 28 | 28 | 102 | 3527 | 907 | 1635 | 138 |
| KA3 | 15125 | 102 | 80 | 770 | 17519 | 10583 | 1006 | 324 |
| KA4 | 44000 | 828 | 644 | 2034 | 95 | 55695 | 476 | 126 |
| KA5 | 187 | 8 | 36 | 156 | 181 | 140 | 311 | 0 |
| KA6 | 83440 | 182 | 476 | 3395 | 105403 | 53921 | 921 | 0 |
| KA7 | 6488 | 132 | 1752 | 2005 | 13559 | 8147 | 329 | 0 |
| KA8 | 648 | 18 | 192 | 347 | 407 | 1818 | 327 | 0 |
| Gudzh_water | 35400 | n.d. | 30 | 440 | 7130 | 55900 | 9760 | 1500 |
| NB30 | 8030 | 69 | 13 | 26 | 4086 | 145 | 7961 | 6210 |
| NB31 | 8030 | 69 | 13 | 26 | 4086 | 145 | 7961 | 6210 |

| | | | | | | | | |
|---|---|---|---|---|---|---|---|---|
| NB32 | 8030 | 69 | 13 | 26 | 4086 | 145 | 7961 | 6210 |
| NB41 | 8030 | 69 | 13 | 26 | 4086 | 145 | 7961 | 6210 |
| NB42 | 8030 | 69 | 13 | 26 | 4086 | 145 | 7961 | 6210 |
| NB43 | 8030 | 69 | 13 | 26 | 4086 | 145 | 7961 | 6210 |
| Bangong Co | n.d. | n.d. | n.d. | n.d. | 83 | 103 | n.d. | n.d. |
| Bong Co | 26 | 21 | 19 | 12 | 14 | 112 | 46 | 3 |
| Co Ngoin (1) | 30 | 0 | 20 | 40 | 10 | 20 | 280 | n.d. |
| Mapam Yumco | 52 | 6 | 28 | 31 | 14 | 32 | 323 | n.d. |
| Urru Co | n.d. | n.d. | n.d. | n.d. | n.d. | n.d. | n.d. | n.d. |
| Bam Co | 2533 | 292 | 76 | 196 | 1362 | 4195 | 483 | 112 |
| Bero Zeco | 32000 | 3000 | 0 | 5000 | 4000 | 65000 | 2000 | 0 |
| Co Ngoin (2) | 1478 | 146 | 6 | 118 | 809 | 1174 | 1011 | 629 |
| Dawa Co | 7390 | 945 | 57 | 542 | 3155 | 14193 | 272 | 112 |
| Tangra Yumco | n.d. | n.d. | n.d. | n.d. | n.d. | n.d. | n.d. | n.d. |
| Dong Co | 16576 | 5011 | 95 | 993 | 11115 | 29610 | 277 | 13 |
| Kunggyu | 1387 | 297 | 148 | 108 | 972 | 52 | 2057 | 1294 |
| Nam Co | 770 | 0 | 10 | 90 | 80 | 270 | 1330 | 310 |
| Pung Co | 2881 | 357 | 76 | 242 | 861 | 3997 | 2176 | 445 |
| Selin Co | 5550 | 580 | 0 | 260 | 3390 | 6980 | 1440 | 410 |
| Zhari Namco | 3970 | 353 | 57 | 231 | 1219 | 7567 | 396 | 147 |
| Zhaxi Co | 5170 | 1006 | 57 | 358 | 2151 | 9870 | 580 | 208 |
| Bangkog Co | 8288 | 1252 | 57 | 346 | 6310 | 11284 | 400 | 125 |
| Dangqiong Co | 54000 | 7000 | 0 | 0 | 68000 | 7000 | 0 | 11000 |
| Nyer Co (Nieer Co) | 40000 | 17000 | 0 | 16000 | 92000 | 43000 | 0 | 0 |
| BU1 | 1380 | 12 | 12 | 12 | 415 | 150 | 2003 | 260 |
| SO1 | 914 | 0 | 8 | 9 | 505 | 157 | 1213 | 73 |
| ZA1 | 1722 | 13 | 15 | 15 | 723 | 256 | 2295 | 429 |
| SO2 | 914 | 0 | 8 | 9 | 505 | 157 | 1213 | 73 |
| SO3 | 914 | 0 | 8 | 9 | 505 | 157 | 1213 | 73 |
| SO4 | 914 | 0 | 8 | 9 | 505 | 157 | 1213 | 73 |
| SO5 | 914 | 0 | 8 | 9 | 505 | 157 | 1213 | 73 |
| SO6 | 914 | 0 | 8 | 9 | 505 | 157 | 1213 | 73 |
| SO9 | 914 | 0 | 8 | 9 | 505 | 157 | 1213 | 73 |

| | | | | | | | | |
|---|---|---|---|---|---|---|---|---|
| SO11 | 914 | 0 | 8 | 9 | 505 | 157 | 1213 | 73 |
| SO12 | 914 | 0 | 8 | 9 | 505 | 157 | 1213 | 73 |
| SO13 | 914 | 0 | 8 | 9 | 505 | 157 | 1213 | 73 |
| SO14 | 914 | 0 | 8 | 9 | 505 | 157 | 1213 | 73 |
| SO15 | 914 | 0 | 8 | 9 | 505 | 157 | 1213 | 73 |
| SO16 | 914 | 0 | 8 | 9 | 505 | 157 | 1213 | 73 |
| VBK | 1145 | 8 | 7 | 9 | 592 | 255 | 1893 | 47 |
| ZA2 | 1722 | 13 | 15 | 15 | 723 | 256 | 2295 | 429 |
| ZA3 | 1722 | 13 | 15 | 15 | 723 | 256 | 2295 | 429 |
| ZA4 | 1722 | 13 | 15 | 15 | 723 | 256 | 2295 | 429 |
| ZA5 | 1722 | 13 | 15 | 15 | 723 | 256 | 2295 | 429 |
| ZA6 | 1722 | 13 | 15 | 15 | 723 | 256 | 2295 | 429 |
| ZA7 | 1722 | 13 | 15 | 15 | 723 | 256 | 2295 | 429 |
| ZA8 | 1722 | 13 | 15 | 15 | 723 | 256 | 2295 | 429 |
| ZA9 | 1722 | 13 | 15 | 15 | 723 | 256 | 2295 | 429 |
| ZA10 | 1722 | 13 | 15 | 15 | 723 | 256 | 2295 | 429 |
| ZA11 | 1722 | 13 | 15 | 15 | 723 | 256 | 2295 | 429 |
| ZA12 | 1722 | 13 | 15 | 15 | 723 | 256 | 2295 | 429 |
| ZA13 | 1722 | 13 | 15 | 15 | 723 | 256 | 2295 | 429 |
| ZA14 | 1722 | 13 | 15 | 15 | 723 | 256 | 2295 | 429 |
| ZA15 | 1722 | 13 | 15 | 15 | 723 | 256 | 2295 | 429 |
| ZA16 | 1722 | 13 | 15 | 15 | 723 | 256 | 2295 | 429 |
| ZA17 | 1722 | 13 | 15 | 15 | 723 | 256 | 2295 | 429 |
| M05 | 26300 | 32 | 112 | 20 | 41000 | 231 | 366 | 200 |
| U05 | n.d. | n.d. | n.d. | n.d. | 19000 | 126 | n.d. | n.d. |
| U15 | n.d. | n.d. | n.d. | n.d. | 39000 | 120 | n.d. | n.d. |
| U28 | n.d. | n.d. | n.d. | n.d. | 60000 | 126 | n.d. | n.d. |
| B01w | 426 | 34 | 16 | 156 | 329 | 524 | 769 | 29 |
| B02w | 426 | 34 | 16 | 156 | 329 | 524 | 769 | 29 |
| B03w | 426 | 34 | 16 | 156 | 329 | 524 | 769 | 29 |
| B04w | 426 | 34 | 16 | 156 | 329 | 524 | 769 | 29 |
| B05w | 426 | 34 | 16 | 156 | 329 | 524 | 769 | 29 |
| B06w | 426 | 34 | 16 | 156 | 329 | 524 | 769 | 29 |

| | | | | | | | | |
|---|---|---|---|---|---|---|---|---|
| B07w | 426 | 34 | 16 | 156 | 329 | 524 | 769 | 29 |
| B08w | 426 | 34 | 16 | 156 | 329 | 524 | 769 | 29 |
| B09w | 426 | 34 | 16 | 156 | 329 | 524 | 769 | 29 |
| B010w | 426 | 34 | 16 | 156 | 329 | 524 | 769 | 29 |
| B011w | 426 | 34 | 16 | 156 | 329 | 524 | 769 | 29 |
| KH1w | 424 | 35 | 26 | 158 | 315 | 500 | 769 | 58 |
| KH2w | 424 | 35 | 26 | 158 | 315 | 500 | 769 | 58 |
| KH3w | 424 | 35 | 26 | 158 | 315 | 500 | 769 | 58 |
| KH4w | 424 | 35 | 26 | 158 | 315 | 500 | 769 | 58 |
| KH5w | 424 | 35 | 26 | 158 | 315 | 500 | 769 | 58 |
| KH6w | 424 | 35 | 26 | 158 | 315 | 500 | 769 | 58 |
| KH7w | 424 | 35 | 26 | 158 | 315 | 500 | 769 | 58 |
| KH8w | 424 | 35 | 26 | 158 | 315 | 500 | 769 | 58 |
| KH9w | 424 | 35 | 26 | 158 | 315 | 500 | 769 | 58 |
| KH10w | 424 | 35 | 26 | 158 | 315 | 500 | 769 | 58 |
| KH11w | 424 | 35 | 26 | 158 | 315 | 500 | 769 | 58 |
| B01 | 1567 | 9 | 15 | 18 | 1101 | 280 | 1314 | 249 |
| B02 | 1567 | 9 | 15 | 18 | 1101 | 280 | 1314 | 249 |
| B03 | 1567 | 9 | 15 | 18 | 1101 | 280 | 1314 | 249 |
| B04 | 1567 | 9 | 15 | 18 | 1101 | 280 | 1314 | 249 |
| B05 | 1567 | 9 | 15 | 18 | 1101 | 280 | 1314 | 249 |
| B06 | 1567 | 9 | 15 | 18 | 1101 | 280 | 1314 | 249 |
| B07 | 1567 | 9 | 15 | 18 | 1101 | 280 | 1314 | 249 |
| B08 | 1567 | 9 | 15 | 18 | 1101 | 280 | 1314 | 249 |
| B09 | 1567 | 9 | 15 | 18 | 1101 | 280 | 1314 | 249 |
| B10 | 1567 | 9 | 15 | 18 | 1101 | 280 | 1314 | 249 |
| B11 | 1567 | 9 | 15 | 18 | 1101 | 280 | 1314 | 249 |
| B12 | 1567 | 9 | 15 | 18 | 1101 | 280 | 1314 | 249 |
| B13 | 1567 | 9 | 15 | 18 | 1101 | 280 | 1314 | 249 |
| B14 | 1567 | 9 | 15 | 18 | 1101 | 280 | 1314 | 249 |
| K01 | 1192 | 12 | 15 | 16 | 680 | 183 | 1290 | 296 |
| K02 | 1192 | 12 | 15 | 16 | 680 | 183 | 1290 | 296 |
| K03 | 1192 | 12 | 15 | 16 | 680 | 183 | 1290 | 296 |

| | | | | | | | | |
|---|---|---|---|---|---|---|---|---|
| K04 | 1192 | 12 | 15 | 16 | 680 | 183 | 1290 | 296 |
| K05 | 1192 | 12 | 15 | 16 | 680 | 183 | 1290 | 296 |
| K06 | 1192 | 12 | 15 | 16 | 680 | 183 | 1290 | 296 |
| K09 | 1192 | 12 | 15 | 16 | 680 | 183 | 1290 | 296 |
| K12 | 1192 | 12 | 15 | 16 | 680 | 183 | 1290 | 296 |
| K14 | 1192 | 12 | 15 | 16 | 680 | 183 | 1290 | 296 |
| S01 | 914 | 0 | 8 | 9 | 505 | 157 | 1213 | 73 |
| S02 | 914 | 0 | 8 | 9 | 505 | 157 | 1213 | 73 |
| S03 | 914 | 0 | 8 | 9 | 505 | 157 | 1213 | 73 |
| S04 | 914 | 0 | 8 | 9 | 505 | 157 | 1213 | 73 |
| S05 | 914 | 0 | 8 | 9 | 505 | 157 | 1213 | 73 |
| S06 | 914 | 0 | 8 | 9 | 505 | 157 | 1213 | 73 |
| S07 | 914 | 0 | 8 | 9 | 505 | 157 | 1213 | 73 |
| S08 | 914 | 0 | 8 | 9 | 505 | 157 | 1213 | 73 |
| S09 | 914 | 0 | 8 | 9 | 505 | 157 | 1213 | 73 |
| S10 | 914 | 0 | 8 | 9 | 505 | 157 | 1213 | 73 |
| S11 | 914 | 0 | 8 | 9 | 505 | 157 | 1213 | 73 |
| S12 | 914 | 0 | 8 | 9 | 505 | 157 | 1213 | 73 |
| S13 | 914 | 0 | 8 | 9 | 505 | 157 | 1213 | 73 |
| S14 | 914 | 0 | 8 | 9 | 505 | 157 | 1213 | 73 |
| V01 | 1145 | 8 | 7 | 9 | 592 | 255 | 1893 | 47 |
| V02 | 1145 | 8 | 7 | 9 | 592 | 255 | 1893 | 47 |
| V03 | 1145 | 8 | 7 | 9 | 592 | 255 | 1893 | 47 |
| V04 | 1145 | 8 | 7 | 9 | 592 | 255 | 1893 | 47 |
| V05 | 1145 | 8 | 7 | 9 | 592 | 255 | 1893 | 47 |
| V06 | 1145 | 8 | 7 | 9 | 592 | 255 | 1893 | 47 |
| V07 | 1145 | 8 | 7 | 9 | 592 | 255 | 1893 | 47 |
| V08 | 1145 | 8 | 7 | 9 | 592 | 255 | 1893 | 47 |
| V09 | 1145 | 8 | 7 | 9 | 592 | 255 | 1893 | 47 |
| V10 | 1145 | 8 | 7 | 9 | 592 | 255 | 1893 | 47 |
| V11 | 1145 | 8 | 7 | 9 | 592 | 255 | 1893 | 47 |
| V12 | 1145 | 8 | 7 | 9 | 592 | 255 | 1893 | 47 |
| V13 | 1145 | 8 | 7 | 9 | 592 | 255 | 1893 | 47 |

| | | | | | | | | |
|---|---|---|---|---|---|---|---|---|
| V14 | 1145 | 8 | 7 | 9 | 592 | 255 | 1893 | 47 |
| Z01 | 1722 | 13 | 15 | 15 | 723 | 256 | 2295 | 429 |
| Z02 | 1722 | 13 | 15 | 15 | 723 | 256 | 2295 | 429 |
| Z03 | 1722 | 13 | 15 | 15 | 723 | 256 | 2295 | 429 |
| Z04 | 1722 | 13 | 15 | 15 | 723 | 256 | 2295 | 429 |
| Z05 | 1722 | 13 | 15 | 15 | 723 | 256 | 2295 | 429 |
| Z06 | 1722 | 13 | 15 | 15 | 723 | 256 | 2295 | 429 |
| Z07 | 1722 | 13 | 15 | 15 | 723 | 256 | 2295 | 429 |
| Z08 | 1722 | 13 | 15 | 15 | 723 | 256 | 2295 | 429 |
| Z09 | 1722 | 13 | 15 | 15 | 723 | 256 | 2295 | 429 |
| Z10 | 1722 | 13 | 15 | 15 | 723 | 256 | 2295 | 429 |
| Z12 | 1722 | 13 | 15 | 15 | 723 | 256 | 2295 | 429 |
| Z14 | 1722 | 13 | 15 | 15 | 723 | 256 | 2295 | 429 |
| BV | 31 | 7 | 47 | 47 | 30 | 115 | 251 | 15 |
| K0 | n.d. | n.d. | n.d. | n.d. | n.d. | n.d. | n.d. | n.d. |
| K01 | n.d. | n.d. | n.d. | n.d. | n.d. | n.d. | n.d. | n.d. |
| A | n.d. | n.d. | n.d. | n.d. | n.d. | n.d. | n.d. | n.d. |
| B | n.d. | n.d. | n.d. | n.d. | n.d. | n.d. | n.d. | n.d. |
| B0 | 426 | 34 | 16 | 156 | 329 | 524 | 769 | 29 |
| C | n.d. | n.d. | n.d. | n.d. | n.d. | n.d. | n.d. | n.d. |
| D | n.d. | n.d. | n.d. | n.d. | n.d. | n.d. | n.d. | n.d. |
| Ds | n.d. | n.d. | n.d. | n.d. | n.d. | n.d. | n.d. | n.d. |
| E | n.d. | n.d. | n.d. | n.d. | n.d. | n.d. | n.d. | n.d. |
| F | n.d. | n.d. | n.d. | n.d. | n.d. | n.d. | n.d. | n.d. |
| FDN | n.d. | n.d. | n.d. | n.d. | n.d. | n.d. | n.d. | n.d. |
| FDO | n.d. | n.d. | n.d. | n.d. | n.d. | n.d. | n.d. | n.d. |
| Ff | 34 | 8 | 37 | 54 | 33 | 137 | 228 | 19 |
| K | 31 | 7 | 47 | 47 | 30 | 115 | 251 | 15 |
| KH | 424 | 35 | 26 | 158 | 315 | 500 | 769 | 58 |
| Kn | n.d. | n.d. | n.d. | n.d. | n.d. | n.d. | n.d. | n.d. |
| LK | n.d. | n.d. | n.d. | n.d. | n.d. | n.d. | n.d. | n.d. |
| LN | n.d. | n.d. | n.d. | n.d. | n.d. | n.d. | n.d. | n.d. |
| NH | 424 | 35 | 26 | 158 | 315 | 500 | 769 | 58 |

| | | | | | | | | |
|---|---|---|---|---|---|---|---|---|
| R | n.d. | n.d. | n.d. | n.d. | n.d. | n.d. | n.d. | n.d. |
| TLO | n.d. | n.d. | n.d. | n.d. | n.d. | n.d. | n.d. | n.d. |
| TNB | n.d. | n.d. | n.d. | n.d. | n.d. | n.d. | n.d. | n.d. |
| TO | n.d. | n.d. | n.d. | n.d. | n.d. | n.d. | n.d. | n.d. |
| TS | n.d. | n.d. | n.d. | n.d. | n.d. | n.d. | n.d. | n.d. |
| Z | 31 | 5 | 78 | 36 | 30 | 69 | 381 | 0 |
| SS15 | 2068 | 121 | 8 | 101 | 1148 | 500 | 3552 | 212 |
| SS16 | 2068 | 121 | 8 | 101 | 1148 | 500 | 3552 | 212 |
| SS17 | 2068 | 121 | 8 | 101 | 1148 | 500 | 3552 | 212 |
| US15 | 584 | 18 | 14 | 77 | 149 | 87 | 1438 | 87 |
| US16 | 584 | 18 | 14 | 77 | 149 | 87 | 1438 | 87 |
| US17 | 584 | 18 | 14 | 77 | 149 | 87 | 1438 | 87 |
| ZL15 | 1257 | 69 | 15 | 89 | 551 | 1036 | 1755 | 128 |
| ZL16 | 1257 | 69 | 15 | 89 | 551 | 1036 | 1755 | 128 |
| ZL17 | 1257 | 69 | 15 | 89 | 551 | 1036 | 1755 | 128 |
| Sp17_01 | 773 | 54 | 13 | 102 | 361 | 145 | 1370 | 85 |
| Sp17_02 | 538 | 4 | 8 | 8 | 59 | 135 | 1193 | 91 |
| Sp17_04 | 251 | 16 | 124 | 50 | 23 | 110 | 1052 | 20 |
| Sp17_05 | 1766 | 93 | 0 | 550 | 1305 | 2067 | 1893 | 291 |
| Sp17_07 | 1320 | 13 | 12 | 62 | 287 | 393 | 2286 | 140 |
| Sp17_09 | n.d. | n.d. | n.d. | n.d. | n.d. | n.d. | n.d. | n.d. |
| Sp17_10 | 597 | 7 | 15 | 32 | 118 | 85 | 1049 | 62 |
| Sp17_11 | 423 | 21 | 22 | 71 | 105 | 305 | 741 | 75 |
| Sp17_12 | 521 | 7 | 7 | 24 | 73 | 57 | 1088 | 74 |
| Sp17_13 | 2418 | 93 | 4 | 85 | 720 | 498 | 5877 | 370 |
| Sp17_14 | n.d. | n.d. | n.d. | n.d. | n.d. | n.d. | n.d. | n.d. |
| Sp17_15 | 722 | 49 | 16 | 249 | 516 | 849 | 1124 | 87 |
| Sp17_16 | 712 | 18 | 13 | 77 | 166 | 143 | 1327 | 96 |
| Sp17_17 | 2090 | 77 | 3 | 53 | 642 | 397 | 4856 | 308 |
| Sp17_18 | 5350 | 51 | 8 | 69 | 1130 | 1814 | 17636 | 1019 |
| Sp17_19 | 574 | 7 | 12 | 41 | 143 | 78 | 1160 | 87 |
| Sp17_20 | 350 | 17 | 32 | 66 | 119 | 353 | 667 | 61 |
| Sp17_21 | 1735 | 65 | 4 | 94 | 561 | 428 | 3518 | 217 |

| | | | | | | | | |
|---|---|---|---|---|---|---|---|---|
| Sp17_22 | 701 | 3 | 11 | 34 | 154 | 130 | 1339 | 80 |
| Sp17_23 | 383 | 12 | 13 | 45 | 116 | 87 | 912 | 67 |
| Sp17_24 | 2068 | 121 | 8 | 101 | 1148 | 500 | 3552 | 212 |
| Sp17_27 | 719 | 10 | 10 | 6 | 175 | 400 | 1268 | 49 |
| Sp17_28 | 378 | 19 | 27 | 78 | 110 | 243 | 717 | 70 |
| Sp17_29 | 1257 | 69 | 15 | 89 | 551 | 1036 | 1755 | 128 |
| Sp17_31 | 1766 | 93 | 0 | 550 | 1305 | 2067 | 1893 | 291 |
| Sp17_Neu2 | n.d. | n.d. | n.d. | n.d. | n.d. | n.d. | n.d. | n.d. |
| Sp18_01 | 773 | 54 | 13 | 102 | 361 | 145 | 1370 | 85 |
| Sp18_04 | 251 | 16 | 124 | 50 | 23 | 110 | 1052 | 20 |
| Sp18_05 | 1766 | 93 | 0 | 550 | 1305 | 2067 | 1893 | 291 |
| Sp18_07 | 1320 | 13 | 12 | 62 | 287 | 393 | 2286 | 140 |
| Sp18_08 | 692 | 63 | 8 | 71 | 245 | 101 | 1336 | 103 |
| Sp18_10 | 597 | 7 | 15 | 32 | 118 | 85 | 1049 | 62 |
| Sp18_11 | 423 | 21 | 22 | 71 | 105 | 305 | 741 | 75 |
| Sp18_12 | 521 | 7 | 7 | 24 | 73 | 57 | 1088 | 74 |
| Sp18_13 | 2418 | 93 | 4 | 85 | 720 | 498 | 5877 | 370 |
| Sp18_15 | 722 | 49 | 16 | 249 | 516 | 849 | 1124 | 87 |
| Sp18_16 | 712 | 18 | 13 | 77 | 166 | 143 | 1327 | 96 |
| Sp18_17 | 2090 | 77 | 3 | 53 | 642 | 397 | 4856 | 308 |
| Sp18_18 | 5350 | 51 | 8 | 69 | 1130 | 1814 | 17636 | 1019 |
| Sp18_19 | 574 | 7 | 12 | 41 | 143 | 78 | 1160 | 87 |
| Sp18_1_d | 538 | 4 | 8 | 8 | 59 | 135 | 1193 | 91 |
| Sp18_20 | 350 | 17 | 32 | 66 | 119 | 353 | 667 | 61 |
| Sp18_21 | 1735 | 65 | 4 | 94 | 561 | 428 | 3518 | 217 |
| Sp18_22 | 701 | 3 | 11 | 34 | 154 | 130 | 1339 | 80 |
| Sp18_23 | 383 | 12 | 13 | 45 | 116 | 87 | 912 | 67 |
| Sp18_24 | 2068 | 121 | 8 | 101 | 1148 | 500 | 3552 | 212 |
| Sp18_27 | 719 | 10 | 10 | 6 | 175 | 400 | 1268 | 49 |
| Sp18_28 | 378 | 19 | 27 | 78 | 110 | 243 | 717 | 70 |
| Sp18_29 | 1257 | 69 | 15 | 89 | 551 | 1036 | 1755 | 128 |
| Sp18_30 | 774 | 45 | 13 | 65 | 317 | 1000 | 589 | 82 |
| Sp18_31 | 2159 | 46 | 21 | 263 | 609 | 1046 | 1957 | 246 |

| | | | | | | | | |
|---|---|---|---|---|---|---|---|---|
| Sp18_9 | n.d. | n.d. | n.d. | n.d. | n.d. | n.d. | n.d. | n.d. |
| Sp18_Neu2 | n.d. | n.d. | n.d. | n.d. | n.d. | n.d. | n.d. | n.d. |
| 01Boddi | 1567 | 9 | 15 | 18 | 1101 | 280 | 1314 | 249 |
| 02Soser | 914 | 0 | 8 | 9 | 505 | 157 | 1213 | 73 |
| 03KolonNy | n.d. | n.d. | n.d. | n.d. | n.d. | n.d. | n.d. | n.d. |
| 04KolonT18 | n.d. | n.d. | n.d. | n.d. | n.d. | n.d. | n.d. | n.d. |
| 05Szelid18 | 279 | 3 | 18 | 31 | 250 | 35 | 420 | 80 |
| 06Rusanda18 | 1159 | 39 | 13 | 15 | 616 | 928 | 1252 | 55 |
| 07SlanoKopovo18 | 552 | 5 | 12 | 8 | 484 | 285 | 525 | 0 |
| 08VBK18 | 1145 | 8 | 7 | 9 | 592 | 255 | 1893 | 47 |
| 09Kelemen18 | 1192 | 12 | 15 | 16 | 680 | 183 | 1290 | 296 |
| 10Zab18 | 1722 | 13 | 15 | 15 | 723 | 256 | 2295 | 429 |
| 12KH18 | 424 | 35 | 26 | 158 | 315 | 500 | 769 | 58 |
| 13FertoB018 | 426 | 34 | 16 | 156 | 329 | 524 | 769 | 29 |
| 14VelenceiKo18 | 662 | 82 | 0 | 416 | 559 | 1104 | 1124 | 182 |
| 15Dinnyesi18 | 775 | 78 | 0 | 355 | 641 | 754 | 1376 | 218 |
| 16VelenceiNy18 | 688 | 85 | 0 | 427 | 580 | 1144 | 1095 | 189 |
| 17Zicklacke18 | 1257 | 69 | 15 | 89 | 551 | 1036 | 1755 | 128 |
| 18ObererStinker18 | 2090 | 77 | 3 | 53 | 642 | 397 | 4856 | 308 |
| 19SudlicherSilbersee18 | 2068 | 121 | 8 | 101 | 1148 | 500 | 3552 | 212 |
| 20LangeLacke18 | 423 | 21 | 22 | 71 | 105 | 305 | 741 | 75 |
| 21BalatonTihany18 | 34 | 8 | 37 | 54 | 33 | 137 | 228 | 19 |
| 22BalatonKeszthely18 | 31 | 7 | 47 | 47 | 30 | 115 | 251 | 15 |
| 23UntererStinker18 | 584 | 18 | 14 | 77 | 149 | 87 | 1438 | 87 |
| 01T | 4298 | 77 | 0 | 67 | 5024 | 2234 | 503 | 87 |
| 02T | 2853 | 360 | 0 | 443 | 1946 | 3966 | 1198 | 291 |
| 03T | 950 | 0 | 13 | 16 | 304 | 261 | 1435 | 138 |
| 04T | 403 | 5 | 0 | 95 | 127 | 99 | 873 | 160 |
| 05T | 451 | 0 | 8 | 26 | 195 | 30 | 843 | 58 |
| 06T | 914 | 0 | 8 | 9 | 505 | 157 | 1213 | 73 |
| 07T | 1845 | 0 | 2 | 0 | 1528 | 359 | 1479 | 262 |
| 08T | 1584 | 0 | 1 | 0 | 1309 | 169 | 1317 | 182 |
| 09T | 432 | 0 | 30 | 3 | 243 | 54 | 666 | 36 |

| | | | | | | | | |
|---|---|---|---|---|---|---|---|---|
| 10T | 1122 | 0 | 0 | 12 | 646 | 149 | 1183 | 233 |
| 11T | 361 | 4 | 42 | 7 | 120 | 41 | 900 | 36 |
| 12T | 408 | 0 | 28 | 3 | 188 | 49 | 740 | 36 |
| 13T | 489 | 0 | 17 | 3 | 230 | 56 | 950 | 36 |
| 14T | 1567 | 0 | 9 | 17 | 593 | 379 | 2189 | 204 |
| 15T | 1594 | 0 | 15 | 25 | 591 | 378 | 2248 | 233 |
| 16T | 17 | 2 | 45 | 16 | 23 | 35 | 185 | 0 |
| 17T | 255 | 5 | 28 | 16 | 52 | 124 | 670 | 0 |
| 18T | 96 | 2 | 55 | 23 | 33 | 21 | 414 | 0 |
| 19T | 722 | 49 | 16 | 249 | 516 | 849 | 1124 | 87 |
| 20T | 1766 | 93 | 0 | 550 | 1305 | 2067 | 1893 | 291 |
| 21T | 424 | 35 | 26 | 158 | 315 | 500 | 769 | 58 |
| 22T | 410 | 33 | 16 | 151 | 315 | 502 | 769 | 29 |
| 23T | 426 | 34 | 16 | 156 | 329 | 524 | 769 | 29 |
| 24T | 432 | 36 | 28 | 164 | 315 | 498 | 828 | 29 |
| 25T | 464 | 37 | 0 | 166 | 355 | 568 | 828 | 58 |
| 26T | 279 | 3 | 18 | 31 | 250 | 35 | 420 | 80 |
| 27T | 294 | 2 | 27 | 33 | 265 | 37 | 450 | 90 |
| 28T | 688 | 85 | 0 | 427 | 580 | 1144 | 1095 | 189 |
| 29T | 662 | 82 | 0 | 416 | 559 | 1104 | 1124 | 182 |
| 30T | 602 | 74 | 0 | 393 | 511 | 1014 | 1139 | 153 |
| 31T | 775 | 78 | 0 | 355 | 641 | 754 | 1376 | 218 |
| DL-M Day 0 | 7296 | 97 | 3 | 34 | 128 | 21 | 11896 | 3421 |
| DLM 2015 | 7296 | 97 | 3 | 34 | 128 | 21 | 11896 | 3421 |
| DLM 2017 | 6576 | 86 | 4 | 38 | 110 | 25 | 3113 | 6703 |
| GEL-M Day 0 | 11548 | 360 | 4 | 46 | 1235 | 1326 | 16433 | 5167 |
| GEM 2015 | 11548 | 360 | 4 | 46 | 1235 | 1326 | 16433 | 5167 |
| GEM 2016 | 11548 | 360 | 4 | 46 | 1235 | 1326 | 16433 | 5167 |
| GEM 2017 | 5053 | 167 | 5 | 29 | 572 | 674 | 2072 | 5238 |
| LCL-M Day 0 | 17540 | 277 | 5 | 15 | 817 | 2664 | 20400 | 6571 |
| LCM 2015 | 17540 | 277 | 5 | 15 | 817 | 2664 | 20400 | 6571 |
| LCM 2016 | 17540 | 277 | 5 | 15 | 817 | 2664 | 20400 | 6571 |
| LCM 2017 | 19734 | 269 | 9 | 25 | 809 | 3009 | 5075 | 16448 |

| | | | | | | | | |
|---|---|---|---|---|---|---|---|---|
| PL-M Day 0 | 7967 | 58 | 5 | 6 | 302 | 46 | 11883 | 3558 |
| PLM 2015 | 7967 | 58 | 5 | 6 | 302 | 46 | 11883 | 3558 |
| PLM 2016 | 7967 | 58 | 5 | 6 | 302 | 46 | 11883 | 3558 |
| PLM 2017 | 6575 | 88 | 4 | 38 | 110 | 20 | 3100 | 6727 |
| ML16S_10m | 27300 | 1460 | 4 | 37 | 17300 | 9880 | 30400 | n.d. |
| MLW_0517_00 | 27300 | 1460 | 4 | 37 | 17300 | 9880 | 30400 | n.d. |
| MLW_0517_10 | 27300 | 1460 | 4 | 37 | 17300 | 9880 | 30400 | n.d. |
| MLW_0617_00 | 27300 | 1460 | 4 | 37 | 17300 | 9880 | 30400 | n.d. |
| MLW_0917_00_1;MLW_0917_00_2;MLW_0917_00_3 | 27300 | 1460 | 4 | 37 | 17300 | 9880 | 30400 | n.d. |
| MLW_0917_05_1;MLW_0917_05_2;MLW_0917_05_3 | 27300 | 1460 | 4 | 37 | 17300 | 9880 | 30400 | n.d. |
| MLW_0917_10_1;MLW_0917_10_2;MLW_0917_10_3 | 27300 | 1460 | 4 | 37 | 17300 | 9880 | 30400 | n.d. |
| MLW_1018_00 | 27300 | 1460 | 4 | 37 | 17300 | 9880 | 30400 | n.d. |
| MLW_1018_05 | 27300 | 1460 | 4 | 37 | 17300 | 9880 | 30400 | n.d. |
| MLW_1018_12 | 27300 | 1460 | 4 | 37 | 17300 | 9880 | 30400 | n.d. |

n.d. - not determined

*defined according to Boros & Kolpakova 2018

| Sample ID | $Na^+$ (e%) | $K^+$ (e%) | $Ca^{2+}$ (e%) | $Mg^{2+}$ (e%) | $Cl^-$ (e%) | $SO_4^{2-}$ (e%) | $HCO_3^-$ (e%) | $CO_3^{2-}$ (e%) | $HCO_3^-$+$CO_3^{2-}$ (e%) | Ion Balance | Reference for ionic composition |
|---|---|---|---|---|---|---|---|---|---|---|---|
| Bac-12S | 92.8 | 7.0 | 0.1 | 0.1 | 6.8 | 1.9 | 59.8 | 31.5 | 91.3 | 1 | Fazi et al., 2021 |
| Bac-18E | 95.9 | 4.1 | 0.0 | 0.0 | 31.0 | 1.7 | 59.3 | 8.1 | 67.4 | -3 | Melack & Kilham, 1974 (Lameck et al., 2023) |
| Bac-6M | 98.8 | 1.0 | 0.0 | 0.2 | 26.0 | 0.4 | 17.5 | 56.1 | 73.7 | 0 | Getenet et al., 2022 |
| B1-W | 99.0 | 0.9 | 0.0 | 0.0 | 12.9 | 0.3 | 49.9 | 36.9 | 86.8 | 0 | Jirsa et al., 2013 |
| B2-W | 99.0 | 0.9 | 0.0 | 0.0 | 12.9 | 0.3 | 49.9 | 36.9 | 86.8 | 0 | Jirsa et al., 2013 |
| KZ02 | 80.3 | 0.5 | 8.4 | 10.9 | 38.4 | 60.3 | 1.0 | 0.3 | 1.3 | -5 | Boros & Kolpakova 2018 |
| K03bact | 95.3 | 1.7 | 0.0 | 3.0 | 84.9 | 12.6 | 2.2 | 0.3 | 2.5 | -5 | Boros & Kolpakova 2018 |
| KZ04 | 90.4 | 0.4 | 0.5 | 8.6 | 91.2 | 8.5 | 0.2 | 0.0 | 0.2 | 4 | Boros & Kolpakova 2018 |
| KZ05 | 24.8 | 11.0 | 37.4 | 26.8 | 38.0 | 59.2 | 2.8 | 0.0 | 2.8 | 5 | this study |
| K06bact | 80.7 | 0.5 | 1.4 | 17.4 | 84.9 | 14.9 | 0.2 | 0.0 | 0.2 | -6 | Boros & Kolpakova 2018 |
| KZ08 | 81.8 | 1.3 | 1.8 | 15.1 | 67.6 | 32.2 | 0.2 | 0.0 | 0.2 | -3 | Boros & Kolpakova 2018 |
| K10bact | 77.3 | 1.1 | 2.2 | 19.3 | 85.6 | 11.6 | 2.2 | 0.7 | 2.9 | -4 | Boros & Kolpakova 2018 |
| KZ12 | 73.7 | 0.5 | 5.6 | 20.1 | 33.0 | 65.4 | 1.6 | 0.0 | 1.6 | -3 | Boros & Kolpakova 2018 |
| K13bact | 70.5 | 0.9 | 11.7 | 16.8 | 72.1 | 27.1 | 0.8 | 0.0 | 0.8 | 17 | Boros & Kolpakova 2018 |
| K15bact | 53.0 | 0.2 | 5.2 | 41.6 | 89.7 | 10.2 | 0.1 | 0.0 | 0.1 | 9 | Boros & Kolpakova 2018 |
| K16bact | 52.6 | 1.0 | 4.9 | 41.5 | 84.2 | 13.2 | 2.5 | 0.0 | 2.5 | 8 | Boros & Kolpakova 2018 |
| K18bact | 38.1 | 3.0 | 42.8 | 16.0 | 53.0 | 40.8 | 6.2 | 0.0 | 6.2 | -8 | this study |
| KZ19 | 81.0 | 0.5 | 1.1 | 17.4 | 74.1 | 25.7 | 0.2 | 0.0 | 0.2 | -3 | Boros & Kolpakova 2018 |
| K20bact | 23.8 | 5.1 | 55.7 | 15.4 | 22.4 | 45.8 | 31.7 | 0.0 | 31.7 | 3 | this study |
| KT01 | 63.0 | 3.2 | 1.6 | 32.2 | 38.2 | 44.5 | 14.3 | 3.0 | 17.3 | 0 | this study |
| KT02 | 83.1 | 0.7 | 0.1 | 16.1 | 67.6 | 27.8 | 3.4 | 1.2 | 4.6 | -2 | this study |
| KT03 | 63.4 | 1.5 | 1.5 | 33.6 | 41.4 | 39.5 | 14.8 | 4.3 | 19.1 | -1 | this study |
| KT04 | 64.7 | 2.0 | 2.4 | 30.9 | 39.5 | 42.8 | 13.9 | 3.8 | 17.7 | 0 | this study |
| KT05 | 59.8 | 0.6 | 1.2 | 38.4 | 47.7 | 33.8 | 13.8 | 4.8 | 18.6 | -3 | this study |
| KT06 | 64.0 | 2.9 | 1.6 | 31.5 | 40.8 | 41.2 | 12.4 | 5.6 | 18.0 | -2 | this study |
| KT07 | 61.7 | 4.1 | 1.7 | 32.4 | 39.2 | 43.6 | 14.0 | 3.2 | 17.2 | 0 | this study |
| KT08 | 63.4 | 4.2 | 1.3 | 31.1 | 37.6 | 46.1 | 13.3 | 3.0 | 16.3 | -2 | this study |
| KT09 | 41.8 | 1.1 | 6.9 | 50.2 | 72.1 | 26.6 | 1.0 | 0.3 | 1.3 | 3 | this study |
| KT10 | 95.8 | 0.9 | 0.2 | 3.2 | 27.7 | 51.8 | 13.5 | 7.1 | 20.6 | 2 | this study |
| KT11 | 71.1 | 1.0 | 4.8 | 23.1 | 45.9 | 29.3 | 20.5 | 4.3 | 24.8 | 5 | this study |
| KT12 | 94.5 | 0.9 | 0.1 | 4.5 | 21.3 | 42.1 | 23.0 | 13.6 | 36.6 | 3 | this study |
| KT13 | 98.6 | 0.9 | 0.1 | 0.4 | 5.9 | 22.0 | 55.3 | 16.8 | 72.1 | 1 | this study |
| KT14 | 91.3 | 1.1 | 0.1 | 7.5 | 68.2 | 30.0 | 1.4 | 0.4 | 1.8 | -1 | this study |
| KT15 | 87.3 | 1.3 | 0.1 | 11.2 | 46.4 | 52.8 | 0.6 | 0.3 | 0.9 | 2 | this study |
| KT16 | 76.0 | 3.8 | 17.4 | 2.7 | 39.6 | 44.2 | 11.5 | 4.7 | 16.2 | 0 | this study |
| KT17 | 76.3 | 1.6 | 0.4 | 21.7 | 32.8 | 65.9 | 1.0 | 0.4 | 1.4 | 0 | this study |
| KT18 | 76.7 | 2.4 | 0.7 | 20.1 | 26.0 | 25.9 | 6.5 | 41.6 | 48.1 | 2 | this study |
| KT19 | 80.0 | 3.0 | 3.4 | 13.6 | 35.2 | 41.5 | 13.5 | 9.8 | 23.3 | 0 | this study |
| KT20 | 56.4 | 4.2 | 5.4 | 34.0 | 28.3 | 30.6 | 31.5 | 9.6 | 41.1 | -2 | this study |
| KT21 | 78.3 | 0.9 | 8.5 | 12.3 | 74.3 | 6.2 | 19.5 | 0.0 | 19.5 | -1 | this study |
| KT22 | 58.7 | 1.9 | 1.2 | 38.1 | 5.9 | 62.6 | 25.7 | 5.9 | 31.6 | -3 | this study |
| KT23 | 75.4 | 1.1 | 2.3 | 21.2 | 34.3 | 46.0 | 12.8 | 6.8 | 19.6 | 1 | this study |

| | | | | | | | | | | | |
|---|---|---|---|---|---|---|---|---|---|---|---|
| KT24 | 73.8 | 1.1 | 1.6 | 23.5 | 38.6 | 40.6 | 16.9 | 3.9 | 20.8 | 2 | this study |
| KT25 | 77.4 | 0.9 | 0.4 | 21.3 | 34.6 | 46.7 | 13.1 | 5.5 | 18.6 | 0 | this study |
| KT26 | 78.4 | 0.9 | 1.3 | 19.4 | 34.4 | 47.4 | 14.1 | 4.0 | 18.1 | 6 | this study |
| KT27 | 75.6 | 1.3 | 0.5 | 22.6 | 39.3 | 40.8 | 13.9 | 6.1 | 20.0 | -2 | this study |
| 0S | n.d. | n.d. | n.d. | n.d. | n.d. | n.d. | n.d. | n.d. | n.d. | n.d. | |
| 02S | n.d. | n.d. | n.d. | n.d. | n.d. | n.d. | n.d. | n.d. | n.d. | n.d. | |
| 04S | n.d. | n.d. | n.d. | n.d. | n.d. | n.d. | n.d. | n.d. | n.d. | n.d. | |
| 06S | n.d. | n.d. | n.d. | n.d. | n.d. | n.d. | n.d. | n.d. | n.d. | n.d. | |
| 08S | n.d. | n.d. | n.d. | n.d. | n.d. | n.d. | n.d. | n.d. | n.d. | n.d. | |
| 09S | n.d. | n.d. | n.d. | n.d. | n.d. | n.d. | n.d. | n.d. | n.d. | n.d. | |
| 11S | n.d. | n.d. | n.d. | n.d. | n.d. | n.d. | n.d. | n.d. | n.d. | n.d. | |
| SHV | 63.3 | 1.3 | 25.3 | 10.1 | 42.0 | 19.0 | 36.0 | 3.0 | 39.0 | 19 | this study |
| KA1 | 93.6 | 0.1 | 0.6 | 5.6 | 55.5 | 43.5 | 0.1 | 0.8 | 0.9 | -1 | this study |
| KA2 | 93.1 | 0.5 | 0.9 | 5.5 | 66.4 | 12.6 | 17.9 | 3.1 | 21.0 | 1 | this study |
| KA3 | 90.4 | 0.4 | 0.5 | 8.7 | 66.6 | 29.7 | 2.2 | 1.5 | 3.7 | -1 | this study |
| KA4 | 89.7 | 1.0 | 1.5 | 7.8 | 0.2 | 98.8 | 0.7 | 0.4 | 1.1 | 29 | this study |
| KA5 | 35.4 | 0.9 | 7.8 | 55.9 | 38.9 | 22.2 | 38.9 | 0.0 | 38.9 | 27 | this study |
| KA6 | 92.2 | 0.1 | 0.6 | 7.1 | 72.3 | 27.3 | 0.4 | 0.0 | 0.4 | -2 | this study |
| KA7 | 52.5 | 0.6 | 16.2 | 30.7 | 68.6 | 30.4 | 1.0 | 0.0 | 1.0 | -2 | this study |
| KA8 | 42.2 | 0.7 | 14.3 | 42.8 | 21.0 | 69.2 | 9.8 | 0.0 | 9.8 | 10 | this study |
| Gudzh_water | 97.6 | n.d. | 0.1 | 2.3 | 12.8 | 73.9 | 10.2 | 3.2 | 13.4 | ? | Lavrentyeva et al., 2020 |
| NB30 | 98.7 | 0.5 | 0.2 | 0.6 | 25.3 | 0.7 | 28.6 | 45.4 | 74.0 | -13 | Gorlenko et al., 2010 |
| NB31 | 98.7 | 0.5 | 0.2 | 0.6 | 25.3 | 0.7 | 28.6 | 45.4 | 74.0 | -13 | Gorlenko et al., 2010 |
| NB32 | 98.7 | 0.5 | 0.2 | 0.6 | 25.3 | 0.7 | 28.6 | 45.4 | 74.0 | -13 | Gorlenko et al., 2010 |
| NB41 | 98.7 | 0.5 | 0.2 | 0.6 | 25.3 | 0.7 | 28.6 | 45.4 | 74.0 | -13 | Gorlenko et al., 2010 |
| NB42 | 98.7 | 0.5 | 0.2 | 0.6 | 25.3 | 0.7 | 28.6 | 45.4 | 74.0 | -13 | Gorlenko et al., 2010 |
| NB43 | 98.7 | 0.5 | 0.2 | 0.6 | 25.3 | 0.7 | 28.6 | 45.4 | 74.0 | -13 | Gorlenko et al., 2010 |
| Bangong Co | n.d. | n.d. | n.d. | n.d. | n.d. | n.d. | n.d. | n.d. | n.d. | n.d. | Lin et al., 2021 |
| Bong Co | 31.4 | 14.9 | 26.3 | 27.4 | 11.0 | 65.1 | 21.1 | 2.8 | 23.9 | 0 | Yuan et al., 2011 |
| Co Ngoin (1) | 23.3 | 0.0 | 17.8 | 58.8 | 5.3 | 7.9 | 86.8 | n.d. | 86.8 | 3 | Sun et al., 2020 |
| Mapam Yumco | 35.5 | 2.4 | 22.0 | 40.1 | 6.2 | 10.5 | 83.3 | n.d. | 83.3 | 0 | Yao et al., 2015 |
| Urru Co | n.d. | n.d. | n.d. | n.d. | n.d. | n.d. | n.d. | n.d. | n.d. | n.d. | not found |
| Bam Co | 80.1 | 5.4 | 2.8 | 11.7 | 28.0 | 63.6 | 5.8 | 2.7 | 8.5 | 0 | Yuan et al., 2011 |
| Bero Zeco | 74.0 | 4.1 | 0.0 | 21.9 | 7.5 | 90.3 | 2.2 | 0.0 | 2.2 | 11 | Li et al., 2019 |
| Co Ngoin (2) | 82.4 | 4.8 | 0.4 | 12.4 | 26.9 | 28.8 | 19.5 | 24.7 | 44.3 | -4 | Xiong et al., 2020 |
| Dawa Co | 81.8 | 6.1 | 0.7 | 11.3 | 22.7 | 75.3 | 1.1 | 1.0 | 2.1 | 0 | Yuan et al., 2011 |
| Tangra Yumco | 71.1 | 0.0 | 0.0 | 11.5 | 24.5 | 18.6 | 38.2 | n.d. | 38.2 | n.d. | Qiao et al., 2017; Günther et al., 2014 |
| Dong Co | 77.1 | 13.7 | 0.5 | 8.7 | 33.5 | 65.9 | 0.5 | 0.0 | 0.5 | 0 | Yuan et al., 2011; Li et al., 2019; Sun et al., 2020 |
| Kunggyu | 71.7 | 9.0 | 8.8 | 10.6 | 26.0 | 1.0 | 32.0 | 40.9 | 72.9 | -11 | Günther et al., 2014; Shen et al., 2022 |
| Nam Co | 80.9 | 0.0 | 1.2 | 17.9 | 5.6 | 14.1 | 54.5 | 25.8 | 80.3 | 2 | Williams, 1991, Wang et al., 2010 |
| Pung Co | 79.2 | 5.8 | 2.4 | 12.6 | 15.4 | 52.7 | 22.6 | 9.4 | 32.0 | 0 | Yuan et al., 2011 |
| Selin Co | 86.9 | 5.3 | 0.0 | 7.7 | 34.4 | 52.2 | 8.5 | 4.9 | 13.4 | 0 | Williams, 1991 |
| Zhari Namco | 84.8 | 4.4 | 1.4 | 9.3 | 16.9 | 77.5 | 3.2 | 2.4 | 5.6 | 0 | Yuan et al., 2011 |
| Zhaxi Co | 79.5 | 9.1 | 1.0 | 10.4 | 21.5 | 72.7 | 3.4 | 2.5 | 5.9 | 0 | Yuan et al., 2011 |

| | | | | | | | | | | | |
|---|---|---|---|---|---|---|---|---|---|---|---|
| Bangkog Co | 85.1 | 7.6 | 0.7 | 6.7 | 42.0 | 55.5 | 1.5 | 1.0 | 2.5 | 0 | Yuan et al., 2011 |
| Dangqiong Co | 92.9 | 7.1 | 0.0 | 0.0 | 78.9 | 6.0 | 0.0 | 15.1 | 15.1 | 2 | Li et al., 2019 |
| Nyer Co (Nieer Co) | 49.8 | 12.5 | 0.0 | 37.7 | 74.4 | 25.6 | 0.0 | 0.0 | 0.0 | 0 | Li et al., 2019 |
| BU1 | 96.9 | 0.5 | 1.0 | 1.6 | 20.8 | 5.5 | 58.3 | 15.4 | 73.7 | 5 | Boros & Kolpakova 2018 |
| SO1 | 97.3 | 0.0 | 0.9 | 1.8 | 35.8 | 8.2 | 49.9 | 6.1 | 56.0 | 1 | this study |
| ZA1 | 97.0 | 0.4 | 1.0 | 1.6 | 26.3 | 6.9 | 48.4 | 18.4 | 66.8 | 0 | Boros & Kolpakova 2018 |
| SO2 | 97.3 | 0.0 | 0.9 | 1.8 | 35.8 | 8.2 | 49.9 | 6.1 | 56.0 | 1 | this study |
| SO3 | 97.3 | 0.0 | 0.9 | 1.8 | 35.8 | 8.2 | 49.9 | 6.1 | 56.0 | 1 | this study |
| SO4 | 97.3 | 0.0 | 0.9 | 1.8 | 35.8 | 8.2 | 49.9 | 6.1 | 56.0 | 1 | this study |
| SO5 | 97.3 | 0.0 | 0.9 | 1.8 | 35.8 | 8.2 | 49.9 | 6.1 | 56.0 | 1 | this study |
| SO6 | 97.3 | 0.0 | 0.9 | 1.8 | 35.8 | 8.2 | 49.9 | 6.1 | 56.0 | 1 | this study |
| SO9 | 97.3 | 0.0 | 0.9 | 1.8 | 35.8 | 8.2 | 49.9 | 6.1 | 56.0 | 1 | this study |
| SO11 | 97.3 | 0.0 | 0.9 | 1.8 | 35.8 | 8.2 | 49.9 | 6.1 | 56.0 | 1 | this study |
| SO12 | 97.3 | 0.0 | 0.9 | 1.8 | 35.8 | 8.2 | 49.9 | 6.1 | 56.0 | 1 | this study |
| SO13 | 97.3 | 0.0 | 0.9 | 1.8 | 35.8 | 8.2 | 49.9 | 6.1 | 56.0 | 1 | this study |
| SO14 | 97.3 | 0.0 | 0.9 | 1.8 | 35.8 | 8.2 | 49.9 | 6.1 | 56.0 | 1 | this study |
| SO15 | 97.3 | 0.0 | 0.9 | 1.8 | 35.8 | 8.2 | 49.9 | 6.1 | 56.0 | 1 | this study |
| SO16 | 97.3 | 0.0 | 0.9 | 1.8 | 35.8 | 8.2 | 49.9 | 6.1 | 56.0 | 1 | this study |
| VBK | 97.4 | 0.4 | 0.7 | 1.5 | 30.6 | 9.7 | 56.8 | 2.9 | 59.7 | -3 | Korponai et al., 2019 |
| ZA2 | 97.0 | 0.4 | 1.0 | 1.6 | 26.3 | 6.9 | 48.4 | 18.4 | 66.8 | 0 | Boros & Kolpakova 2018 |
| ZA3 | 97.0 | 0.4 | 1.0 | 1.6 | 26.3 | 6.9 | 48.4 | 18.4 | 66.8 | 0 | Boros & Kolpakova 2018 |
| ZA4 | 97.0 | 0.4 | 1.0 | 1.6 | 26.3 | 6.9 | 48.4 | 18.4 | 66.8 | 0 | Boros & Kolpakova 2018 |
| ZA5 | 97.0 | 0.4 | 1.0 | 1.6 | 26.3 | 6.9 | 48.4 | 18.4 | 66.8 | 0 | Boros & Kolpakova 2018 |
| ZA6 | 97.0 | 0.4 | 1.0 | 1.6 | 26.3 | 6.9 | 48.4 | 18.4 | 66.8 | 0 | Boros & Kolpakova 2018 |
| ZA7 | 97.0 | 0.4 | 1.0 | 1.6 | 26.3 | 6.9 | 48.4 | 18.4 | 66.8 | 0 | Boros & Kolpakova 2018 |
| ZA8 | 97.0 | 0.4 | 1.0 | 1.6 | 26.3 | 6.9 | 48.4 | 18.4 | 66.8 | 0 | Boros & Kolpakova 2018 |
| ZA9 | 97.0 | 0.4 | 1.0 | 1.6 | 26.3 | 6.9 | 48.4 | 18.4 | 66.8 | 0 | Boros & Kolpakova 2018 |
| ZA10 | 97.0 | 0.4 | 1.0 | 1.6 | 26.3 | 6.9 | 48.4 | 18.4 | 66.8 | 0 | Boros & Kolpakova 2018 |
| ZA11 | 97.0 | 0.4 | 1.0 | 1.6 | 26.3 | 6.9 | 48.4 | 18.4 | 66.8 | 0 | Boros & Kolpakova 2018 |
| ZA12 | 97.0 | 0.4 | 1.0 | 1.6 | 26.3 | 6.9 | 48.4 | 18.4 | 66.8 | 0 | Boros & Kolpakova 2018 |
| ZA13 | 97.0 | 0.4 | 1.0 | 1.6 | 26.3 | 6.9 | 48.4 | 18.4 | 66.8 | 0 | Boros & Kolpakova 2018 |
| ZA14 | 97.0 | 0.4 | 1.0 | 1.6 | 26.3 | 6.9 | 48.4 | 18.4 | 66.8 | 0 | Boros & Kolpakova 2018 |
| ZA15 | 97.0 | 0.4 | 1.0 | 1.6 | 26.3 | 6.9 | 48.4 | 18.4 | 66.8 | 0 | Boros & Kolpakova 2018 |
| ZA16 | 97.0 | 0.4 | 1.0 | 1.6 | 26.3 | 6.9 | 48.4 | 18.4 | 66.8 | 0 | Boros & Kolpakova 2018 |
| ZA17 | 97.0 | 0.4 | 1.0 | 1.6 | 26.3 | 6.9 | 48.4 | 18.4 | 66.8 | 0 | Boros & Kolpakova 2018 |
| M05 | 99.3 | 0.1 | 0.5 | 0.1 | 98.5 | 0.4 | 0.5 | 0.6 | 1.1 | -1 | Andrei et al., 2015 |
| U05 | n.d. | n.d. | n.d. | n.d. | n.d. | n.d. | n.d. | n.d. | n.d. | n.d. | this study |
| U15 | n.d. | n.d. | n.d. | n.d. | n.d. | n.d. | n.d. | n.d. | n.d. | n.d. | this study |
| U28 | n.d. | n.d. | n.d. | n.d. | n.d. | n.d. | n.d. | n.d. | n.d. | n.d. | this study |
| B01w | 56.1 | 2.6 | 2.4 | 38.9 | 27.5 | 32.3 | 37.3 | 2.9 | 40.2 | -1 | this study |
| B02w | 56.1 | 2.6 | 2.4 | 38.9 | 27.5 | 32.3 | 37.3 | 2.9 | 40.2 | -1 | this study |
| B03w | 56.1 | 2.6 | 2.4 | 38.9 | 27.5 | 32.3 | 37.3 | 2.9 | 40.2 | -1 | this study |
| B04w | 56.1 | 2.6 | 2.4 | 38.9 | 27.5 | 32.3 | 37.3 | 2.9 | 40.2 | -1 | this study |
| B05w | 56.1 | 2.6 | 2.4 | 38.9 | 27.5 | 32.3 | 37.3 | 2.9 | 40.2 | -1 | this study |

| | | | | | | | | | | | |
|---|---|---|---|---|---|---|---|---|---|---|---|
| B06w | 56.1 | 2.6 | 2.4 | 38.9 | 27.5 | 32.3 | 37.3 | 2.9 | 40.2 | -1 | this study |
| B07w | 56.1 | 2.6 | 2.4 | 38.9 | 27.5 | 32.3 | 37.3 | 2.9 | 40.2 | -1 | this study |
| B08w | 56.1 | 2.6 | 2.4 | 38.9 | 27.5 | 32.3 | 37.3 | 2.9 | 40.2 | -1 | this study |
| B09w | 56.1 | 2.6 | 2.4 | 38.9 | 27.5 | 32.3 | 37.3 | 2.9 | 40.2 | -1 | this study |
| B010w | 56.1 | 2.6 | 2.4 | 38.9 | 27.5 | 32.3 | 37.3 | 2.9 | 40.2 | -1 | this study |
| B011w | 56.1 | 2.6 | 2.4 | 38.9 | 27.5 | 32.3 | 37.3 | 2.9 | 40.2 | -1 | this study |
| KH1w | 54.8 | 2.7 | 3.9 | 38.7 | 26.3 | 30.8 | 37.3 | 5.7 | 43.0 | 0 | this study |
| KH2w | 54.8 | 2.7 | 3.9 | 38.7 | 26.3 | 30.8 | 37.3 | 5.7 | 43.0 | 0 | this study |
| KH3w | 54.8 | 2.7 | 3.9 | 38.7 | 26.3 | 30.8 | 37.3 | 5.7 | 43.0 | 0 | this study |
| KH4w | 54.8 | 2.7 | 3.9 | 38.7 | 26.3 | 30.8 | 37.3 | 5.7 | 43.0 | 0 | this study |
| KH5w | 54.8 | 2.7 | 3.9 | 38.7 | 26.3 | 30.8 | 37.3 | 5.7 | 43.0 | 0 | this study |
| KH6w | 54.8 | 2.7 | 3.9 | 38.7 | 26.3 | 30.8 | 37.3 | 5.7 | 43.0 | 0 | this study |
| KH7w | 54.8 | 2.7 | 3.9 | 38.7 | 26.3 | 30.8 | 37.3 | 5.7 | 43.0 | 0 | this study |
| KH8w | 54.8 | 2.7 | 3.9 | 38.7 | 26.3 | 30.8 | 37.3 | 5.7 | 43.0 | 0 | this study |
| KH9w | 54.8 | 2.7 | 3.9 | 38.7 | 26.3 | 30.8 | 37.3 | 5.7 | 43.0 | 0 | this study |
| KH10w | 54.8 | 2.7 | 3.9 | 38.7 | 26.3 | 30.8 | 37.3 | 5.7 | 43.0 | 0 | this study |
| KH11w | 54.8 | 2.7 | 3.9 | 38.7 | 26.3 | 30.8 | 37.3 | 5.7 | 43.0 | 0 | this study |
| B01 | 96.5 | 0.3 | 1.1 | 2.1 | 46.5 | 8.7 | 32.3 | 12.4 | 44.7 | 3 | Boros & Kolpakova 2018 |
| B02 | 96.5 | 0.3 | 1.1 | 2.1 | 46.5 | 8.7 | 32.3 | 12.4 | 44.7 | 3 | Boros & Kolpakova 2018 |
| B03 | 96.5 | 0.3 | 1.1 | 2.1 | 46.5 | 8.7 | 32.3 | 12.4 | 44.7 | 3 | Boros & Kolpakova 2018 |
| B04 | 96.5 | 0.3 | 1.1 | 2.1 | 46.5 | 8.7 | 32.3 | 12.4 | 44.7 | 3 | Boros & Kolpakova 2018 |
| B05 | 96.5 | 0.3 | 1.1 | 2.1 | 46.5 | 8.7 | 32.3 | 12.4 | 44.7 | 3 | Boros & Kolpakova 2018 |
| B06 | 96.5 | 0.3 | 1.1 | 2.1 | 46.5 | 8.7 | 32.3 | 12.4 | 44.7 | 3 | Boros & Kolpakova 2018 |
| B07 | 96.5 | 0.3 | 1.1 | 2.1 | 46.5 | 8.7 | 32.3 | 12.4 | 44.7 | 3 | Boros & Kolpakova 2018 |
| B08 | 96.5 | 0.3 | 1.1 | 2.1 | 46.5 | 8.7 | 32.3 | 12.4 | 44.7 | 3 | Boros & Kolpakova 2018 |
| B09 | 96.5 | 0.3 | 1.1 | 2.1 | 46.5 | 8.7 | 32.3 | 12.4 | 44.7 | 3 | Boros & Kolpakova 2018 |
| B10 | 96.5 | 0.3 | 1.1 | 2.1 | 46.5 | 8.7 | 32.3 | 12.4 | 44.7 | 3 | Boros & Kolpakova 2018 |
| B11 | 96.5 | 0.3 | 1.1 | 2.1 | 46.5 | 8.7 | 32.3 | 12.4 | 44.7 | 3 | Boros & Kolpakova 2018 |
| B12 | 96.5 | 0.3 | 1.1 | 2.1 | 46.5 | 8.7 | 32.3 | 12.4 | 44.7 | 3 | Boros & Kolpakova 2018 |
| B13 | 96.5 | 0.3 | 1.1 | 2.1 | 46.5 | 8.7 | 32.3 | 12.4 | 44.7 | 3 | Boros & Kolpakova 2018 |
| B14 | 96.5 | 0.3 | 1.1 | 2.1 | 46.5 | 8.7 | 32.3 | 12.4 | 44.7 | 3 | Boros & Kolpakova 2018 |
| K01 | 95.6 | 0.6 | 1.4 | 2.4 | 35.5 | 7.1 | 39.1 | 18.3 | 57.4 | 0 | Boros & Kolpakova 2018 |
| K02 | 95.6 | 0.6 | 1.4 | 2.4 | 35.5 | 7.1 | 39.1 | 18.3 | 57.4 | 0 | Boros & Kolpakova 2018 |
| K03 | 95.6 | 0.6 | 1.4 | 2.4 | 35.5 | 7.1 | 39.1 | 18.3 | 57.4 | 0 | Boros & Kolpakova 2018 |
| K04 | 95.6 | 0.6 | 1.4 | 2.4 | 35.5 | 7.1 | 39.1 | 18.3 | 57.4 | 0 | Boros & Kolpakova 2018 |
| K05 | 95.6 | 0.6 | 1.4 | 2.4 | 35.5 | 7.1 | 39.1 | 18.3 | 57.4 | 0 | Boros & Kolpakova 2018 |
| K06 | 95.6 | 0.6 | 1.4 | 2.4 | 35.5 | 7.1 | 39.1 | 18.3 | 57.4 | 0 | Boros & Kolpakova 2018 |
| K09 | 95.6 | 0.6 | 1.4 | 2.4 | 35.5 | 7.1 | 39.1 | 18.3 | 57.4 | 0 | Boros & Kolpakova 2018 |
| K12 | 95.6 | 0.6 | 1.4 | 2.4 | 35.5 | 7.1 | 39.1 | 18.3 | 57.4 | 0 | Boros & Kolpakova 2018 |
| K14 | 95.6 | 0.6 | 1.4 | 2.4 | 35.5 | 7.1 | 39.1 | 18.3 | 57.4 | 0 | Boros & Kolpakova 2018 |
| S01 | 97.3 | 0.0 | 0.9 | 1.8 | 35.8 | 8.2 | 49.9 | 6.1 | 56.0 | 1 | this study |
| S02 | 97.3 | 0.0 | 0.9 | 1.8 | 35.8 | 8.2 | 49.9 | 6.1 | 56.0 | 1 | this study |
| S03 | 97.3 | 0.0 | 0.9 | 1.8 | 35.8 | 8.2 | 49.9 | 6.1 | 56.0 | 1 | this study |
| S04 | 97.3 | 0.0 | 0.9 | 1.8 | 35.8 | 8.2 | 49.9 | 6.1 | 56.0 | 1 | this study |

| | | | | | | | | | | | |
|---|---|---|---|---|---|---|---|---|---|---|---|
| S05 | 97.3 | 0.0 | 0.9 | 1.8 | 35.8 | 8.2 | 49.9 | 6.1 | 56.0 | 1 | this study |
| S06 | 97.3 | 0.0 | 0.9 | 1.8 | 35.8 | 8.2 | 49.9 | 6.1 | 56.0 | 1 | this study |
| S07 | 97.3 | 0.0 | 0.9 | 1.8 | 35.8 | 8.2 | 49.9 | 6.1 | 56.0 | 1 | this study |
| S08 | 97.3 | 0.0 | 0.9 | 1.8 | 35.8 | 8.2 | 49.9 | 6.1 | 56.0 | 1 | this study |
| S09 | 97.3 | 0.0 | 0.9 | 1.8 | 35.8 | 8.2 | 49.9 | 6.1 | 56.0 | 1 | this study |
| S10 | 97.3 | 0.0 | 0.9 | 1.8 | 35.8 | 8.2 | 49.9 | 6.1 | 56.0 | 1 | this study |
| S11 | 97.3 | 0.0 | 0.9 | 1.8 | 35.8 | 8.2 | 49.9 | 6.1 | 56.0 | 1 | this study |
| S12 | 97.3 | 0.0 | 0.9 | 1.8 | 35.8 | 8.2 | 49.9 | 6.1 | 56.0 | 1 | this study |
| S13 | 97.3 | 0.0 | 0.9 | 1.8 | 35.8 | 8.2 | 49.9 | 6.1 | 56.0 | 1 | this study |
| S14 | 97.3 | 0.0 | 0.9 | 1.8 | 35.8 | 8.2 | 49.9 | 6.1 | 56.0 | 1 | this study |
| V01 | 97.4 | 0.4 | 0.7 | 1.5 | 30.6 | 9.7 | 56.8 | 2.9 | 59.7 | -3 | Korponai et al., 2020 |
| V02 | 97.4 | 0.4 | 0.7 | 1.5 | 30.6 | 9.7 | 56.8 | 2.9 | 59.7 | -3 | Korponai et al., 2020 |
| V03 | 97.4 | 0.4 | 0.7 | 1.5 | 30.6 | 9.7 | 56.8 | 2.9 | 59.7 | -3 | Korponai et al., 2020 |
| V04 | 97.4 | 0.4 | 0.7 | 1.5 | 30.6 | 9.7 | 56.8 | 2.9 | 59.7 | -3 | Korponai et al., 2020 |
| V05 | 97.4 | 0.4 | 0.7 | 1.5 | 30.6 | 9.7 | 56.8 | 2.9 | 59.7 | -3 | Korponai et al., 2020 |
| V06 | 97.4 | 0.4 | 0.7 | 1.5 | 30.6 | 9.7 | 56.8 | 2.9 | 59.7 | -3 | Korponai et al., 2020 |
| V07 | 97.4 | 0.4 | 0.7 | 1.5 | 30.6 | 9.7 | 56.8 | 2.9 | 59.7 | -3 | Korponai et al., 2020 |
| V08 | 97.4 | 0.4 | 0.7 | 1.5 | 30.6 | 9.7 | 56.8 | 2.9 | 59.7 | -3 | Korponai et al., 2020 |
| V09 | 97.4 | 0.4 | 0.7 | 1.5 | 30.6 | 9.7 | 56.8 | 2.9 | 59.7 | -3 | Korponai et al., 2020 |
| V10 | 97.4 | 0.4 | 0.7 | 1.5 | 30.6 | 9.7 | 56.8 | 2.9 | 59.7 | -3 | Korponai et al., 2020 |
| V11 | 97.4 | 0.4 | 0.7 | 1.5 | 30.6 | 9.7 | 56.8 | 2.9 | 59.7 | -3 | Korponai et al., 2020 |
| V12 | 97.4 | 0.4 | 0.7 | 1.5 | 30.6 | 9.7 | 56.8 | 2.9 | 59.7 | -3 | Korponai et al., 2020 |
| V13 | 97.4 | 0.4 | 0.7 | 1.5 | 30.6 | 9.7 | 56.8 | 2.9 | 59.7 | -3 | Korponai et al., 2020 |
| V14 | 97.4 | 0.4 | 0.7 | 1.5 | 30.6 | 9.7 | 56.8 | 2.9 | 59.7 | -3 | Korponai et al., 2020 |
| Z01 | 97.0 | 0.4 | 1.0 | 1.6 | 26.3 | 6.9 | 48.4 | 18.4 | 66.8 | 0 | Boros & Kolpakova 2018 |
| Z02 | 97.0 | 0.4 | 1.0 | 1.6 | 26.3 | 6.9 | 48.4 | 18.4 | 66.8 | 0 | Boros & Kolpakova 2018 |
| Z03 | 97.0 | 0.4 | 1.0 | 1.6 | 26.3 | 6.9 | 48.4 | 18.4 | 66.8 | 0 | Boros & Kolpakova 2018 |
| Z04 | 97.0 | 0.4 | 1.0 | 1.6 | 26.3 | 6.9 | 48.4 | 18.4 | 66.8 | 0 | Boros & Kolpakova 2018 |
| Z05 | 97.0 | 0.4 | 1.0 | 1.6 | 26.3 | 6.9 | 48.4 | 18.4 | 66.8 | 0 | Boros & Kolpakova 2018 |
| Z06 | 97.0 | 0.4 | 1.0 | 1.6 | 26.3 | 6.9 | 48.4 | 18.4 | 66.8 | 0 | Boros & Kolpakova 2018 |
| Z07 | 97.0 | 0.4 | 1.0 | 1.6 | 26.3 | 6.9 | 48.4 | 18.4 | 66.8 | 0 | Boros & Kolpakova 2018 |
| Z08 | 97.0 | 0.4 | 1.0 | 1.6 | 26.3 | 6.9 | 48.4 | 18.4 | 66.8 | 0 | Boros & Kolpakova 2018 |
| Z09 | 97.0 | 0.4 | 1.0 | 1.6 | 26.3 | 6.9 | 48.4 | 18.4 | 66.8 | 0 | Boros & Kolpakova 2018 |
| Z10 | 97.0 | 0.4 | 1.0 | 1.6 | 26.3 | 6.9 | 48.4 | 18.4 | 66.8 | 0 | Boros & Kolpakova 2018 |
| Z12 | 97.0 | 0.4 | 1.0 | 1.6 | 26.3 | 6.9 | 48.4 | 18.4 | 66.8 | 0 | Boros & Kolpakova 2018 |
| Z14 | 97.0 | 0.4 | 1.0 | 1.6 | 26.3 | 6.9 | 48.4 | 18.4 | 66.8 | 0 | Boros & Kolpakova 2018 |
| BV | 17.4 | 2.3 | 30.3 | 50.0 | 10.8 | 30.5 | 52.4 | 6.4 | 58.7 | -1 | Szilágyi2005 |
| K0 | n.d. | n.d. | n.d. | n.d. | n.d. | n.d. | n.d. | n.d. | n.d. | n.d. | not found |
| K01 | n.d. | n.d. | n.d. | n.d. | n.d. | n.d. | n.d. | n.d. | n.d. | n.d. | not found |
| A | n.d. | n.d. | n.d. | n.d. | n.d. | n.d. | n.d. | n.d. | n.d. | n.d. | not found |
| B | n.d. | n.d. | n.d. | n.d. | n.d. | n.d. | n.d. | n.d. | n.d. | n.d. | not found |
| B0 | 56.1 | 2.6 | 2.4 | 38.9 | 27.5 | 32.3 | 37.3 | 2.9 | 40.2 | -1 | this study |
| C | n.d. | n.d. | n.d. | n.d. | n.d. | n.d. | n.d. | n.d. | n.d. | n.d. | not found |
| D | n.d. | n.d. | n.d. | n.d. | n.d. | n.d. | n.d. | n.d. | n.d. | n.d. | not found |

| | | | | | | | | | | | |
|---|---|---|---|---|---|---|---|---|---|---|---|
| Ds | n.d. | n.d. | n.d. | n.d. | n.d. | n.d. | n.d. | n.d. | n.d. | n.d. | not found |
| E | n.d. | n.d. | n.d. | n.d. | n.d. | n.d. | n.d. | n.d. | n.d. | n.d. | not found |
| F | n.d. | n.d. | n.d. | n.d. | n.d. | n.d. | n.d. | n.d. | n.d. | n.d. | not found |
| FDN | n.d. | n.d. | n.d. | n.d. | n.d. | n.d. | n.d. | n.d. | n.d. | n.d. | not found |
| FDO | n.d. | n.d. | n.d. | n.d. | n.d. | n.d. | n.d. | n.d. | n.d. | n.d. | not found |
| Ff | 18.5 | 2.6 | 23.2 | 55.7 | 11.4 | 35.0 | 45.8 | 7.8 | 53.6 | -1 | Szilágyi2005 |
| K | 17.4 | 2.3 | 30.3 | 50.0 | 10.8 | 30.5 | 52.4 | 6.4 | 58.7 | -1 | Szilágyi2005 |
| KH | 54.8 | 2.7 | 3.9 | 38.7 | 26.3 | 30.8 | 37.3 | 5.7 | 43.0 | 0 | this study |
| Kn | n.d. | n.d. | n.d. | n.d. | n.d. | n.d. | n.d. | n.d. | n.d. | n.d. | not found |
| LK | n.d. | n.d. | n.d. | n.d. | n.d. | n.d. | n.d. | n.d. | n.d. | n.d. | not found |
| LN | n.d. | n.d. | n.d. | n.d. | n.d. | n.d. | n.d. | n.d. | n.d. | n.d. | not found |
| NH | 54.8 | 2.7 | 3.9 | 38.7 | 26.3 | 30.8 | 37.3 | 5.7 | 43.0 | 0 | this study |
| R | n.d. | n.d. | n.d. | n.d. | n.d. | n.d. | n.d. | n.d. | n.d. | n.d. | not found |
| TLO | n.d. | n.d. | n.d. | n.d. | n.d. | n.d. | n.d. | n.d. | n.d. | n.d. | not found |
| TNB | n.d. | n.d. | n.d. | n.d. | n.d. | n.d. | n.d. | n.d. | n.d. | n.d. | not found |
| TO | n.d. | n.d. | n.d. | n.d. | n.d. | n.d. | n.d. | n.d. | n.d. | n.d. | not found |
| TS | n.d. | n.d. | n.d. | n.d. | n.d. | n.d. | n.d. | n.d. | n.d. | n.d. | not found |
| Z | 16.2 | 1.5 | 46.7 | 35.6 | 9.9 | 16.8 | 73.2 | 0.0 | 73.2 | -1 | Szilágyi2005 |
| SS15 | 88.4 | 3.0 | 0.4 | 8.2 | 30.0 | 9.6 | 53.9 | 6.5 | 60.4 | -3 | Boros & Kolpakova 2018 |
| SS16 | 88.4 | 3.0 | 0.4 | 8.2 | 30.0 | 9.6 | 53.9 | 6.5 | 60.4 | -3 | Boros & Kolpakova 2018 |
| SS17 | 88.4 | 3.0 | 0.4 | 8.2 | 30.0 | 9.6 | 53.9 | 6.5 | 60.4 | -3 | Boros & Kolpakova 2018 |
| US15 | 77.2 | 1.4 | 2.1 | 19.3 | 12.9 | 5.6 | 72.6 | 8.9 | 81.5 | 1 | Boros & Kolpakova 2018 |
| US16 | 77.2 | 1.4 | 2.1 | 19.3 | 12.9 | 5.6 | 72.6 | 8.9 | 81.5 | 1 | Boros & Kolpakova 2018 |
| US17 | 77.2 | 1.4 | 2.1 | 19.3 | 12.9 | 5.6 | 72.6 | 8.9 | 81.5 | 1 | Boros & Kolpakova 2018 |
| ZL15 | 84.8 | 2.7 | 1.2 | 11.4 | 22.2 | 30.8 | 41.0 | 6.1 | 47.1 | -4 | Boros & Kolpakova 2018 |
| ZL16 | 84.8 | 2.7 | 1.2 | 11.4 | 22.2 | 30.8 | 41.0 | 6.1 | 47.1 | -4 | Boros & Kolpakova 2018 |
| ZL17 | 84.8 | 2.7 | 1.2 | 11.4 | 22.2 | 30.8 | 41.0 | 6.1 | 47.1 | -4 | Boros & Kolpakova 2018 |
| Sp17_01 | 76.3 | 3.1 | 1.5 | 19.1 | 26.5 | 7.8 | 58.3 | 7.4 | 65.7 | 7 | Boros & Kolpakova 2018 |
| Sp17_02 | 95.3 | 0.4 | 1.6 | 2.7 | 6.2 | 10.4 | 72.3 | 11.2 | 83.5 | -5 | Boros & Kolpakova 2018 |
| Sp17_04 | 50.5 | 1.9 | 28.6 | 19.0 | 3.1 | 11.0 | 82.7 | 3.2 | 85.9 | 2 | Boros & Kolpakova 2018 |
| Sp17_05 | 61.7 | 1.9 | 0.0 | 36.4 | 30.5 | 35.7 | 25.7 | 8.0 | 33.7 | 2 | this study |
| Sp17_07 | 90.5 | 0.5 | 0.9 | 8.0 | 13.9 | 14.0 | 64.1 | 8.0 | 72.1 | 4 | Boros & Kolpakova 2018 |
| Sp17_09 | 83.0 | 2.1 | 1.2 | 13.8 | 17.7 | 25.5 | 46.1 | 10.7 | 56.8 | n.d. | Boros et al., 2013 |
| Sp17_10 | 87.9 | 0.6 | 2.5 | 8.9 | 13.7 | 7.3 | 70.6 | 8.5 | 79.1 | 10 | Boros & Kolpakova 2018 |
| Sp17_11 | 71.1 | 2.1 | 4.2 | 22.6 | 12.4 | 26.5 | 50.7 | 10.4 | 61.1 | 4 | Boros & Kolpakova 2018 |
| Sp17_12 | 90.1 | 0.7 | 1.4 | 7.8 | 8.7 | 5.0 | 75.7 | 10.5 | 86.2 | 3 | Boros & Kolpakova 2018 |
| Sp17_13 | 91.7 | 2.1 | 0.2 | 6.1 | 14.6 | 7.4 | 69.1 | 8.9 | 78.0 | -10 | Boros & Kolpakova 2018 |
| Sp17_14 | 69.2 | 1.2 | 2.2 | 24.4 | 14.9 | 11.7 | 70.7 | 2.7 | 73.4 | n.d. | Boros et al., 2013 |
| Sp17_15 | 58.2 | 2.3 | 1.4 | 38.0 | 27.2 | 33.0 | 34.4 | 5.4 | 39.8 | 0 | this study |
| Sp17_16 | 80.6 | 1.2 | 1.7 | 16.5 | 14.4 | 9.1 | 66.7 | 9.8 | 76.5 | 8 | Boros & Kolpakova 2018 |
| Sp17_17 | 93.3 | 2.0 | 0.2 | 4.5 | 15.6 | 7.1 | 68.5 | 8.8 | 77.3 | -9 | Boros & Kolpakova 2018 |
| Sp17_18 | 96.9 | 0.5 | 0.2 | 2.4 | 8.1 | 9.6 | 73.6 | 8.7 | 82.3 | -24 | Boros & Kolpakova 2018; Boros et al., 2013 |
| Sp17_19 | 85.7 | 0.6 | 2.1 | 11.6 | 14.6 | 5.9 | 69.0 | 10.5 | 79.5 | 3 | Boros & Kolpakova 2018 |
| Sp17_20 | 67.1 | 1.9 | 7.0 | 23.9 | 14.2 | 31.0 | 46.2 | 8.6 | 54.8 | -2 | Boros & Kolpakova 2018 |

| | | | | | | | | | | | |
|---|---|---|---|---|---|---|---|---|---|---|---|
| Sp17_21 | 88.7 | 2.0 | 0.2 | 9.1 | 17.7 | 9.9 | 64.3 | 8.1 | 72.4 | -3 | Boros & Kolpakova 2018 |
| Sp17_22 | 89.9 | 0.2 | 1.6 | 8.3 | 13.7 | 8.5 | 69.3 | 8.4 | 77.7 | 3 | Boros & Kolpakova 2018 |
| Sp17_23 | 78.1 | 1.4 | 3.0 | 17.4 | 14.7 | 8.1 | 67.1 | 10.0 | 77.1 | -2 | Boros & Kolpakova 2018 |
| Sp17_24 | 88.4 | 3.0 | 0.4 | 8.2 | 30.0 | 9.6 | 53.9 | 6.5 | 60.4 | -3 | Boros & Kolpakova 2018 |
| Sp17_27 | 96.2 | 0.8 | 1.5 | 1.5 | 13.8 | 23.3 | 58.2 | 4.6 | 62.8 | -5 | Boros & Kolpakova 2018 |
| Sp17_28 | 66.6 | 2.0 | 5.5 | 26.0 | 13.9 | 22.7 | 52.8 | 10.5 | 63.3 | 5 | Boros & Kolpakova 2018 |
| Sp17_29 | 84.8 | 2.7 | 1.2 | 11.4 | 22.2 | 30.8 | 41.0 | 6.1 | 47.1 | -4 | Boros & Kolpakova 2018 |
| Sp17_31 | 61.7 | 1.9 | 0.0 | 36.4 | 30.5 | 35.7 | 25.7 | 8.0 | 33.7 | 2 | this study |
| Sp17_Neu2 | 94.0 | 2.4 | 1.0 | 2.6 | 12.6 | 34.1 | 51.6 | 1.8 | 53.4 | n.d. | Boros et al., 2013 |
| Sp18_01 | 76.3 | 3.1 | 1.5 | 19.1 | 26.5 | 7.8 | 58.3 | 7.4 | 65.7 | 7 | Boros & Kolpakova 2018 |
| Sp18_04 | 50.5 | 1.9 | 28.6 | 19.0 | 3.1 | 11.0 | 82.7 | 3.2 | 85.9 | 2 | Boros & Kolpakova 2018 |
| Sp18_05 | 61.7 | 1.9 | 0.0 | 36.4 | 30.5 | 35.7 | 25.7 | 8.0 | 33.7 | 2 | this study |
| Sp18_07 | 90.5 | 0.5 | 0.9 | 8.0 | 13.9 | 14.0 | 64.1 | 8.0 | 72.1 | 4 | Boros & Kolpakova 2018 |
| Sp18_08 | 79.3 | 4.2 | 1.1 | 15.4 | 20.1 | 6.1 | 63.8 | 10.0 | 73.8 | 5 | Boros & Kolpakova 2018 |
| Sp18_10 | 87.9 | 0.6 | 2.5 | 8.9 | 13.7 | 7.3 | 70.6 | 8.5 | 79.1 | 10 | Boros & Kolpakova 2018 |
| Sp18_11 | 71.1 | 2.1 | 4.2 | 22.6 | 12.4 | 26.5 | 50.7 | 10.4 | 61.1 | 4 | Boros & Kolpakova 2018 |
| Sp18_12 | 90.1 | 0.7 | 1.4 | 7.8 | 8.7 | 5.0 | 75.7 | 10.5 | 86.2 | 3 | Boros & Kolpakova 2018 |
| Sp18_13 | 91.7 | 2.1 | 0.2 | 6.1 | 14.6 | 7.4 | 69.1 | 8.9 | 78.0 | -10 | Boros & Kolpakova 2018 |
| Sp18_15 | 58.2 | 2.3 | 1.4 | 38.0 | 27.2 | 33.0 | 34.4 | 5.4 | 39.8 | 0 | this study |
| Sp18_16 | 80.6 | 1.2 | 1.7 | 16.5 | 14.4 | 9.1 | 66.7 | 9.8 | 76.5 | 8 | Boros & Kolpakova 2018 |
| Sp18_17 | 93.3 | 2.0 | 0.2 | 4.5 | 15.6 | 7.1 | 68.5 | 8.8 | 77.3 | -9 | Boros & Kolpakova 2018 |
| Sp18_18 | 96.9 | 0.5 | 0.2 | 2.4 | 8.1 | 9.6 | 73.6 | 8.7 | 82.3 | -24 | Boros & Kolpakova 2018; Boros et al., 2013 |
| Sp18_19 | 85.7 | 0.6 | 2.1 | 11.6 | 14.6 | 5.9 | 69.0 | 10.5 | 79.5 | 3 | Boros & Kolpakova 2018 |
| Sp18_1_d | 95.3 | 0.4 | 1.6 | 2.7 | 6.2 | 10.4 | 72.3 | 11.2 | 83.5 | -5 | Boros & Kolpakova 2018 |
| Sp18_20 | 67.1 | 1.9 | 7.0 | 23.9 | 14.2 | 31.0 | 46.2 | 8.6 | 54.8 | -2 | Boros & Kolpakova 2018 |
| Sp18_21 | 88.7 | 2.0 | 0.2 | 9.1 | 17.7 | 9.9 | 64.3 | 8.1 | 72.4 | -3 | Boros & Kolpakova 2018 |
| Sp18_22 | 89.9 | 0.2 | 1.6 | 8.3 | 13.7 | 8.5 | 69.3 | 8.4 | 77.7 | 3 | Boros & Kolpakova 2018 |
| Sp18_23 | 78.1 | 1.4 | 3.0 | 17.4 | 14.7 | 8.1 | 67.1 | 10.0 | 77.1 | -2 | Boros & Kolpakova 2018 |
| Sp18_24 | 88.4 | 3.0 | 0.4 | 8.2 | 30.0 | 9.6 | 53.9 | 6.5 | 60.4 | -3 | Boros & Kolpakova 2018 |
| Sp18_27 | 96.2 | 0.8 | 1.5 | 1.5 | 13.8 | 23.3 | 58.2 | 4.6 | 62.8 | -5 | Boros & Kolpakova 2018 |
| Sp18_28 | 66.6 | 2.0 | 5.5 | 26.0 | 13.9 | 22.7 | 52.8 | 10.5 | 63.3 | 5 | Boros & Kolpakova 2018 |
| Sp18_29 | 84.8 | 2.7 | 1.2 | 11.4 | 22.2 | 30.8 | 41.0 | 6.1 | 47.1 | -4 | Boros & Kolpakova 2018 |
| Sp18_30 | 82.5 | 2.8 | 1.6 | 13.1 | 21.2 | 49.4 | 22.9 | 6.5 | 29.4 | -2 | Boros & Kolpakova 2018 |
| Sp18_31 | 79.7 | 1.0 | 0.9 | 18.4 | 21.7 | 27.5 | 40.5 | 10.3 | 50.8 | 20 | Boros & Kolpakova 2018 |
| Sp18_9 | 83.0 | 2.1 | 1.2 | 13.8 | 17.7 | 25.5 | 46.1 | 10.7 | 56.8 | n.d. | Boros et al., 2013 |
| Sp18_Neu2 | 94.0 | 2.4 | 1.0 | 2.6 | 12.6 | 34.1 | 51.6 | 1.8 | 53.4 | n.d. | Boros et al., 2013 |
| 01Boddi | 96.5 | 0.3 | 1.1 | 2.1 | 46.5 | 8.7 | 32.3 | 12.4 | 44.7 | 3 | Boros & Kolpakova 2018 |
| 02Soser | 97.3 | 0.0 | 0.9 | 1.8 | 35.8 | 8.2 | 49.9 | 6.1 | 56.0 | 1 | Boros & Kolpakova 2018 |
| 03KolonNy | n.d. | n.d. | n.d. | n.d. | n.d. | n.d. | n.d. | n.d. | n.d. | n.d. | not found |
| 04KolonT18 | n.d. | n.d. | n.d. | n.d. | n.d. | n.d. | n.d. | n.d. | n.d. | n.d. | not found |
| 05Szelid18 | 77.3 | 0.5 | 5.7 | 16.5 | 40.7 | 4.2 | 39.7 | 15.4 | 55.1 | -5 | this study |
| 06Rusanda18 | 94.6 | 1.9 | 1.2 | 2.3 | 29.4 | 32.7 | 34.7 | 3.1 | 37.9 | -5 | Boros & Kolpakova 2018 |
| 07SlanoKopovo18 | 94.5 | 0.5 | 2.4 | 2.6 | 48.4 | 21.0 | 30.5 | 0.0 | 30.5 | -5 | Boros & Kolpakova 2018 |
| 08VBK18 | 97.4 | 0.4 | 0.7 | 1.5 | 30.6 | 9.7 | 56.8 | 2.9 | 59.7 | -3 | Korponai et al., 2020 |

| | | | | | | | | | | | |
|---|---|---|---|---|---|---|---|---|---|---|---|
| 09Kelemen18 | 95.6 | 0.6 | 1.4 | 2.4 | 35.5 | 7.1 | 39.1 | 18.3 | 57.4 | 0 | Boros & Kolpakova 2018 |
| 10Zab18 | 97.0 | 0.4 | 1.0 | 1.6 | 26.3 | 6.9 | 48.4 | 18.4 | 66.8 | 0 | Boros & Kolpakova 2018 |
| 12KH18 | 54.8 | 2.6 | 3.9 | 38.6 | 26.3 | 30.8 | 37.2 | 5.7 | 42.9 | 0 | this study |
| 13FertoB018 | 56.0 | 2.6 | 2.4 | 39.0 | 27.5 | 32.3 | 37.3 | 2.9 | 40.2 | -1 | this study |
| 14VelenceiKo18 | 44.2 | 3.2 | 0.0 | 52.6 | 24.9 | 36.3 | 29.1 | 9.6 | 38.7 | 1 | this study |
| 15Dinnyesi18 | 51.9 | 3.1 | 0.0 | 45.0 | 28.4 | 24.7 | 35.4 | 11.4 | 46.8 | 1 | this study |
| 16VelenceiNy18 | 44.5 | 3.2 | 0.0 | 52.3 | 25.4 | 37.0 | 27.8 | 9.8 | 37.6 | 2 | this study |
| 17Zicklacke18 | 84.8 | 2.7 | 1.2 | 11.4 | 22.2 | 30.8 | 41.0 | 6.1 | 47.1 | -4 | Boros & Kolpakova 2018 |
| 18ObererStinker18 | 93.3 | 2.0 | 0.2 | 4.5 | 15.6 | 7.1 | 68.5 | 8.8 | 77.3 | -9 | Boros & Kolpakova 2018 |
| 19SudlicherSilbersee18 | 88.4 | 3.0 | 0.4 | 8.2 | 30.0 | 9.6 | 53.9 | 6.5 | 60.4 | -3 | Boros & Kolpakova 2018 |
| 20LangeLacke18 | 71.1 | 2.1 | 4.2 | 22.6 | 12.4 | 26.5 | 50.7 | 10.4 | 61.1 | 4 | Boros & Kolpakova 2018 |
| 21BalatonTihany18 | 18.5 | 2.6 | 23.2 | 55.7 | 11.4 | 35.0 | 45.8 | 7.8 | 53.6 | -1 | Szilágyi2005 |
| 22BalatonKeszthely18 | 17.4 | 2.3 | 30.3 | 50.0 | 10.8 | 30.5 | 52.4 | 6.4 | 58.7 | -1 | Szilágyi2005 |
| 23UntererStinker18 | 77.2 | 1.4 | 2.1 | 19.3 | 12.9 | 5.6 | 72.6 | 8.9 | 81.5 | 1 | Boros & Kolpakova 2018 |
| 01T | 96.2 | 1.0 | 0.0 | 2.8 | 71.1 | 23.3 | 4.1 | 1.5 | 5.6 | -1 | this study |
| 02T | 73.1 | 5.4 | 0.0 | 21.5 | 32.9 | 49.5 | 11.8 | 5.8 | 17.6 | 1 | this study |
| 03T | 95.5 | 0.0 | 1.5 | 3.1 | 20.3 | 12.9 | 55.8 | 10.9 | 66.7 | 1 | this study |
| 04T | 68.9 | 0.5 | 0.0 | 30.6 | 14.2 | 8.2 | 56.6 | 21.1 | 77.7 | 0 | this study |
| 05T | 88.4 | 0.0 | 1.9 | 9.8 | 25.1 | 2.9 | 63.1 | 8.9 | 72.0 | 1 | this study |
| 06T | 97.3 | 0.0 | 0.9 | 1.8 | 35.8 | 8.2 | 49.9 | 6.1 | 56.0 | 1 | this study |
| 07T | 99.9 | 0.0 | 0.1 | 0.0 | 51.6 | 8.9 | 29.0 | 10.4 | 39.4 | -2 | this study |
| 08T | 99.9 | 0.0 | 0.1 | 0.0 | 54.2 | 5.2 | 31.7 | 8.9 | 40.6 | 1 | this study |
| 09T | 91.5 | 0.0 | 7.3 | 1.2 | 34.1 | 5.6 | 54.3 | 6.0 | 60.3 | 1 | this study |
| 10T | 98.1 | 0.0 | 0.0 | 1.9 | 37.6 | 6.4 | 40.0 | 16.0 | 56.0 | 1 | this study |
| 11T | 84.9 | 0.5 | 11.4 | 3.2 | 16.8 | 4.2 | 73.0 | 6.0 | 79.0 | -4 | this study |
| 12T | 91.4 | 0.0 | 7.1 | 1.5 | 27.0 | 5.2 | 61.6 | 6.2 | 67.8 | -1 | this study |
| 13T | 95.1 | 0.0 | 3.9 | 1.0 | 26.5 | 4.7 | 63.8 | 5.0 | 68.8 | -4 | this study |
| 14T | 97.3 | 0.0 | 0.6 | 2.0 | 24.9 | 11.7 | 53.3 | 10.1 | 63.4 | 2 | this study |
| 15T | 96.1 | 0.0 | 1.1 | 2.8 | 24.1 | 11.4 | 53.3 | 11.2 | 64.5 | 2 | this study |
| 16T | 17.5 | 1.1 | 51.8 | 29.7 | 14.8 | 16.6 | 68.6 | 0.0 | 68.6 | -1 | this study |
| 17T | 79.9 | 0.9 | 10.0 | 9.3 | 9.8 | 17.1 | 73.0 | 0.0 | 73.0 | -4 | this study |
| 18T | 47.0 | 0.7 | 30.8 | 21.5 | 11.3 | 5.5 | 83.2 | 0.0 | 83.2 | 4 | this study |
| 19T | 58.2 | 2.3 | 1.4 | 38.0 | 27.2 | 33.0 | 34.4 | 5.4 | 39.8 | 0 | this study |
| 20T | 61.7 | 1.9 | 0.0 | 36.4 | 30.5 | 35.7 | 25.7 | 8.0 | 33.7 | 2 | this study |
| 21T | 54.8 | 2.6 | 3.9 | 38.6 | 26.3 | 30.8 | 37.2 | 5.7 | 42.9 | 0 | this study |
| 22T | 55.9 | 2.6 | 2.5 | 39.0 | 27.0 | 31.8 | 38.3 | 2.9 | 41.2 | -2 | this study |
| 23T | 56.0 | 2.6 | 2.4 | 39.0 | 27.5 | 32.3 | 37.3 | 2.9 | 40.2 | -1 | this study |
| 24T | 54.2 | 2.7 | 4.0 | 39.1 | 26.3 | 30.7 | 40.2 | 2.9 | 43.1 | 1 | this study |
| 25T | 58.0 | 2.8 | 0.0 | 39.2 | 26.8 | 31.7 | 36.3 | 5.2 | 41.5 | -4 | this study |
| 26T | 77.3 | 0.5 | 5.7 | 16.5 | 40.7 | 4.2 | 39.7 | 15.4 | 55.1 | -5 | this study |
| 27T | 75.6 | 0.3 | 8.0 | 16.1 | 40.2 | 4.1 | 39.6 | 16.1 | 55.7 | -5 | this study |
| 28T | 44.5 | 3.2 | 0.0 | 52.3 | 25.4 | 37.0 | 27.8 | 9.8 | 37.6 | 2 | this study |
| 29T | 44.2 | 3.2 | 0.0 | 52.6 | 24.9 | 36.3 | 29.1 | 9.6 | 38.7 | 1 | this study |
| 30T | 43.3 | 3.1 | 0.0 | 53.5 | 24.3 | 35.6 | 31.5 | 8.6 | 40.1 | 1 | this study |

| | | | | | | | | | | | |
|---|---|---|---|---|---|---|---|---|---|---|---|
| 31T | 51.9 | 3.1 | 0.0 | 45.0 | 28.4 | 24.7 | 35.4 | 11.4 | 46.8 | 1 | this study |
| DL-M Day 0 | 98.3 | 0.8 | 0.1 | 0.9 | 1.2 | 0.1 | 62.3 | 36.4 | 98.7 | 2 | Zorz et al., 2019 |
| DLM 2015 | 98.3 | 0.8 | 0.1 | 0.9 | 1.2 | 0.1 | 62.3 | 36.4 | 98.7 | 2 | Zorz et al., 2019 |
| DLM 2017 | 98.1 | 0.8 | 0.1 | 1.1 | 1.1 | 0.2 | 18.3 | 80.4 | 98.7 | 2 | Zorz et al., 2019 |
| GEL-M Day 0 | 97.4 | 1.8 | 0.0 | 0.7 | 6.9 | 5.5 | 53.4 | 34.2 | 87.6 | 1 | Zorz et al., 2019 |
| GEM 2015 | 97.4 | 1.8 | 0.0 | 0.7 | 6.9 | 5.5 | 53.4 | 34.2 | 87.6 | 1 | Zorz et al., 2019 |
| GEM 2016 | 97.4 | 1.8 | 0.0 | 0.7 | 6.9 | 5.5 | 53.4 | 34.2 | 87.6 | 1 | Zorz et al., 2019 |
| GEM 2017 | 96.9 | 1.9 | 0.1 | 1.1 | 6.8 | 5.9 | 14.2 | 73.1 | 87.3 | -3 | Zorz et al., 2019 |
| LCL-M Day 0 | 98.9 | 0.9 | 0.0 | 0.2 | 3.6 | 8.8 | 52.9 | 34.7 | 87.6 | 10 | Zorz et al., 2019 |
| LCM 2015 | 98.9 | 0.9 | 0.0 | 0.2 | 3.6 | 8.8 | 52.9 | 34.7 | 87.6 | 10 | Zorz et al., 2019 |
| LCM 2016 | 98.9 | 0.9 | 0.0 | 0.2 | 3.6 | 8.8 | 52.9 | 34.7 | 87.6 | 10 | Zorz et al., 2019 |
| LCM 2017 | 98.9 | 0.8 | 0.0 | 0.2 | 3.2 | 8.7 | 11.6 | 76.5 | 88.1 | 10 | Zorz et al., 2019 |
| PL-M Day 0 | 99.4 | 0.4 | 0.1 | 0.2 | 2.6 | 0.3 | 60.3 | 36.7 | 97.0 | 4 | Zorz et al., 2019 |
| PLM 2015 | 99.4 | 0.4 | 0.1 | 0.2 | 2.6 | 0.3 | 60.3 | 36.7 | 97.0 | 4 | Zorz et al., 2019 |
| PLM 2016 | 99.4 | 0.4 | 0.1 | 0.2 | 2.6 | 0.3 | 60.3 | 36.7 | 97.0 | 4 | Zorz et al., 2019 |
| PLM 2017 | 98.1 | 0.8 | 0.1 | 1.1 | 1.1 | 0.2 | 18.2 | 80.5 | 98.7 | 2 | Zorz et al., 2019 |
| ML16S_10m | 96.7 | 3.0 | 0.0 | 0.2 | 40.9 | 17.3 | 41.8 | n.d. | 41.8 | 1 | Nielsen & DePaolo 2013 |
| MLW_0517_00 | 96.7 | 3.0 | 0.0 | 0.2 | 40.9 | 17.3 | 41.8 | n.d. | 41.8 | 1 | Nielsen & DePaolo 2013 |
| MLW_0517_10 | 96.7 | 3.0 | 0.0 | 0.2 | 40.9 | 17.3 | 41.8 | n.d. | 41.8 | 1 | Nielsen & DePaolo 2013 |
| MLW_0617_00 | 96.7 | 3.0 | 0.0 | 0.2 | 40.9 | 17.3 | 41.8 | n.d. | 41.8 | 1 | Nielsen & DePaolo 2013 |
| MLW_0917_00_1;MLW_0917_00_2;MLW_0917_00_3 | 96.7 | 3.0 | 0.0 | 0.2 | 40.9 | 17.3 | 41.8 | n.d. | 41.8 | 1 | Nielsen & DePaolo 2013 |
| MLW_0917_05_1;MLW_0917_05_2;MLW_0917_05_3 | 96.7 | 3.0 | 0.0 | 0.2 | 40.9 | 17.3 | 41.8 | n.d. | 41.8 | 1 | Nielsen & DePaolo 2013 |
| MLW_0917_10_1;MLW_0917_10_2;MLW_0917_10_3 | 96.7 | 3.0 | 0.0 | 0.2 | 40.9 | 17.3 | 41.8 | n.d. | 41.8 | 1 | Nielsen & DePaolo 2013 |
| MLW_1018_00 | 96.7 | 3.0 | 0.0 | 0.2 | 40.9 | 17.3 | 41.8 | n.d. | 41.8 | 1 | Nielsen & DePaolo 2013 |
| MLW_1018_05 | 96.7 | 3.0 | 0.0 | 0.2 | 40.9 | 17.3 | 41.8 | n.d. | 41.8 | 1 | Nielsen & DePaolo 2013 |
| MLW_1018_12 | 96.7 | 3.0 | 0.0 | 0.2 | 40.9 | 17.3 | 41.8 | n.d. | 41.8 | 1 | Nielsen & DePaolo 2013 |

n.d. - not determined

*defined according to Boros & Kolpakova 2018

| Sample ID | Collection date | Location | Coordinates | Publication | SRA |
|---|---|---|---|---|---|
| Bac-12S | 2012-08-01 | Kenya: Crater Lake Sonachi | n.d. | Kambura, 2017 | SRR769740 |
| Bac-18E | 2012-08-01 | Kenya: Lake Elmenteita | n.d. | Kambura, 2017 | SRR769721 |
| Bac-6M | 2012-08-01 | Kenya: Lake Magadi | n.d. | Kambura, 2017 | SRR769693 |
| B1-W | 2012-08-01 | Kenya: Lake Bogoria | n.d. | Kambura, 2017 | SRR769632 |
| B2-W | 2012-08-01 | Kenya: Lake Bogoria | n.d. | Kambura, 2017 | SRR769634 |
| KZ02 | 2015-05-04 | Kazakhstan:Ulendy | 51.534005 N 63.690941 E | | SRR28201961 |
| K03bact | 2015-05-02 | Kazakhstan:Ulendy | 51.410577 N 63.023247 E | | SRR28201960 |
| KZ04 | 2015-04-30 | Kazakhstan:Bestauskiy | 51.367788 N 62.807252 E | | SRR28201955 |
| KZ05 | 2015-05-02 | Kazakhstan:Ulendy | 51.547755 N 63.692891 E | | SRR28201954 |
| K06bact | 2015-05-02 | Kazakhstan:Bestauskiy | 51.357808 N 63.023875 E | | SRR28201953 |
| KZ08 | 2015-04-29 | Kazakhstan:Bestauskiy | 51.218719 N 62.582525 E | | SRR28201952 |
| K10bact | 2015-05-02 | Kazakhstan:Bestauskiy | 51.370077 N 63.029700 E | | SRR28201951 |
| KZ12 | 2015-05-04 | Kazakhstan:Ulendy | 51.535319 N 63.683444 E | | SRR28201950 |
| K13bact | 2015-04-24 | Kazakhstan:Naurzum | 51.461108 N 64.501608 E | | SRR28201949 |
| K15bact | 2015-04-24 | Kazakhstan:Naurzum | 51.519640 N 64.460410 E | | SRR28201959 |
| K16bact | 2015-04-29 | Kazakhstan:Urkash | 51.321700 N 62.357161 E | | SRR10674068 |
| K18bact | 2015-04-24 | Kazakhstan:Naurzum | 51.505272 N 64.493991 E | | SRR28201958 |
| KZ19 | 2015-04-30 | Kazakhstan:Bestauskiy | 51.210013 N 62.525958 E | | SRR28201957 |
| K20bact | 2015-04-25 | Kazakhstan:Naurzum | 51.588500 N 64.445855 E | | SRR28201956 |
| KT01 | 2021-06-25 | Kazakhstan:Lake Balkhash | 46.596348 N 79.221823 E | | SRR28207516 |
| KT02 | 2021-06-25 | Kazakhstan:Lake Balkhash | 46.533832 N 79.145109 E | | SRR28207551 |
| KT03 | 2021-06-25 | Kazakhstan:Lake Balkhash | 46.530906 N 79.094986 E | | SRR28207549 |
| KT04 | 2021-06-26 | Kazakhstan:Lake Balkhash | 46.301449 N 78.461765 E | | SRR28207548 |
| KT05 | 2021-06-26 | Kazakhstan:Lake Balkhash | 46.335161 N 78.432911 E | | SRR28207547 |
| KT06 | 2021-06-26 | Kazakhstan:Lake Balkhash | 46.383465 N 78.418964 E | | SRR28207546 |
| KT07 | 2021-06-27 | Kazakhstan:Lake Balkhash | 46.38716 N 78.675855 E | | SRR28207545 |
| KT08 | 2021-06-27 | Kazakhstan:Lake Balkhash | 46.370155 N 78.921802 E | | SRR28207544 |
| KT09 | 2021-06-28 | Kazakhstan:Sarkand District | 46.131397 N 78.224211 E | | SRR28207542 |
| KT10 | 2021-06-28 | Kazakhstan:Aksu District | 45.965475 N 78.325367 E | | SRR28207541 |
| KT11 | 2021-06-28 | Kazakhstan:Aksu District | 46.000596 N 78.293745 E | | SRR28207540 |
| KT12 | 2021-06-28 | Kazakhstan:Aksu District | 45.967705 N 78.327392 E | | SRR28207539 |
| KT13 | 2021-06-28 | Kazakhstan:Aksu District | 45.890096 N 78.461639 E | | SRR28207538 |
| KT14 | 2021-06-30 | Kazakhstan:Aksu District | 46.298573 N 78.215385 E | | SRR28207537 |
| KT15 | 2021-06-30 | Kazakhstan:Aksu District | 46.298894 N 78.474564 E | | SRR28207536 |
| KT16 | 2021-06-30 | Kazakhstan:Lake Balkhash | 46.285278 N 78.43688 E | | SRR28207535 |
| KT17 | 2021-07-01 | Kazakhstan:Lake Balkhash | 46.462005 N 79.053697 E | | SRR28207534 |
| KT18 | 2021-07-01 | Kazakhstan:Lake Balkhash | 46.455212 N 79.042745 E | | SRR28207543 |
| KT19 | 2021-07-02 | Kazakhstan:Karatal District | 45.666505 N 78.081244 E | | SRR28207531 |
| KT20 | 2021-07-02 | Kazakhstan:Karatal District | 45.743376 N 78.006456 E | | SRR28207530 |
| KT21 | 2021-07-20 | Kazakhstan:Lake Sasykkol | 46.488666 N 80.860082 E | | SRR28207529 |

| | | | | | |
|---|---|---|---|---|---|
| KT22 | 2021-07-20 | Kazakhstan:Lake Alakol | 46.39485 N 81.382522 E | | SRR28207527 |
| KT23 | 2021-07-20 | Kazakhstan:Lake Alakol | 46.468637 N 81.54499 E | | SRR28207526 |
| KT24 | 2021-07-21 | Kazakhstan:Lake Alakol | 46.450718 N 81.507218 E | | SRR28207525 |
| KT25 | 2021-07-21 | Kazakhstan:Lake Alakol | 46.43978 N 81.452084 E | | SRR28207524 |
| KT26 | 2021-07-21 | Kazakhstan:Lake Alakol | 46.429081 N 81.422508 E | | SRR28207523 |
| KT27 | 2021-07-21 | Kazakhstan:Lake Alakol | 45.996245 N 81.366174 E | | SRR28207522 |
| 0S | 2018-09-01 | Kazakhstan:Lake Balkhash | n.d. | | SRR28207514 |
| 02S | 2018-09-01 | Kazakhstan:Lake Balkhash | n.d. | | SRR28207513 |
| 04S | 2018-09-01 | Kazakhstan:Lake Balkhash | n.d. | | SRR28207512 |
| 06S | 2018-09-01 | Kazakhstan:Lake Balkhash | n.d. | | SRR28207511 |
| 08S | 2018-09-01 | Kazakhstan:Lake Balkhash | n.d. | | SRR28207510 |
| 09S | 2018-09-01 | Kazakhstan:Lake Balkhash | n.d. | | SRR28207509 |
| 11S | 2018-09-01 | Kazakhstan:Lake Balkhash | n.d. | | SRR28207508 |
| SHV | 2018-07-12 | Kazakhstan:Shalkar | 47.77216 N 59.56621 E | | SRR28207533 |
| KA1 | 2019-05-14 | Kazakhstan:Shalkar | 48.05287 N 59.57196 E | | SRR28207532 |
| KA2 | 2019-05-15 | Kazakhstan:Shalkar | 47.52056 N 59.07887 E | | SRR28207550 |
| KA3 | 2019-05-15 | Kazakhstan:Shalkar | 47.51450 N 59.08419 E | | SRR28207528 |
| KA4 | 2019-05-15 | Kazakhstan:Shalkar | 47.72670 N 59.14562 E | | SRR28207521 |
| KA5 | 2019-05-15 | Kazakhstan:Shalkar | 47.78893 N 59.58138 E | | SRR28207520 |
| KA6 | 2019-05-17 | Kazakhstan:Shalkar | 47.84792 N 59.61716 E | | SRR28207519 |
| KA7 | 2019-05-19 | Kazakhstan:Belkopa | 50.06536 N 59.59568 E | | SRR28207518 |
| KA8 | 2019-06-01 | Kazakhstan:Akkol | 43.54248 N 70.70400 E | | SRR28207517 |
| Gudzh_water | 2017-10 | Russia: Buryatiya, Suvo | 53.64000 N 109.96000 E | Lavrentyeva et al., 2020 | SRR10801163 |
| NB30 | 2013-09-05 | Russia:Lake Doroninskoe, Transbaikalia | 51.23000 N 112.23000 E | Matyugina et al., 2018 | SRR6329113 |
| NB31 | 2013-09-05 | Russia:Lake Doroninskoe, Transbaikalia | 51.23000 N 112.23000 E | Matyugina et al., 2018 | SRR6329112 |
| NB32 | 2013-09-05 | Russia:Lake Doroninskoe, Transbaikalia | 51.23000 N 112.23000 E | Matyugina et al., 2018 | SRR6329115 |
| NB41 | 2013-09-06 | Russia:Lake Doroninskoe, Transbaikalia | 51.23000 N 112.23000 E | Matyugina et al., 2018 | SRR6329106 |
| NB42 | 2013-09-06 | Russia:Lake Doroninskoe, Transbaikalia | 51.23000 N 112.23000 E | Matyugina et al., 2018 | SRR6329105 |
| NB43 | 2013-09-06 | Russia:Lake Doroninskoe, Transbaikalia | 51.23000 N 112.23000 E | Matyugina et al., 2018 | SRR6329104 |
| Bangong Co | 2015-07-01 | China:Tibetan Plateau | 33.50688333 N 79.81849166 E | Ji et al. 2019 FEMS Micr Ecol | SRR8156978 |
| Bong Co | 2015-07-01 | China:Tibetan Plateau | 31.22901666 N 91.10875 E | Ji et al. 2019 FEMS Micr Ecol | SRR8156959 |
| Co Ngoin (1) | 2015-07-01 | China:Tibetan Plateau | 31.68666666 N 88.70861111 E | Ji et al. 2019 FEMS Micr Ecol | SRR8156955 |
| Mapam Yumco | 2015-07-01 | China:Tibetan Plateau | 30.74192222 N 81.57108055 E | Ji et al. 2019 FEMS Micr Ecol | SRR8156992 |
| Urru Co | 2015-07-01 | China:Tibetan Plateau | 31.77962777 N 88.07386111 E | Ji et al. 2019 FEMS Micr Ecol | SRR8156920 |
| Bam Co | 2015-07-01 | China:Tibetan Plateau | 31.33558611 N 90.58961666 E | Ji et al. 2019 FEMS Micr Ecol | SRR8156983 |
| Bero Zeco | 2015-07-01 | China:Tibetan Plateau | 32.41774444 N 82.97188611 E | Ji et al. 2019 FEMS Micr Ecol | SRR8156953 |
| Co Ngoin (2) | 2015-07-01 | China:Tibetan Plateau | 31.52669444 N 91.53168611 E | Ji et al. 2019 FEMS Micr Ecol | SRR8156923 |
| Dawa Co | 2015-07-01 | China:Tibetan Plateau | 31.22166944 N 85.00583611 E | Ji et al. 2019 FEMS Micr Ecol | SRR8156926 |
| Tangra Yumco | 2015-07-01 | China:Tibetan Plateau | 31.35506388 N 86.6539 E | Ji et al. 2019 FEMS Micr Ecol | SRR8156932 |
| Dong Co | 2015-07-01 | China:Tibetan Plateau | 32.15925 N 84.72611111 E | Ji et al. 2019 FEMS Micr Ecol | SRR8156963 |
| Kunggyu | 2015-07-01 | China:Tibetan Plateau | 30.63209444 N 82.14183611 E | Ji et al. 2019 FEMS Micr Ecol | SRR8156971 |

| | | | | | |
|---|---|---|---|---|---|
| Nam Co | 2015-07-01 | China:Tibetan Plateau | 30.66490555 N 90.80185277 E | Ji et al. 2019 FEMS Micr Ecol | SRR8156946 |
| Pung Co | 2015-07-01 | China:Tibetan Plateau | 31.44505277 N 90.92033055 E | Ji et al. 2019 FEMS Micr Ecol | SRR8156975 |
| Selin Co | 2015-07-01 | China:Tibetan Plateau | 31.68833333 N 88.78416666 E | Ji et al. 2019 FEMS Micr Ecol | SRR8156976 |
| Zhari Namco | 2015-07-01 | China:Tibetan Plateau | 30.99296666 N 85.59155833 E | Ji et al. 2019 FEMS Micr Ecol | SRR8156995 |
| Zhaxi Co | 2015-07-01 | China:Tibetan Plateau | 32.19537777 N 85.07547777 E | Ji et al. 2019 FEMS Micr Ecol | SRR8156974 |
| Bangkog Co | 2015-07-01 | China:Tibetan Plateau | 31.71047777 N 89.43625 E | Ji et al. 2019 FEMS Micr Ecol | SRR8156952 |
| Dangqiong Co | 2015-07-01 | China:Tibetan Plateau | 31.56467777 N 86.78088333 E | Ji et al. 2019 FEMS Micr Ecol | SRR8156966 |
| Nyer Co (Nieer Co) | 2015-07-01 | China:Tibetan Plateau | 32.28039166 N 82.2315 E | Ji et al. 2019 FEMS Micr Ecol | SRR8156981 |
| BU1 | 2012-11-29 | Hungary:Szabadszallas | 46.8663333 N 19.1692167 E | Szabo et al., 2017 | SRR4289875 |
| SO1 | 2012-11-29 | Hungary:Dunatetetlen | 46.7890167 N 19.14464999 E | Szabo et al., 2017 | SRR4292558 |
| ZA1 | 2012-11-29 | Hungary:Szabadszallas | 46.8365 N 19.17138330 E | Szabo et al., 2017 | SRR4292556 |
| SO2 | 2013-04-17 | Hungary:Dunatetetlen | 46.7890167 N 19.14464999 E | Szabo et al., 2020 | SRR8433480 |
| SO3 | 2013-05-30 | Hungary:Dunatetetlen | 46.7890167 N 19.14464999 E | Szabo et al., 2020 | SRR8433481 |
| SO4 | 2013-06-18 | Hungary:Dunatetetlen | 46.7890167 N 19.14464999 E | Szabo et al., 2020 | SRR8433482 |
| SO5 | 2013-07-24 | Hungary:Dunatetetlen | 46.7890167 N 19.14464999 E | Szabo et al., 2020 | SRR8433483 |
| SO6 | 2013-08-16 | Hungary:Dunatetetlen | 46.7890167 N 19.14464999 E | Szabo et al., 2020 | SRR8433484 |
| SO9 | 2013-11-19 | Hungary:Dunatetetlen | 46.7890167 N 19.14464999 E | Szabo et al., 2020 | SRR8433485 |
| SO11 | 2014-01-10 | Hungary:Dunatetetlen | 46.7890167 N 19.14464999 E | Szabo et al., 2020 | SRR8433487 |
| SO12 | 2014-02-26 | Hungary:Dunatetetlen | 46.7890167 N 19.14464999 E | Szabo et al., 2020 | SRR8433488 |
| SO13 | 2014-03-26 | Hungary:Dunatetetlen | 46.7890167 N 19.14464999 E | Szabo et al., 2020 | SRR8433489 |
| SO14 | 2014-04-23 | Hungary:Dunatetetlen | 46.7890167 N 19.14464999 E | Szabo et al., 2020 | SRR8433474 |
| SO15 | 2014-05-22 | Hungary:Dunatetetlen | 46.7890167 N 19.14464999 E | Szabo et al., 2020 | SRR8433475 |
| SO16 | 2014-06-18 | Hungary:Dunatetetlen | 46.7890167 N 19.14464999 E | Szabo et al., 2020 | SRR8433476 |
| VBK | 2014-04-23 | Hungary:Soltszentimre | 46.7636333 N 19.1804667 E | Korponai et al., 2019 | SRR8433706 |
| ZA2 | 2013-04-17 | Hungary:Szabadszallas | 46.8365 N 19.17138330 E | Szabo et al., 2020 | SRR8433477 |
| ZA3 | 2013-05-30 | Hungary:Szabadszallas | 46.8365 N 19.17138330 E | Szabo et al., 2020 | SRR8433470 |
| ZA4 | 2013-06-18 | Hungary:Szabadszallas | 46.8365 N 19.17138330 E | Szabo et al., 2020 | SRR8433471 |
| ZA5 | 2013-07-24 | Hungary:Szabadszallas | 46.8365 N 19.17138330 E | Szabo et al., 2020 | SRR8433472 |
| ZA6 | 2013-08-16 | Hungary:Szabadszallas | 46.8365 N 19.17138330 E | Szabo et al., 2020 | SRR8433473 |
| ZA7 | 2013-09-24 | Hungary:Szabadszallas | 46.8365 N 19.17138330 E | Szabo et al., 2020 | SRR8433478 |
| ZA8 | 2013-10-17 | Hungary:Szabadszallas | 46.8365 N 19.17138330 E | Szabo et al., 2020 | SRR8433479 |
| ZA9 | 2013-11-19 | Hungary:Szabadszallas | 46.8365 N 19.17138330 E | Szabo et al., 2020 | SRR8433491 |
| ZA10 | 2013-12-09 | Hungary:Szabadszallas | 46.8365 N 19.17138330 E | Szabo et al., 2020 | SRR8433490 |
| ZA11 | 2014-01-10 | Hungary:Szabadszallas | 46.8365 N 19.17138330 E | Szabo et al., 2020 | SRR8433493 |
| ZA12 | 2014-02-26 | Hungary:Szabadszallas | 46.8365 N 19.17138330 E | Szabo et al., 2020 | SRR8433492 |
| ZA13 | 2014-03-26 | Hungary:Szabadszallas | 46.8365 N 19.17138330 E | Szabo et al., 2020 | SRR8433495 |
| ZA14 | 2014-04-23 | Hungary:Szabadszallas | 46.8365 N 19.17138330 E | Szabo et al., 2020 | SRR8433494 |
| ZA15 | 2014-05-22 | Hungary:Szabadszallas | 46.8365 N 19.17138330 E | Szabo et al., 2020 | SRR8433497 |
| ZA16 | 2014-06-18 | Hungary:Szabadszallas | 46.8365 N 19.17138330 E | Szabo et al., 2020 | SRR8433496 |
| ZA17 | 2014-07-29 | Hungary:Szabadszallas | 46.8365 N 19.17138330 E | Szabo et al., 2020 | SRR8433498 |
| M05 | 2013-07-19 | Romania:Lacul Ursu | 46.603731 N 25.086142 E | | SRR28211548 |

| | | | | | |
|---|---|---|---|---|---|
| U05 | 2013-07-19 | Romania:Ocna Mureș | 46.384823 N 23.860119 E | | SRR28211547 |
| U15 | 2013-07-19 | Romania:Ocna Mureș | 46.384823 N 23.860119 E | | SRR28211546 |
| U28 | 2013-07-19 | Romania:Ocna Mureș | 46.384823 N 23.860119 E | | SRR28211545 |
| B01w | 2015-11-10 | Hungary:Lake Ferto | 47.73459 N 16.71941 E | Szuroczki et al., 2020 | SRR10841575 |
| B02w | 2015-12-07 | Hungary:Lake Ferto | 47.73459 N 16.71941 E | | SRR28201989 |
| B03w | 2016-02-22 | Hungary:Lake Ferto | 47.73459 N 16.71941 E | | SRR28201988 |
| B04w | 2016-03-29 | Hungary:Lake Ferto | 47.73459 N 16.71941 E | | SRR28201979 |
| B05w | 2016-04-18 | Hungary:Lake Ferto | 47.73459 N 16.71941 E | | SRR28201978 |
| B06w | 2016-05-18 | Hungary:Lake Ferto | 47.73459 N 16.71941 E | | SRR28201977 |
| B07w | 2016-06-20 | Hungary:Lake Ferto | 47.73459 N 16.71941 E | | SRR28201976 |
| B08w | 2016-07-18 | Hungary:Lake Ferto | 47.73459 N 16.71941 E | Szuroczki et al., 2020 | SRR10841574 |
| B09w | 2016-08-20 | Hungary:Lake Ferto | 47.73459 N 16.71941 E | | SRR28201975 |
| B010w | 2016-09-21 | Hungary:Lake Ferto | 47.73459 N 16.71941 E | | SRR28201974 |
| B011w | 2016-10-25 | Hungary:Lake Ferto | 47.73459 N 16.71941 E | | SRR28201973 |
| KH1w | 2015-11-10 | Hungary:Lake Ferto | 47.68460 N 16.70272 E | Szuroczki et al., 2020 | SRR10841571 |
| KH2w | 2015-12-07 | Hungary:Lake Ferto | 47.68460 N 16.70272 E | | SRR28201972 |
| KH3w | 2016-02-22 | Hungary:Lake Ferto | 47.68460 N 16.70272 E | | SRR28201987 |
| KH4w | 2016-03-29 | Hungary:Lake Ferto | 47.68460 N 16.70272 E | | SRR28201986 |
| KH5w | 2016-04-18 | Hungary:Lake Ferto | 47.68460 N 16.70272 E | | SRR28201985 |
| KH6w | 2016-05-18 | Hungary:Lake Ferto | 47.68460 N 16.70272 E | | SRR28201984 |
| KH7w | 2016-06-20 | Hungary:Lake Ferto | 47.68460 N 16.70272 E | | SRR28201983 |
| KH8w | 2016-07-18 | Hungary:Lake Ferto | 47.68460 N 16.70272 E | Szuroczki et al., 2020 | SRR10841570 |
| KH9w | 2016-08-20 | Hungary:Lake Ferto | 47.68460 N 16.70272 E | | SRR28201982 |
| KH10w | 2016-09-21 | Hungary:Lake Ferto | 47.68460 N 16.70272 E | | SRR28201981 |
| KH11w | 2016-10-25 | Hungary:Lake Ferto | 47.68460 N 16.70272 E | | SRR28201980 |
| B01 | 2017-04-12 | Hungary:Dunatetetlen | 46.767833 N 19.150117 E | | SRR22959451 |
| B02 | 2017-04-26 | Hungary:Dunatetetlen | 46.767833 N 19.150117 E | | SRR22959450 |
| B03 | 2017-05-17 | Hungary:Dunatetetlen | 46.767833 N 19.150117 E | | SRR22959437 |
| B04 | 2017-05-29 | Hungary:Dunatetetlen | 46.767833 N 19.150117 E | | SRR22959434 |
| B05 | 2017-06-15 | Hungary:Dunatetetlen | 46.767833 N 19.150117 E | | SRR22959423 |
| B06 | 2017-06-29 | Hungary:Dunatetetlen | 46.767833 N 19.150117 E | | SRR22959372 |
| B07 | 2017-03-13 | Hungary:Dunatetetlen | 46.767833 N 19.150117 E | | SRR22959417 |
| B08 | 2017-07-27 | Hungary:Dunatetetlen | 46.767833 N 19.150117 E | | SRR22959366 |
| B09 | 2017-08-14 | Hungary:Dunatetetlen | 46.767833 N 19.150117 E | | SRR22959347 |
| B10 | 2017-08-29 | Hungary:Dunatetetlen | 46.767833 N 19.150117 E | | SRR22959400 |
| B11 | 2017-09-11 | Hungary:Dunatetetlen | 46.767833 N 19.150117 E | | SRR22959449 |
| B12 | 2017-09-27 | Hungary:Dunatetetlen | 46.767833 N 19.150117 E | | SRR22959390 |
| B13 | 2017-10-16 | Hungary:Dunatetetlen | 46.767833 N 19.150117 E | | SRR22959333 |
| B14 | 2017-11-14 | Hungary:Dunatetetlen | 46.767833 N 19.150117 E | | SRR22959326 |
| K01 | 2017-04-12 | Hungary:Fulopszallas | 46.798217 N 19.174000 E | | SRR22959443 |
| K02 | 2017-04-26 | Hungary:Fulopszallas | 46.798217 N 19.174000 E | | SRR22959442 |

| | | | | |
|---|---|---|---|---|
| K03 | 2017-05-17 | Hungary:Fulopszallas | 46.798217 N 19.174000 E | SRR22959441 |
| K04 | 2017-05-29 | Hungary:Fulopszallas | 46.798217 N 19.174000 E | SRR22959440 |
| K05 | 2017-06-15 | Hungary:Fulopszallas | 46.798217 N 19.174000 E | SRR22959439 |
| K06 | 2017-06-29 | Hungary:Fulopszallas | 46.798217 N 19.174000 E | SRR22959438 |
| K09 | 2017-08-14 | Hungary:Fulopszallas | 46.798217 N 19.174000 E | SRR22959436 |
| K12 | 2017-09-27 | Hungary:Fulopszallas | 46.798217 N 19.174000 E | SRR22959387 |
| K14 | 2017-11-14 | Hungary:Fulopszallas | 46.798217 N 19.174000 E | SRR22959386 |
| S01 | 2017-04-12 | Hungary:Dunatetetlen | 46.789017 N 19.144650 E | SRR22959426 |
| S02 | 2017-04-26 | Hungary:Dunatetetlen | 46.789017 N 19.144650 E | SRR22959425 |
| S03 | 2017-05-17 | Hungary:Dunatetetlen | 46.789017 N 19.144650 E | SRR22959424 |
| S04 | 2017-05-29 | Hungary:Dunatetetlen | 46.789017 N 19.144650 E | SRR22959422 |
| S05 | 2017-06-15 | Hungary:Dunatetetlen | 46.789017 N 19.144650 E | SRR22959421 |
| S06 | 2017-06-29 | Hungary:Dunatetetlen | 46.789017 N 19.144650 E | SRR22959420 |
| S07 | 2017-03-13 | Hungary:Dunatetetlen | 46.789017 N 19.144650 E | SRR22959379 |
| S08 | 2017-07-27 | Hungary:Dunatetetlen | 46.789017 N 19.144650 E | SRR22959378 |
| S09 | 2017-08-14 | Hungary:Dunatetetlen | 46.789017 N 19.144650 E | SRR22959377 |
| S10 | 2017-08-29 | Hungary:Dunatetetlen | 46.789017 N 19.144650 E | SRR22959376 |
| S11 | 2017-09-11 | Hungary:Dunatetetlen | 46.789017 N 19.144650 E | SRR22959375 |
| S12 | 2017-09-27 | Hungary:Dunatetetlen | 46.789017 N 19.144650 E | SRR22959374 |
| S13 | 2017-10-16 | Hungary:Dunatetetlen | 46.789017 N 19.144650 E | SRR22959373 |
| S14 | 2017-11-14 | Hungary:Dunatetetlen | 46.789017 N 19.144650 E | SRR22959355 |
| V01 | 2017-04-12 | Hungary:Soltszentimre | 46.7636333 N 19.1804667 E | SRR22959385 |
| V02 | 2017-04-26 | Hungary:Soltszentimre | 46.7636333 N 19.1804667 E | SRR22959384 |
| V03 | 2017-05-17 | Hungary:Soltszentimre | 46.7636333 N 19.1804667 E | SRR22959383 |
| V04 | 2017-05-29 | Hungary:Soltszentimre | 46.7636333 N 19.1804667 E | SRR22959382 |
| V05 | 2017-06-15 | Hungary:Soltszentimre | 46.7636333 N 19.1804667 E | SRR22959381 |
| V06 | 2017-06-29 | Hungary:Soltszentimre | 46.7636333 N 19.1804667 E | SRR22959380 |
| V07 | 2017-03-13 | Hungary:Soltszentimre | 46.7636333 N 19.1804667 E | SRR22959435 |
| V08 | 2017-07-27 | Hungary:Soltszentimre | 46.7636333 N 19.1804667 E | SRR22959433 |
| V09 | 2017-08-14 | Hungary:Soltszentimre | 46.7636333 N 19.1804667 E | SRR22959432 |
| V10 | 2017-08-29 | Hungary:Soltszentimre | 46.7636333 N 19.1804667 E | SRR22959431 |
| V11 | 2017-09-11 | Hungary:Soltszentimre | 46.7636333 N 19.1804667 E | SRR22959430 |
| V12 | 2017-09-27 | Hungary:Soltszentimre | 46.7636333 N 19.1804667 E | SRR22959429 |
| V13 | 2017-10-16 | Hungary:Soltszentimre | 46.7636333 N 19.1804667 E | SRR22959428 |
| V14 | 2017-11-14 | Hungary:Soltszentimre | 46.7636333 N 19.1804667 E | SRR22959427 |
| Z01 | 2017-04-12 | Hungary:Szabadszallas | 46.836500 N 19.171383 E | SRR22959354 |
| Z02 | 2017-04-26 | Hungary:Szabadszallas | 46.836500 N 19.171383 E | SRR22959353 |
| Z03 | 2017-05-17 | Hungary:Szabadszallas | 46.836500 N 19.171383 E | SRR22959352 |
| Z04 | 2017-05-29 | Hungary:Szabadszallas | 46.836500 N 19.171383 E | SRR22959351 |
| Z05 | 2017-06-15 | Hungary:Szabadszallas | 46.836500 N 19.171383 E | SRR22959350 |
| Z06 | 2017-06-29 | Hungary:Szabadszallas | 46.836500 N 19.171383 E | SRR22959349 |

| | | | | | |
|---|---|---|---|---|---|
| Z07 | 2017-03-13 | Hungary:Szabadszallas | 46.836500 N 19.171383 E | | SRR22959348 |
| Z08 | 2017-07-27 | Hungary:Szabadszallas | 46.836500 N 19.171383 E | | SRR22959419 |
| Z09 | 2017-08-14 | Hungary:Szabadszallas | 46.836500 N 19.171383 E | | SRR22959418 |
| Z10 | 2017-08-29 | Hungary:Szabadszallas | 46.836500 N 19.171383 E | | SRR22959416 |
| Z12 | 2017-09-27 | Hungary:Szabadszallas | 46.836500 N 19.171383 E | | SRR22959415 |
| Z14 | 2017-11-14 | Hungary:Szabadszallas | 46.836500 N 19.171383 E | | SRR22959414 |
| BV | 2013-06-01 | Hungary:Lake Balaton | 46.73 N 17.29 E | Borsodi et al., 2017 | SRR2186745 |
| K0 | 2014-11-18 | Hungary:Izsak | 46.76055556 N 19.34027778 E | Mentes et al., 2018 | SRR5658453;SRR5658454 |
| K01 | 2014-11-18 | Hungary:Izsak | 46.76055556 N 19.34027778 E | Mentes et al., 2018 | SRR5658455;SRR5658457 |
| A | 2018-08-15 | Hungary:Tiszaalpar | 46.8254970 N 20.003828 E | | SRR28205851 |
| B | 2018-08-15 | Hungary:Tiszaalpar | 46.8250223 N 20.0019207 E | | SRR28205850 |
| B0 | 2018-08-07 | Hungary:Lake Ferto | 47.73459 N 16.71941 E | | SRR28205838 |
| C | 2018-08-15 | Hungary:Lakitelek | 46.838694 N 20.008271 E | | SRR28205831 |
| D | 2018-08-15 | Hungary:Lakitelek | 46.839376 N 20.007884 E | | SRR28205830 |
| Ds | 2018-08-09 | Hungary:Kaposvar | 46.3981821 N 17.8193651 E | | SRR28205829 |
| E | 2018-08-15 | Hungary:Martely | 46.469000 N 20.225709 E | | SRR28205828 |
| F | 2018-08-15 | Hungary:Martely | 46.471944 N 20.223845 E | | SRR28205827 |
| FDN | 2018-07-30 | Hungary:Oxbow Fadd-Dombori | n.d. | | SRR28205826 |
| FDO | 2018-07-30 | Hungary:Oxbow Fadd-Dombori | n.d. | | SRR28205825 |
| Ff | 2018-08-06 | Hungary:Lake Balaton | 46.945098 N 18.010683 E | | SRR28205849 |
| K | 2018-08-06 | Hungary:Lake Balaton | 46.731294 N 17.268830 E | | SRR28205847 |
| KH | 2018-08-07 | Hungary:Lake Ferto | 47.68460 N 16.70272 E | | SRR28205846 |
| Kn | 2018-08-09 | Hungary:Kovacsszenaja | 46.175485 N 18.114118 E | | SRR28205845 |
| LK | 2018-08-02 | Hungary:Lipot | 47.865348 N 17.464860 E | | SRR28205844 |
| LN | 2018-08-02 | Hungary:Lipot | 47.864854 N 17.464530 E | | SRR28205843 |
| NH | 2018-08-07 | Hungary:Lake Ferto (Nagy-Herlakni) | n.d. | | SRR28205842 |
| R | 2018-08-13 | Hungary:Homorud | 46.011237 N 18.7661610 E | | SRR28205841 |
| TLO | 2018-07-30 | Hungary:Oxbow Tolnai | n.d. | | SRR28205840 |
| TNB | 2018-08-31 | Hungary:Lake Tisza | 47.641913 N 20.662493 E | | SRR28205839 |
| TO | 2018-08-31 | Hungary:Lake Tisza | 47.643105 N 20.662063 E | | SRR28205837 |
| TS | 2018-08-31 | Hungary:Lake Tisza | 47.642014 N 20.665917 E | | SRR28205836 |
| Z | 2018-08-06 | Hungary:Lake Balaton | 46.705299 N 17.264026 E | | SRR28205835 |
| SS15 | 2012-01-16 | Austria:Seewinkel | 47.791389 N 16.780556 E | Siclair et al., 2015 | SRR1519332 |
| SS16 | 2012-02-06 | Austria:Seewinkel | 47.791389 N 16.780556 E | Siclair et al., 2015 | SRR1519333 |
| SS17 | 2012-03-22 | Austria:Seewinkel | 47.791389 N 16.780556 E | Siclair et al., 2015 | SRR1519334 |
| US15 | 2012-01-16 | Austria:Seewinkel | 47.801667 N 16.783889 E | Siclair et al., 2015 | SRR1519335 |
| US16 | 2012-02-06 | Austria:Seewinkel | 47.801667 N 16.783889 E | Siclair et al., 2015 | SRR1519336 |
| US17 | 2012-03-22 | Austria:Seewinkel | 47.801667 N 16.783889 E | Siclair et al., 2015 | SRR1519337 |
| ZL15 | 2012-01-16 | Austria:Seewinkel | 47.768056 N 16.780833 E | Siclair et al., 2015 | SRR1519338 |
| ZL16 | 2012-02-06 | Austria:Seewinkel | 47.768056 N 16.780833 E | Siclair et al., 2015 | SRR1519339 |
| ZL17 | 2012-03-22 | Austria:Seewinkel | 47.768056 N 16.780833 E | Siclair et al., 2015 | SRR1519340 |

| | | | | | |
|---|---|---|---|---|---|
| Sp17_01 | 2017-04-03 - 06 | Austria:Seewinkel | 47.77513639 N 16.77011917 E | Szabo et al., 2022 | SRR16560283 |
| Sp17_02 | 2017-04-03 - 06 | Austria:Seewinkel | 47.78917083 N 16.88677778 E | Szabo et al., 2022 | SRR16560282 |
| Sp17_04 | 2017-04-03 - 06 | Austria:Seewinkel | 47.81771528 N 16.86491056 E | Szabo et al., 2022 | SRR16560271 |
| Sp17_05 | 2017-04-03 - 06 | Hungary:Fertozug | 47.681925 N 16.84113889 E | Szabo et al., 2022 | SRR16560260 |
| Sp17_07 | 2017-04-03 - 06 | Austria:Seewinkel | 47.78612111 N 16.84219833 E | Szabo et al., 2022 | SRR16560249 |
| Sp17_09 | 2017-04-03 - 06 | Austria:Seewinkel | 47.75602861 N 16.78047972 E | Szabo et al., 2022 | SRR16560238 |
| Sp17_10 | 2017-04-03 - 06 | Austria:Seewinkel | 47.79273556 N 16.87862583 E | Szabo et al., 2022 | SRR16560237 |
| Sp17_11 | 2017-04-03 - 06 | Austria:Seewinkel | 47.75747528 N 16.87876472 E | Szabo et al., 2022 | SRR16560236 |
| Sp17_12 | 2017-04-03 - 06 | Austria:Seewinkel | 47.75050167 N 16.856695 E | Szabo et al., 2022 | SRR16560235 |
| Sp17_13 | 2017-04-03 - 06 | Austria:Seewinkel | 47.80675889 N 16.78752194 E | Szabo et al., 2022 | SRR16560234 |
| Sp17_14 | 2017-04-03 - 06 | Austria:Seewinkel | 47.768036 N 16.844325 E | | SRR22904843 |
| Sp17_15 | 2017-04-03 - 06 | Hungary:Fertozug | 47.67755 N 16.83405556 E | Szabo et al., 2022 | SRR16560281 |
| Sp17_16 | 2017-04-03 - 06 | Austria:Seewinkel | 47.82711944 N 16.80793222 E | Szabo et al., 2022 | SRR16560280 |
| Sp17_17 | 2017-04-03 - 06 | Austria:Seewinkel | 47.81376222 N 16.79251722 E | Szabo et al., 2022 | SRR16560279 |
| Sp17_18 | 2017-04-03 - 06 | Austria:Seewinkel | 47.81073583 N 16.84461722 E | Szabo et al., 2022 | SRR16560278 |
| Sp17_19 | 2017-04-03 - 06 | Austria:Seewinkel | 47.79044083 N 16.86618583 E | Szabo et al., 2022 | SRR16560277 |
| Sp17_20 | 2017-04-03 - 06 | Austria:Seewinkel | 47.77334361 N 16.88009389 E | Szabo et al., 2022 | SRR16560276 |
| Sp17_21 | 2017-04-03 - 06 | Austria:Seewinkel | 47.78574472 N 16.79274361 E | Szabo et al., 2022 | SRR16560275 |
| Sp17_22 | 2017-04-03 - 06 | Austria:Seewinkel | 47.78378861 N 16.88411194 E | Szabo et al., 2022 | SRR16560274 |
| Sp17_23 | 2017-04-03 - 06 | Austria:Seewinkel | 47.79890389 N 16.87170639 E | Szabo et al., 2022 | SRR16560273 |
| Sp17_24 | 2017-04-03 - 06 | Austria:Seewinkel | 47.79113278 N 16.77973694 E | Szabo et al., 2022 | SRR16560272 |
| Sp17_27 | 2017-04-03 - 06 | Austria:Seewinkel | 47.79007667 N 16.85234944 E | Szabo et al., 2022 | SRR16560269 |
| Sp17_28 | 2017-04-03 - 06 | Austria:Seewinkel | 47.77092167 N 16.87077667 E | Szabo et al., 2022 | SRR16560268 |
| Sp17_29 | 2017-04-03 - 06 | Austria:Seewinkel | 47.76696444 N 16.78475194 E | Szabo et al., 2022 | SRR16560267 |
| Sp17_31 | 2017-04-03 - 06 | Hungary:Fertozug | 47.681925 N 16.84113889 E | | SRR22904842 |
| Sp17_Neu2 | 2017-04-03 - 06 | Austria:Seewinkel | 47.76447667 N 16.84024833 | Szabo et al., 2022 | SRR16560266 |
| Sp18_01 | 2018-04-02 - 04 | Austria:Seewinkel | 47.77513639 N 16.77011917 | Szabo et al., 2022 | SRR16560264 |
| Sp18_04 | 2018-04-02 - 04 | Austria:Seewinkel | 47.81771528 N 16.86491056 E | Szabo et al., 2022 | SRR16560263 |
| Sp18_05 | 2018-04-02 - 04 | Hungary:Fertozug | 47.681925 N 16.84113889 E | Szabo et al., 2022 | SRR16560262 |
| Sp18_07 | 2018-04-02 - 04 | Austria:Seewinkel | 47.78612111 N 16.84219833 E | Szabo et al., 2022 | SRR16560261 |
| Sp18_08 | 2018-04-02 - 04 | Austria:Seewinkel | 47.75867861 N 16.78548861 E | | SRR22904841 |
| Sp18_10 | 2018-04-02 - 04 | Austria:Seewinkel | 47.79273556 N 16.87862583 E | Szabo et al., 2022 | SRR16560258 |
| Sp18_11 | 2018-04-02 - 04 | Austria:Seewinkel | 47.75747528 N 16.87876472 E | Szabo et al., 2022 | SRR16560236 |
| Sp18_12 | 2018-04-02 - 04 | Austria:Seewinkel | 47.75050167 N 16.856695 E | Szabo et al., 2022 | SRR16560256 |
| Sp18_13 | 2018-04-02 - 04 | Austria:Seewinkel | 47.80675889 N 16.78752194 E | Szabo et al., 2022 | SRR16560255 |
| Sp18_15 | 2018-04-02 - 04 | Hungary:Fertozug | 47.67755 N 16.83405556 E | Szabo et al., 2022 | SRR16560254 |
| Sp18_16 | 2018-04-02 - 04 | Austria:Seewinkel | 47.82711944 N 16.80793222 E | Szabo et al., 2022 | SRR16560253 |
| Sp18_17 | 2018-04-02 - 04 | Austria:Seewinkel | 47.81376222 N 16.79251722 E | Szabo et al., 2022 | SRR16560252 |
| Sp18_18 | 2018-04-02 - 04 | Austria:Seewinkel | 47.81073583 N 16.84461722 E | Szabo et al., 2022 | SRR16560251 |
| Sp18_19 | 2018-04-02 - 04 | Austria:Seewinkel | 47.79044083 N 16.86618583 E | Szabo et al., 2022 | SRR16560250 |
| Sp18_1_d | 2018-04-02 - 04 | Austria:Seewinkel | 47.78917083 N 16.88677778 E | Szabo et al., 2022 | SRR16560265 |

| | | | | | |
|---|---|---|---|---|---|
| Sp18_20 | 2018-04-02 - 04 | Austria:Seewinkel | 47.77334361 N 16.88009389 E | Szabo et al., 2022 | SRR16560248 |
| Sp18_21 | 2018-04-02 - 04 | Austria:Seewinkel | 47.78574472 N 16.79274361 E | Szabo et al., 2022 | SRR16560247 |
| Sp18_22 | 2018-04-02 - 04 | Austria:Seewinkel | 47.78378861 N 16.88411194 E | Szabo et al., 2022 | SRR16560246 |
| Sp18_23 | 2018-04-02 - 04 | Austria:Seewinkel | 47.79890389 N 16.87170639 E | Szabo et al., 2022 | SRR16560245 |
| Sp18_24 | 2018-04-02 - 04 | Austria:Seewinkel | 47.79113278 N 16.77973694 E | Szabo et al., 2022 | SRR16560244 |
| Sp18_27 | 2018-04-02 - 04 | Austria:Seewinkel | 47.79007667 N 16.85234944 E | Szabo et al., 2022 | SRR16560242 |
| Sp18_28 | 2018-04-02 - 04 | Austria:Seewinkel | 47.77092167 N 16.87077667 E | Szabo et al., 2022 | SRR16560241 |
| Sp18_29 | 2018-04-02 - 04 | Austria:Seewinkel | 47.76696444 N 16.78475194 E | Szabo et al., 2022 | SRR16560240 |
| Sp18_30 | 2018-04-02 - 04 | Austria:Seewinkel | 47.72157583 N 16.82404028 E | | SRR22904840 |
| Sp18_31 | 2018-04-02 - 04 | Austria:Seewinkel | 47.744564 N 16.769988 E | | SRR22904839 |
| Sp18_9 | 2018-04-02 - 04 | Austria:Seewinkel | 47.75602861 N 16.78047972 E | Szabo et al., 2022 | SRR16560259 |
| Sp18_Neu2 | 2018-04-02 - 04 | Austria:Seewinkel | 47.76447667 N 16.84024833 | Szabo et al., 2022 | SRR16560239 |
| 01Boddi | 2018-05-16 | Hungary:Dunatetetlen | 46.767102 N 19.151592 E | | SRR28205968 |
| 02Soser | 2018-05-16 | Hungary:Dunatetetlen | 46.789017 N 19.144650 E | | SRR28205967 |
| 03KolonNy | 2018-05-16 | Hungary:Izsak | 46.760556 N 19.340278 E | | SRR28205848 |
| 04KolonT18 | 2018-05-16 | Hungary:Izsak | 46.772997 N 19.340000 E | | SRR28205834 |
| 05Szelid18 | 2018-05-16 | Hungary:Lake Szelid | 46.629000 N 19.044000 E | | SRR28205955 |
| 06Rusanda18 | 2018-05-17 | Serbia:Melenci | 45.528717 N 20.291136 E | | SRR28205944 |
| 07SlanoKopovo18 | 2018-05-17 | Serbia:Novi Becej | 45.628363 N 20.204091 E | | SRR28205933 |
| 08VBK18 | 2018-05-17 | Hungary:Soltszentimre | 46.763633 N 19.180467 E | | SRR28205923 |
| 09Kelemen18 | 2018-05-17 | Hungary:Fulopszallas | 46.788645 N 19.172599 E | | SRR28205958 |
| 10Zab18 | 2018-05-17 | Hungary:Szabadszallas | 46.827921 N 19.173656 E | | SRR28205922 |
| 12KH18 | 2018-05-22 | Hungary:Lake Ferto | 47.692311 N 16.714088 E | | SRR28205921 |
| 13FertoB018 | 2018-05-22 | Hungary:Lake Ferto | 47.733561 N 16.720937 E | | SRR28205920 |
| 14VelenceiKo18 | 2018-05-22 | Hungary:Gardony | 47.206431 N 18.603350 E | | SRR28205966 |
| 15Dinnyesi18 | 2018-05-22 | Hungary:Dinnyes | 47.171115 N 18.546452 E | | SRR28205965 |
| 16VelenceiNy18 | 2018-05-22 | Hungary:Pakozd | 47.199834 N 18.560681 E | | SRR28205964 |
| 17Zicklacke18 | 2018-05-23 | Austria:Illmitz | 47.766282 N 16.783803 E | | SRR28205963 |
| 18ObererStinker18 | 2018-05-23 | Austria:Illmitz | 47.816178 N 16.795678 E | | SRR28205962 |
| 19SudlicherSilbersee18 | 2018-05-23 | Austria:Illmitz | 47.791089 N 16.779713 E | | SRR28205961 |
| 20LangeLacke18 | 2018-05-23 | Austria:Apetlon | 47.758951 N 16.877194 E | | SRR28205960 |
| 21BalatonTihany18 | 2018-05-23 | Hungary:Tihany | 46.917583 N 17.918469 E | | SRR28205833 |
| 22BalatonKeszthely18 | 2018-05-23 | Hungary:Keszthely | 46.735972 N 17.268286 E | | SRR28205832 |
| 23UntererStinker18 | 2018-05-23 | Austria:Illmitz | 47.802617 N 16.785083 E | | SRR28205959 |
| 01T | 2021-05-18 | Hungary:Sarkeresztur | 46.985362 N 18.550139 E | | SRR28205957 |
| 02T | 2021-05-18 | Hungary:Sarszentagota | 46.971521 N 18.554462 E | | SRR28205956 |
| 03T | 2021-05-19 | Hungary:Pusztaszer | 46.547839 N 20.029932 E | | SRR28205954 |
| 04T | 2021-05-19 | Hungary:Pusztaszer | 46.523881 N 20.040926 E | | SRR28205953 |
| 05T | 2021-05-19 | Hungary:Baks | 46.570495 N 20.062962 E | | SRR28205952 |
| 06T | 2021-05-21 | Hungary:Dunatetetlen | 46.788859 N 19.146177 E | | SRR28205951 |
| 07T | 2021-05-21 | Hungary:Dunatetetlen | 46.763919 N 19.152522 E | | SRR28205950 |

| | | | | | |
|---|---|---|---|---|---|
| 08T | 2021-05-21 | Hungary:Dunatetetlen | 46.765875 N 19.153309 E | | SRR28205949 |
| 09T | 2021-05-21 | Hungary:Fulopszallas | 46.794747 N 19.186312 E | | SRR28205948 |
| 10T | 2021-05-21 | Hungary:Fulopszallas | 46.7994 N 19.177578 E | | SRR28205947 |
| 11T | 2021-05-22 | Hungary:Szabadszallas | 46.862657 N 19.167097 E | | SRR28205946 |
| 12T | 2021-05-22 | Hungary:Szabadszallas | 46.839562 N 19.176819 E | | SRR28205945 |
| 13T | 2021-05-22 | Hungary:Szabadszallas | 46.828591 N 19.17471 E | | SRR28205943 |
| 14T | 2021-05-26 | Hungary:Kardoskut | 46.472287 N 20.626551 E | | SRR28205942 |
| 15T | 2021-05-26 | Hungary:Kardoskut | 46.471891 N 20.629927 E | | SRR28205941 |
| 16T | 2021-05-27 | Hungary:Apaj | 47.156851 N 19.113268 E | | SRR28205940 |
| 17T | 2021-05-27 | Hungary:Apaj | 47.152225 N 19.104857 E | | SRR28205939 |
| 18T | 2021-05-27 | Hungary:Apaj | 47.142151 N 19.113607 E | | SRR28205938 |
| 19T | 2021-06-01 | Hungary:Fertozug | 47.676847 N 16.832391 E | | SRR28205937 |
| 20T | 2021-06-01 | Hungary:Fertozug | 47.680646 N 16.842439 E | | SRR28205936 |
| 21T | 2021-06-01 | Hungary:Lake Ferto | 47.68460 N 16.70272 E | | SRR28205935 |
| 22T | 2021-06-01 | Hungary:Lake Ferto | 47.711684 N 16.704966 E | | SRR28205934 |
| 23T | 2021-06-01 | Hungary:Lake Ferto | 47.734414 N 16.720417 E | | SRR28205932 |
| 24T | 2021-06-01 | Hungary:Fertozug | 47.675823 N 16.730948 E | | SRR28205931 |
| 25T | 2021-06-01 | Hungary:Fertozug | 47.639279 N 16.778645 E | | SRR28205930 |
| 26T | 2021-06-11 | Hungary:Lake Szelid | 46.632289 N 19.060764 E | | SRR28205929 |
| 27T | 2021-06-11 | Hungary:Lake Szelid | 46.621628 N 19.036824 E | | SRR28205928 |
| 28T | 2021-06-18 | Hungary:Lake Velence | 47.207631 N 18.60649 E | | SRR28205927 |
| 29T | 2021-06-18 | Hungary:Lake Velence | 47.194294 N 18.567652 E | | SRR28205926 |
| 30T | 2021-06-18 | Hungary:Lake Velence | 47.18908 N 18.559821 E | | SRR28205925 |
| 31T | 2021-06-18 | Hungary:Dinnyes | 47.164809 N 18.547284 E | | SRR28205924 |
| DL-M Day 0 | 2014-06-18 | Canada: Cariboo Plateau | 51.35416 N 121.24527 W | Zorz et al., 2019 | SRR5291571 |
| DLM 2015 | 2015-05-26 | Canada: Cariboo Plateau | 51.35416 N 121.24527 W | Zorz et al., 2019 | SRR7799343 |
| DLM 2017 | 2017-05-16 | Canada: Cariboo Plateau | 51.35416 N 121.24527 W | Zorz et al., 2019 | SRR7799352 |
| GEL-M Day 0 | 2014-06-18 | Canada: Cariboo Plateau | 51.329722 N 121.641389 W | Zorz et al., 2019 | SRR5291562 |
| GEM 2015 | 2015-05-26 | Canada: Cariboo Plateau | 51.329722 N 121.641389 W | Zorz et al., 2019 | SRR7799345 |
| GEM 2016 | 2016-05-17 | Canada: Cariboo Plateau | 51.329722 N 121.641389 W | Zorz et al., 2019 | SRR7799349 |
| GEM 2017 | 2017-05-16 | Canada: Cariboo Plateau | 51.329722 N 121.641389 W | Zorz et al., 2019 | SRR7799350 |
| LCL-M Day 0 | 2014-06-18 | Canada: Cariboo Plateau | 51.3275 N 121.63305 W | Zorz et al., 2019 | SRR5291553 |
| LCM 2015 | 2015-05-26 | Canada: Cariboo Plateau | 51.3275 N 121.63305 W | Zorz et al., 2019 | SRR7799344 |
| LCM 2016 | 2016-05-17 | Canada: Cariboo Plateau | 51.3275 N 121.63305 W | Zorz et al., 2019 | SRR7799348 |
| LCM 2017 | 2017-05-16 | Canada: Cariboo Plateau | 51.3275 N 121.63305 W | Zorz et al., 2019 | SRR7799347 |
| PL-M Day 0 | 2014-06-18 | Canada: Cariboo Plateau | 51.45055 N 121.3875 W | Zorz et al., 2019 | SRR5291544 |
| PLM 2015 | 2015-05-26 | Canada: Cariboo Plateau | 51.45055 N 121.3875 W | Zorz et al., 2019 | SRR7799342 |
| PLM 2016 | 2016-05-17 | Canada: Cariboo Plateau | 51.45055 N 121.3875 W | Zorz et al., 2019 | SRR7799346 |
| PLM 2017 | 2017-05-16 | Canada: Cariboo Plateau | 51.45055 N 121.3875 W | Zorz et al., 2019 | SRR7799351 |
| ML16S_10m | 2012-07-01 | USA: California, Mono Lake | 37.96 N 119.02 W | Edwardson & Hollibaugh, 2018 | SRR3475672 |
| MLW_0517_00 | 2017-05-23 | USA: California, Mono Lake | 37.977 N 118.991 W | Philips et al., 2021 | SRR13742070 |

| | | | | | |
|---|---|---|---|---|---|
| MLW_0517_10 | 2017-05-23 | USA: California, Mono Lake | 37.977 N 118.991 W | Philips et al., 2021 | SRR13742069 |
| MLW_0617_00 | 2017-06-22 | USA: California, Mono Lake | 37.977 N 118.991 W | Philips et al., 2021 | SRR13742059 |
| MLW_0917_00_1;MLW_0917_00_2;MLW_0917_00_3 | 2017-09-19 | USA: California, Mono Lake | 37.977 N 118.991 W | Philips et al., 2021 | SRR13742068;SRR13742067;SRR13742098 |
| MLW_0917_05_1;MLW_0917_05_2;MLW_0917_05_3 | 2017-09-19 | USA: California, Mono Lake | 37.977 N 118.991 W | Philips et al., 2021 | SRR13742097;SRR13742096;SRR13742095 |
| MLW_0917_10_1;MLW_0917_10_2;MLW_0917_10_3 | 2017-09-19 | USA: California, Mono Lake | 37.977 N 118.991 W | Philips et al., 2021 | SRR13742094;SRR13742093;SRR13742092 |
| MLW_1018_00 | 2018-10-09 | USA: California, Mono Lake | 37.977 N 118.991 W | Philips et al., 2021 | SRR13742072 |
| MLW_1018_05 | 2018-10-09 | USA: California, Mono Lake | 37.977 N 118.991 W | Philips et al., 2021 | SRR13742071 |
| MLW_1018_12 | 2018-10-09 | USA: California, Mono Lake | 37.977 N 118.991 W | Philips et al., 2021 | SRR13742066 |

| Sample ID | Water depth (m) | Sampling depth (m) | Conductivity (mS/cm) | Salinity (g/L) from conductivity | Salinity (g/L) from TDS | pH | T (°C) | Dissolved oxigen (mg/l) | Chlorophyll-a (µg/l) | TOC (mg/L) | DOC (mg/L) |
|---|---|---|---|---|---|---|---|---|---|---|---|
| Bac-12S | 7.00 | 0.01 | n.d. | n.d. | 8.2 | 9.5 | n.d. | n.d. | n.d. | n.d. | n.d. |
| Bac-18E | 2.00 | 0.01 | n.d. | n.d. | 13.2 | 9.4 | n.d. | n.d. | n.d. | n.d. | n.d. |
| Bac-6M | 5.00 | 0.01 | 68.6 | n.d. | 38.1 | 9.8 | n.d. | n.d. | n.d. | n.d. | n.d. |
| B1-W | 10.00 | 0.01 | 67.0 | 43.1 | 79.3 | 10.0 | n.d. | n.d. | n.d. | n.d. | 45 |
| B2-W | 10.00 | 0.01 | 67.0 | 43.1 | 79.3 | 10.0 | n.d. | n.d. | n.d. | n.d. | 45 |
| KZ02 | 0.20 | 0.01 | 21.4 | n.d. | 16.4 | 8.7 | 19.4 | 9.2 | 57.2 | 62.9 | n.d. |
| K03bact | 0.20 | 0.01 | 99.5 | n.d. | 79.2 | 8.9 | 18.7 | 13.5 | 9.6 | 124.8 | n.d. |
| KZ04 | 0.10 | 0.01 | 69.8 | n.d. | 69.7 | 8.5 | 24.4 | 10.5 | 9.4 | 25.2 | n.d. |
| KZ05 | 0.40 | 0.10 | 1.3 | n.d. | 0.7 | 8.9 | 17.1 | 12.6 | 19.4 | 21.8 | n.d. |
| K06bact | 0.10 | 0.01 | 159.5 | n.d. | 148.7 | 8.1 | 25.9 | 12.9 | 9.7 | 74.2 | n.d. |
| KZ08 | 0.20 | 0.01 | 99.3 | n.d. | 88.2 | 8.7 | 14.0 | n.d. | 40.8 | 65.8 | n.d. |
| K10bact | 0.70 | 0.10 | 22.2 | n.d. | 14.0 | 8.6 | 23.6 | 14.3 | 7.5 | 43.3 | n.d. |
| KZ12 | 0.10 | 0.01 | 31.3 | n.d. | 23.1 | 8.5 | 20.0 | 9.3 | 46.8 | 78.6 | n.d. |
| K13bact | 0.20 | 0.01 | 7.7 | n.d. | 13.4 | 7.9 | 9.6 | n.d. | 12.3 | 12.9 | n.d. |
| K15bact | 0.20 | 0.01 | 67.9 | n.d. | 49.1 | 9.5 | 11.5 | n.d. | 6 | 48.5 | n.d. |
| K16bact | 0.70 | 0.10 | 10.2 | n.d. | 6.4 | 8.9 | 14.7 | 19.4 | 378 | 56.2 | n.d. |
| K18bact | 0.50 | 0.10 | 2.7 | n.d. | 1.9 | 8.4 | 9.4 | n.d. | 53.4 | 24.2 | n.d. |
| KZ19 | 0.20 | 0.01 | 137.3 | n.d. | 131.0 | 8.2 | n.d. | n.d. | 23 | 48 | n.d. |
| K20bact | 0.03 | 0.01 | n.d. | n.d. | 0.5 | 8.5 | 10.4 | n.d. | 10.7 | 24.4 | n.d. |
| KT01 | 0.30 | 0.01 | 8.1 | n.d. | 5.7 | 8.9 | 18.4 | 9.2 | 1.6 | 6.8 | 5.2 |
| KT02 | 0.07 | 0.01 | 60.6 | n.d. | 85.0 | 8.7 | 25.7 | 9.2 | 2 | 5.7 | 4.5 |
| KT03 | 0.95 | 0.01 | 8.3 | n.d. | 5.8 | 8.9 | 23.2 | 8.9 | 1.9 | 6.8 | 5.9 |
| KT04 | 0.67 | 0.01 | 5.9 | n.d. | 4.1 | 9.0 | 20.4 | 9.1 | 1.7 | 5.4 | 4.2 |
| KT05 | 0.45 | 0.01 | 14.5 | n.d. | 9.2 | 9.2 | 22.4 | 16.6 | 28.1 | 0.1 | 0.1 |
| KT06 | 0.40 | 0.01 | 6.4 | n.d. | 4.5 | 8.9 | 21.7 | 8.9 | 1.2 | 6.2 | 5.6 |
| KT07 | 0.18 | 0.01 | 6.9 | n.d. | 4.8 | 8.9 | 26.1 | 8.7 | 1.4 | 0.1 | 0.1 |
| KT08 | 0.40 | 0.01 | 7.3 | n.d. | 5.1 | 8.9 | 25.8 | 8.4 | 1.1 | 7.5 | 2.9 |
| KT09 | 0.73 | 0.01 | 63.0 | n.d. | 50.7 | 8.1 | 21.9 | 6.9 | 1.7 | 15.6 | 13.9 |
| KT10 | 0.31 | 0.01 | 51.1 | n.d. | 51.1 | 9.2 | 24.9 | 9.3 | 0.9 | 27.5 | 27.5 |
| KT11 | 1.00 | 0.01 | 3.3 | n.d. | 2.5 | 8.7 | 28.9 | 9.1 | 2.5 | 23.9 | 22.1 |
| KT12 | 0.15 | 0.01 | 20.9 | n.d. | 19.3 | 9.5 | 31.7 | 13.1 | 3.9 | 0.1 | 0.1 |
| KT13 | 0.20 | 0.01 | 107.8 | 115.7 | 368.2 | 9.6 | 29.5 | 15.2 | 18.1 | 0.1 | 0.1 |
| KT14 | 0.04 | 0.01 | 181.1 | 196.3 | 370.1 | 7.5 | 30.3 | 0.8 | 8.8 | 8 | 0.1 |
| KT15 | 0.20 | 0.01 | 151.6 | 163.8 | 279.1 | 7.7 | 36.1 | 18.0 | 5.8 | 0.1 | 0.1 |
| KT16 | 0.15 | 0.01 | 8.5 | n.d. | 6.8 | 8.7 | 29.9 | 8.0 | 2.9 | 0.1 | 0.1 |
| KT17 | 0.03 | 0.01 | 145.7 | 157.4 | 525.8 | 7.3 | 34.2 | 1.3 | 7.8 | 0.1 | 0.1 |
| KT18 | 0.50 | 0.01 | 7.4 | n.d. | 8.1 | 8.8 | 27.9 | 7.6 | 1.4 | 19.6 | 17.8 |
| KT19 | 0.57 | 0.01 | 3.0 | n.d. | 2.1 | 10.3 | 28.8 | 13.8 | 1.5 | 57.8 | 40.1 |
| KT20 | 2.00 | 0.01 | 2.8 | n.d. | 1.9 | 9.1 | 29.7 | 11.7 | 13.7 | 22.7 | 18.9 |
| KT21 | 0.40 | 0.01 | 0.7 | n.d. | 1.3 | 8.7 | 26.4 | 9.6 | 8.1 | 0.1 | 0.1 |
| KT22 | 1.00 | 0.01 | 16.6 | n.d. | 9.2 | 8.7 | 29.5 | 8.9 | 24.3 | 0.1 | 0.1 |
| KT23 | 0.50 | 0.01 | 9.7 | n.d. | 7.6 | 9.0 | 30.5 | 14.4 | 1.6 | 0.1 | 0.1 |
| KT24 | 0.45 | 0.01 | 10.2 | n.d. | 6.8 | 9.0 | 27.3 | 7.7 | 2.4 | 0.1 | 0.1 |
| KT25 | 0.33 | 0.01 | 12.3 | n.d. | 9.5 | 9.4 | 30.1 | 14.3 | 4 | 0.1 | 0.1 |
| KT26 | 0.09 | 0.01 | 8.6 | n.d. | 6.8 | 8.9 | 32.2 | 10.5 | 3.8 | 0.1 | 0.1 |
| KT27 | 0.20 | 0.01 | 13.5 | n.d. | 9.4 | 9.2 | 29.2 | 7.8 | 8.1 | 0.1 | 0.1 |
| 0S | n.d. | n.d. | n.d. | n.d. | 1.2 | 8.7 | n.d. | n.d. | n.d. | n.d. | n.d. |

| | | | | | | | | | | | |
|---|---|---|---|---|---|---|---|---|---|---|---|
| 02S | n.d. | n.d. | n.d. | n.d. | 2.4 | 8.6 | n.d. | n.d. | n.d. | n.d. | n.d. |
| 04S | n.d. | n.d. | n.d. | n.d. | 2.1 | 8.6 | n.d. | n.d. | n.d. | n.d. | n.d. |
| 06S | n.d. | n.d. | n.d. | n.d. | 1.2 | 8.6 | n.d. | n.d. | n.d. | n.d. | n.d. |
| 08S | n.d. | n.d. | n.d. | n.d. | 1.2 | 8.5 | n.d. | n.d. | n.d. | n.d. | n.d. |
| 09S | n.d. | n.d. | n.d. | n.d. | 0.2 | 7.0 | n.d. | n.d. | n.d. | n.d. | n.d. |
| 11S | n.d. | n.d. | n.d. | n.d. | 3.3 | 8.4 | n.d. | n.d. | n.d. | n.d. | n.d. |
| SHV | 0.29 | 0.05 | 3.5 | n.d. | 2.6 | 9.3 | 30.3 | n.d. | n.d. | n.d. | n.d. |
| KA1 | 0.20 | 0.05 | 78.0 | n.d. | 58.2 | 9.4 | 22.4 | n.d. | n.d. | n.d. | n.d. |
| KA2 | 0.30 | 0.05 | 14.3 | n.d. | 9.6 | 8.9 | 23.0 | 9.0 | n.d. | n.d. | n.d. |
| KA3 | 0.30 | 0.05 | 65.2 | n.d. | 45.5 | 8.8 | 29.2 | 13.2 | n.d. | n.d. | n.d. |
| KA4 | 0.18 | 0.05 | 135.2 | n.d. | 103.9 | 8.4 | 20.6 | 8.6 | n.d. | n.d. | n.d. |
| KA5 | n.d. | 0.05 | n.d. | n.d. | 1.0 | n.d. | n.d. | n.d. | n.d. | n.d. | n.d. |
| KA6 | 0.20 | 0.05 | 181.8 | n.d. | 247.7 | 8.2 | 24.5 | 7.7 | n.d. | n.d. | n.d. |
| KA7 | n.d. | 0.05 | 41.0 | n.d. | 32.4 | 8.0 | 15.6 | 9.8 | n.d. | n.d. | n.d. |
| KA8 | 0.65 | 0.05 | 4.4 | n.d. | 3.8 | 8.0 | 25.5 | 6.3 | n.d. | n.d. | n.d. |
| Gudzh_water | 0.10 | 0.10 | n.d. | n.d. | 110.1 | 9.7 | 12.0 | n.d. | n.d. | n.d. | n.d. |
| NB30 | 6.20 | 2.50 | 23.4 | 22.2 | 26.5 | 10.1 | 16.3 | 6.1 | n.d. | n.d. | n.d. |
| NB31 | 6.20 | 2.50 | 34.3 | 32.6 | 26.5 | 10.0 | 17.1 | 4.9 | n.d. | n.d. | n.d. |
| NB32 | 6.20 | 3.00 | 35.8 | 34.0 | 26.5 | 10.0 | 16.7 | 4.5 | n.d. | n.d. | n.d. |
| NB41 | 6.20 | 3.00 | 23.4 | 22.2 | 26.5 | 10.1 | 16.2 | 5.9 | n.d. | n.d. | n.d. |
| NB42 | 6.20 | 3.15 | 35.4 | 33.6 | 26.5 | 10.0 | 15.1 | 4.9 | n.d. | n.d. | n.d. |
| NB43 | 6.20 | 3.15 | 36.2 | 34.4 | 26.5 | 10.0 | 15.0 | 4.4 | n.d. | n.d. | n.d. |
| Bangong Co | n.d. | 0 - 0,3 | n.d. | 0.5 | n.d. | 8.9 | 16.1 | 6.2 | n.d. | 5.89 | n.d. |
| Bong Co | n.d. | 0 - 0,3 | n.d. | 0.1 | n.d. | 8.7 | 12.5 | 5.7 | n.d. | 3.09 | n.d. |
| Co Ngoin (1) | n.d. | 0 - 0,3 | n.d. | 0.2 | n.d. | 9.0 | 15.6 | 6.0 | n.d. | 9.00 | n.d. |
| Mapam Yumco | n.d. | 0 - 0,3 | n.d. | 0.2 | n.d. | 8.9 | 12.4 | 6.2 | n.d. | 2.48 | n.d. |
| Urru Co | n.d. | 0 - 0,3 | n.d. | 0.2 | n.d. | 8.8 | 13.2 | 6.4 | n.d. | 7.61 | n.d. |
| Bam Co | n.d. | 0 - 0,3 | n.d. | 7.3 | n.d. | 9.7 | 14.7 | 6.1 | n.d. | 14.88 | n.d. |
| Bero Zeco | n.d. | 0 - 0,3 | n.d. | 29.7 | n.d. | 9.0 | 18.1 | 4.4 | n.d. | 34.39 | n.d. |
| Co Ngoin (2) | n.d. | 0 - 0,3 | n.d. | 3.8 | n.d. | 9.5 | 15.9 | 5.1 | n.d. | 18.92 | n.d. |
| Dawa Co | n.d. | 0 - 0,3 | n.d. | 17.7 | n.d. | 9.3 | n.d. | n.d. | n.d. | n.d. | n.d. |
| Tangra Yumco | n.d. | 0 - 0,3 | n.d. | 6.8 | n.d. | 9.3 | 14.0 | 5.4 | n.d. | 3.93 | n.d. |
| Dong Co | n.d. | 0 - 0,3 | n.d. | 42.2 | n.d. | 8.9 | n.d. | n.d. | n.d. | n.d. | n.d. |
| Kunggyu | n.d. | 0 - 0,3 | n.d. | 4.9 | n.d. | 9.4 | 15.3 | 6.2 | n.d. | 52.55 | n.d. |
| Nam Co | n.d. | 0 - 0,3 | n.d. | 0.9 | n.d. | 8.7 | n.d. | n.d. | n.d. | n.d. | n.d. |
| Pung Co | n.d. | 0 - 0,3 | n.d. | 8.6 | n.d. | 9.8 | 15.5 | 5.2 | n.d. | 12.96 | n.d. |
| Selin Co | n.d. | 0 - 0,3 | n.d. | 7.8 | n.d. | 9.4 | 13.5 | 5.1 | n.d. | 6.65 | n.d. |
| Zhari Namco | n.d. | 0 - 0,3 | n.d. | 10.1 | n.d. | 9.5 | 14.2 | 6.0 | n.d. | 8.78 | n.d. |
| Zhaxi Co | n.d. | 0 - 0,3 | n.d. | 13.0 | n.d. | 9.4 | 18.1 | 6.0 | n.d. | 15.14 | n.d. |
| Bangkog Co | n.d. | 0 - 0,3 | n.d. | 56.9 | n.d. | 9.5 | 15.9 | 2.4 | n.d. | 66.69 | n.d. |
| Dangqiong Co | n.d. | 0 - 0,3 | n.d. | 118.1 | n.d. | n.d. | n.d. | n.d. | n.d. | n.d. | n.d. |
| Nyer Co (Nieer Co) | n.d. | 0 - 0,3 | n.d. | 109.4 | n.d. | 8.1 | n.d. | n.d. | n.d. | n.d. | n.d. |
| BU1 | 0.02 | 0.01 | 4.6 | 3.7 | n.d. | 9.2 | 14.7 | n.d. | 59.6 | n.d. | 48 |
| SO1 | 0.25 | 0.05 | 13.3 | 10.6 | n.d. | 9.2 | 13.3 | n.d. | 20.8 | n.d. | 814 |
| ZA1 | 0.07 | 0.05 | 11.0 | 8.8 | n.d. | 9.7 | 14.1 | n.d. | 32.4 | n.d. | 60 |
| SO2 | 0.50 | 0.05 | 1.9 | 1.5 | n.d. | 8.3 | 17.0 | n.d. | 1.5 | 85 | 84 |
| SO3 | 0.45 | 0.05 | 3.0 | 2.4 | n.d. | 8.5 | 17.0 | n.d. | 3.6 | 148 | 144 |
| SO4 | 0.35 | 0.05 | 3.8 | 3.0 | n.d. | 8.7 | 30.0 | n.d. | 0.5 | 174 | 174 |

| | | | | | | | | | | | |
|---|---|---|---|---|---|---|---|---|---|---|---|
| SO5 | 0.18 | 0.05 | 7.9 | 6.3 | n.d. | 9.1 | 25.0 | n.d. | 1.8 | 355 | 350 |
| SO6 | 0.10 | 0.02 | 23.6 | 18.9 | n.d. | 9.6 | 27.0 | n.d. | 463 | 1223 | 1164 |
| SO9 | 0.04 | 0.01 | 11.4 | 9.1 | n.d. | 10.0 | 8.0 | n.d. | 348.9 | 672 | 626 |
| SO11 | 0.09 | 0.02 | 9.5 | 7.6 | n.d. | 9.9 | 5.0 | n.d. | 379 | 614 | 595 |
| SO12 | 0.18 | 0.05 | 4.5 | 3.6 | n.d. | 9.1 | 7.0 | n.d. | 148.5 | 294 | 276 |
| SO13 | 0.13 | 0.03 | 6.8 | 5.4 | n.d. | 9.2 | 11.0 | n.d. | 64.9 | 459 | 449 |
| SO14 | 0.08 | 0.02 | 10.7 | 8.6 | n.d. | 9.3 | 21.0 | n.d. | 49.4 | 697 | 682 |
| SO15 | 0.13 | 0.02 | 7.7 | 6.2 | n.d. | 9.3 | 25.0 | n.d. | 2.2 | 476 | 466 |
| SO16 | 0.07 | 0.02 | 15.0 | 12.0 | n.d. | 9.6 | 24.0 | n.d. | 92 | 855 | 830 |
| VBK | 0.05 | 0.04 | 15.5 | 12.4 | n.d. | 10.2 | n.d. | n.d. | 10.6 | 1143 | n.d. |
| ZA2 | 0.45 | 0.05 | 2.4 | 1.9 | n.d. | 9.1 | 24.0 | n.d. | 6.8 | 37 | 31 |
| ZA3 | 0.31 | 0.05 | 3.0 | 2.4 | n.d. | 9.3 | 17.0 | n.d. | 11.7 | 38 | 33 |
| ZA4 | 0.32 | 0.05 | 3.3 | 2.6 | n.d. | 9.2 | 26.0 | n.d. | 1.7 | 40 | 32 |
| ZA5 | 0.20 | 0.05 | 5.5 | 4.4 | n.d. | 9.5 | 31.0 | n.d. | 3.8 | 54 | 50 |
| ZA6 | 0.13 | 0.03 | 9.4 | 7.5 | n.d. | 9.8 | 30.0 | n.d. | 44.1 | 84 | 75 |
| ZA7 | 0.13 | 0.03 | 12.3 | 9.8 | n.d. | 10.0 | 16.0 | n.d. | 153.2 | 128 | 92 |
| ZA8 | 0.06 | 0.01 | 8.0 | 6.4 | n.d. | 10.1 | 17.0 | n.d. | 259.8 | 101 | 60 |
| ZA9 | 0.04 | 0.01 | 9.4 | 7.5 | n.d. | 10.2 | 11.0 | n.d. | 430.2 | 115 | 74 |
| ZA10 | 0.06 | 0.01 | 6.4 | 5.1 | n.d. | 10.2 | 4.0 | n.d. | 455.7 | 96 | 59 |
| ZA11 | 0.04 | 0.01 | 8.0 | 6.4 | n.d. | 10.2 | 6.0 | n.d. | 1043.7 | 138 | 64 |
| ZA12 | 0.12 | 0.02 | 4.8 | 3.8 | n.d. | 9.6 | 10.0 | n.d. | 1499.4 | 127 | 41 |
| ZA13 | 0.07 | 0.01 | 6.4 | 5.1 | n.d. | 9.7 | 14.0 | n.d. | 2555.5 | 180 | 53 |
| ZA14 | 0.03 | 0.01 | 9.0 | 7.2 | n.d. | 9.7 | 25.0 | n.d. | 822 | 221 | 74 |
| ZA15 | 0.03 | 0.01 | 8.3 | 6.6 | n.d. | 9.4 | 28.0 | n.d. | 59.6 | 152 | 97 |
| ZA16 | 0.02 | 0.01 | 21.2 | 17.0 | n.d. | 9.9 | 28.0 | n.d. | 199 | 239 | 217 |
| ZA17 | 0.07 | 0.01 | 12.6 | 10.1 | n.d. | 9.7 | 31.0 | n.d. | 6.8 | 150 | 142 |
| M05 | 18.00 | 0.5 | 92.0 | 66.6 | n.d. | 8.4 | 27.0 | n.d. | 0.6 | n.d. | n.d. |
| U05 | 23.00 | 0.5 | 46.8 | 30.0 | n.d. | 8.3 | 27.2 | 9.0 | 2 | 18.6 | n.d. |
| U15 | 23.00 | 1.5 | 128.2 | 68.0 | n.d. | 7.5 | 38.2 | 9.8 | 0.6 | 13 | n.d. |
| U28 | 23.00 | 2.8 | 204.0 | 134.0 | n.d. | 7.6 | 41.1 | 11.6 | 2.5 | 15.88 | n.d. |
| B01w | 1.50 | 0.5 | 1.9 | 1.5 | n.d. | 9.7 | 9.8 | n.d. | 7.8 | n.d. | n.d. |
| B02w | 1.40 | 0.5 | 1.9 | 1.5 | n.d. | 9.7 | 5.0 | n.d. | 8 | n.d. | n.d. |
| B03w | 1.50 | 0.5 | 1.8 | 1.4 | n.d. | 9.5 | 5.2 | 12.3 | 10.4 | n.d. | n.d. |
| B04w | 1.50 | 0.5 | 1.7 | 1.4 | n.d. | 9.4 | 9.3 | n.d. | 12.4 | n.d. | n.d. |
| B05w | 1.00 | 0.5 | 1.9 | 1.5 | n.d. | 9.6 | 14.5 | 9.5 | 18.9 | n.d. | n.d. |
| B06w | 1.68 | 0.5 | 1.8 | 1.4 | n.d. | 9.7 | 14.5 | 10.1 | 6.6 | n.d. | n.d. |
| B07w | 1.30 | 0.5 | 1.9 | 1.5 | n.d. | 9.7 | 20.1 | 8.6 | 14.5 | n.d. | n.d. |
| B08w | 1.30 | 0.5 | 1.9 | 1.6 | n.d. | 9.0 | 18.2 | 9.4 | 19 | n.d. | n.d. |
| B09w | 0.80 | 0.5 | 1.9 | 1.5 | n.d. | 8.8 | 22.7 | 8.1 | 13.2 | n.d. | n.d. |
| B010w | 1.30 | 0.5 | 2.0 | 1.6 | n.d. | 9.0 | 15.6 | 9.4 | 14.1 | n.d. | n.d. |
| B011w | 1.20 | 0.5 | 1.9 | 1.5 | n.d. | 9.0 | 9.8 | 10.7 | 4.7 | n.d. | n.d. |
| KH1w | 1.00 | 0.5 | 2.1 | 1.7 | n.d. | 9.2 | 9.9 | n.d. | 2.3 | n.d. | n.d. |
| KH2w | 0.80 | 0.5 | 2.1 | 1.7 | n.d. | 9.1 | 4.8 | n.d. | 1.5 | n.d. | n.d. |
| KH3w | 1.02 | 0.5 | 1.7 | 1.4 | n.d. | 9.1 | 5.9 | 8.0 | 3.7 | n.d. | n.d. |
| KH4w | 1.00 | 0.5 | 1.8 | 1.5 | n.d. | 8.7 | 11.0 | n.d. | 1.5 | n.d. | n.d. |
| KH5w | 1.20 | 0.5 | 1.9 | 1.5 | n.d. | 9.1 | 15.1 | 5.5 | 2.8 | n.d. | n.d. |
| KH6w | 1.08 | 0.5 | 1.9 | 1.5 | n.d. | 9.1 | 15.5 | 6.7 | 2.8 | n.d. | n.d. |
| KH7w | 0.80 | 0.5 | 2.1 | 1.7 | n.d. | 8.6 | 20.7 | 5.6 | 4 | n.d. | n.d. |

| | | | | | | | | | | | |
|---|---|---|---|---|---|---|---|---|---|---|---|
| KH8w | 0.80 | 0.5 | 2.4 | 1.9 | n.d. | 8.4 | 20.2 | 5.3 | 4.6 | n.d. | n.d. |
| KH9w | 0.65 | 0.5 | 2.5 | 2.0 | n.d. | 8.6 | 23.4 | 7.3 | 2.7 | n.d. | n.d. |
| KH10w | 0.90 | 0.5 | 2.2 | 1.8 | n.d. | 8.7 | 16.0 | 5.4 | 8.3 | n.d. | n.d. |
| KH11w | 0.85 | 0.5 | 2.3 | 1.8 | n.d. | 8.4 | 10.1 | 4.2 | 3.8 | n.d. | n.d. |
| B01 | 0.09 | 0.01 | 6.4 | 5.1 | n.d. | 9.1 | 17.9 | 11.0 | 319.5 | n.d. | 34.9634 |
| B02 | 0.14 | 0.01 | 6.3 | 5.0 | n.d. | 9.4 | 15.7 | 9.0 | 375 | n.d. | 28.2496 |
| B03 | 0.09 | 0.01 | 6.6 | 5.3 | n.d. | 9.2 | 26.3 | 10.0 | 238.5 | n.d. | 29.8441 |
| B04 | 0.05 | 0.01 | 8.1 | 6.5 | n.d. | 9.5 | 25.2 | 8.4 | 105.1 | n.d. | 37.0405 |
| B05 | 0.10 | 0.01 | 12.7 | 10.2 | n.d. | 9.5 | 26.1 | 8.0 | 107.9 | n.d. | 61.7976 |
| B06 | 0.12 | 0.01 | 11.6 | 9.2 | n.d. | 9.8 | 21.4 | 6.9 | 204.5 | n.d. | 51.7479 |
| B07 | 0.07 | 0.01 | 19.1 | 15.3 | n.d. | 9.6 | 24.1 | 8.5 | 230 | n.d. | 446.876 |
| B08 | 0.10 | 0.01 | 11.2 | 9.0 | n.d. | 9.4 | 22.3 | 10.1 | 153.4 | n.d. | 126.061 |
| B09 | 0.05 | 0.01 | 15.5 | 12.4 | n.d. | 9.8 | 24.4 | 11.3 | 85.4 | n.d. | 78.708 |
| B10 | 0.01 | 0.01 | 29.1 | 23.3 | n.d. | 9.7 | 25.7 | 9.8 | 13.2 | n.d. | 1266.38 |
| B11 | 0.04 | 0.01 | 23.5 | 18.8 | n.d. | 9.7 | 24.8 | 16.0 | 264.2 | n.d. | 545.275 |
| B12 | 0.12 | 0.01 | 7.3 | 5.8 | n.d. | 9.8 | 18.2 | 8.6 | 51.1 | n.d. | 29.0259 |
| B13 | 0.09 | 0.01 | 10.8 | 8.6 | n.d. | 9.6 | 17.4 | 12.2 | 255.7 | n.d. | 598.6 |
| B14 | 0.09 | 0.01 | 6.0 | 4.8 | n.d. | 9.6 | 6.0 | 11.4 | n.d. | n.d. | 428.9 |
| K01 | 0.15 | 0.01 | 3.2 | 2.6 | n.d. | 8.9 | 14.1 | 8.5 | 247.2 | n.d. | 20.8854 |
| K02 | 0.14 | 0.01 | 3.1 | 2.4 | n.d. | 9.2 | 16.6 | 10.0 | 251.4 | n.d. | 19.8783 |
| K03 | 0.17 | 0.01 | 3.1 | 2.4 | n.d. | 9.2 | 22.5 | 9.1 | 541.7 | n.d. | 20.172 |
| K04 | 0.13 | 0.01 | 4.0 | 3.2 | n.d. | 9.1 | 27.8 | 7.1 | 89.5 | n.d. | 32.2569 |
| K05 | 0.04 | 0.01 | 7.0 | 5.6 | n.d. | 9.4 | 30.3 | 8.4 | 306.7 | n.d. | 52.1255 |
| K06 | 0.02 | 0.01 | 2.6 | 2.0 | n.d. | 9.8 | 24.8 | 6.1 | 362.2 | n.d. | 10.0804 |
| K09 | 0.05 | 0.01 | 8.8 | 7.0 | n.d. | 9.8 | 27.8 | 14.2 | 498.4 | n.d. | 69.9381 |
| K12 | 0.02 | 0.01 | 5.1 | 4.1 | n.d. | 9.4 | 19.4 | 9.9 | 267.1 | n.d. | 33.6626 |
| K14 | 0.11 | 0.01 | 3.3 | 2.6 | n.d. | 9.4 | 7.1 | 10.6 | n.d. | n.d. | 306.3 |
| S01 | 0.44 | 0.01 | 5.2 | 4.1 | n.d. | 8.5 | 15.9 | 6.5 | 268.4 | n.d. | 83.6 |
| S02 | 0.40 | 0.01 | 5.2 | 4.1 | n.d. | 8.9 | 16.2 | 6.9 | 264.2 | n.d. | 87.8975 |
| S03 | 0.42 | 0.01 | 5.1 | 4.1 | n.d. | 8.7 | 24.0 | 4.0 | 117.1 | n.d. | 83.8273 |
| S04 | 0.37 | 0.01 | 6.1 | 4.9 | n.d. | 8.7 | 24.6 | 4.2 | 78.8 | n.d. | 107.493 |
| S05 | 0.25 | 0.01 | 9.1 | 7.3 | n.d. | 9.2 | 25.9 | 3.8 | 49.2 | n.d. | 162.358 |
| S06 | 0.25 | 0.01 | 8.5 | 6.8 | n.d. | 9.5 | 24.9 | 6.9 | 195.9 | n.d. | 156.672 |
| S07 | 0.23 | 0.01 | 12.5 | 10.0 | n.d. | 9.5 | 26.5 | 9.6 | 257.7 | n.d. | 243.364 |
| S08 | 0.46 | 0.01 | 1.1 | 0.9 | n.d. | 9.7 | 20.3 | 9.8 | 164.3 | n.d. | 212.313 |
| S09 | 0.08 | 0.01 | 10.4 | 8.4 | n.d. | 9.8 | 25.5 | 14.7 | 183.3 | n.d. | 217.977 |
| S10 | 0.07 | 0.01 | 15.8 | 12.6 | n.d. | 9.7 | 27.0 | 12.5 | 106.5 | n.d. | 359.387 |
| S11 | 0.10 | 0.01 | 18.9 | 15.1 | n.d. | 9.5 | 25.4 | 15.2 | 52.3 | n.d. | 352.883 |
| S12 | 0.15 | 0.01 | 6.9 | 5.5 | n.d. | 9.4 | 19.9 | 3.9 | 81 | n.d. | 161.414 |
| S13 | 0.15 | 0.01 | 8.6 | 6.9 | n.d. | 9.2 | 18.8 | 6.6 | 57.9 | n.d. | 800.5 |
| S14 | 0.20 | 0.01 | 4.7 | 3.8 | n.d. | 9.3 | 6.9 | 10.1 | n.d. | n.d. | 377.7 |
| V01 | 0.28 | 0.01 | 5.1 | 4.0 | n.d. | 9.3 | 13.7 | 8.5 | 15.4 | n.d. | 32.0681 |
| V02 | 0.25 | 0.01 | 4.9 | 4.0 | n.d. | 9.4 | 15.2 | 9.7 | 2.9 | n.d. | 31.04 |
| V03 | 0.27 | 0.01 | 5.0 | 4.0 | n.d. | 9.4 | 21.2 | 8.2 | 1.8 | n.d. | 34.8585 |
| V04 | 0.23 | 0.01 | 6.2 | 4.9 | n.d. | 9.2 | 23.9 | 3.8 | 1.9 | n.d. | 46.7125 |
| V05 | 0.12 | 0.01 | 10.0 | 8.0 | n.d. | 9.5 | 29.4 | 11.7 | 3.4 | n.d. | 76.6099 |
| V06 | 0.10 | 0.01 | 6.7 | 5.4 | n.d. | 9.7 | 22.1 | 8.8 | 112.5 | n.d. | 53.6362 |
| V07 | 0.05 | 0.01 | 11.7 | 9.4 | n.d. | 9.8 | 25.4 | 9.4 | 498.5 | n.d. | 276.304 |

| | | | | | | | | | | | |
|---|---|---|---|---|---|---|---|---|---|---|---|
| V08 | 0.15 | 0.01 | 10.0 | 8.0 | n.d. | 9.6 | 21.2 | 10.1 | 426.1 | n.d. | 96.3947 |
| V09 | 0.04 | 0.01 | 11.2 | 9.0 | n.d. | 9.7 | 28.1 | 12.3 | 672.6 | n.d. | 367.15 |
| V10 | 0.02 | 0.01 | 16.4 | 13.1 | n.d. | 9.9 | 25.9 | 5.0 | n.d. | n.d. | 1281.7 |
| V11 | 0.12 | 0.01 | 14.3 | 11.4 | n.d. | 9.6 | 25.2 | 10.5 | 696.7 | n.d. | 820.961 |
| V12 | 0.11 | 0.01 | 6.8 | 5.5 | n.d. | 9.7 | 18.9 | 10.3 | 28.6 | n.d. | 49.419 |
| V13 | 0.06 | 0.01 | 8.9 | 7.1 | n.d. | 9.6 | 19.5 | 8.7 | 26.9 | n.d. | 710.9 |
| V14 | 0.16 | 0.01 | 5.6 | 4.5 | n.d. | 9.7 | 6.7 | 11.6 | n.d. | n.d. | 496.1 |
| Z01 | 0.10 | 0.01 | 4.3 | 3.4 | n.d. | 9.2 | 15.4 | 11.5 | 319.5 | n.d. | 23.7178 |
| Z02 | 0.26 | 0.01 | 4.3 | 3.4 | n.d. | 9.6 | 15.3 | 9.2 | 323.8 | n.d. | 27.1376 |
| Z03 | 0.15 | 0.01 | 4.2 | 3.4 | n.d. | 9.4 | 23.7 | 9.9 | 204.5 | n.d. | 26.0047 |
| Z04 | 0.20 | 0.01 | 5.1 | 4.1 | n.d. | 9.5 | 24.2 | 8.8 | 8.5 | n.d. | 28.5433 |
| Z05 | 0.15 | 0.01 | 7.3 | 5.8 | n.d. | 9.5 | 30.9 | 10.3 | 93.7 | n.d. | 45.8943 |
| Z06 | 0.07 | 0.01 | 7.6 | 6.1 | n.d. | 9.7 | 24.0 | 8.1 | 255.6 | n.d. | 45.0761 |
| Z07 | 0.22 | 0.01 | 11.6 | 9.3 | n.d. | 9.7 | 23.6 | 8.0 | 136.3 | n.d. | 326.028 |
| Z08 | 0.05 | 0.01 | 9.3 | 7.4 | n.d. | 9.8 | 19.6 | 9.0 | 598.6 | n.d. | 54.853 |
| Z09 | 0.08 | 0.01 | 12.9 | 10.3 | n.d. | 9.8 | 27.2 | 9.0 | 51.3 | n.d. | 610.945 |
| Z10 | 0.02 | 0.01 | 34.7 | 27.8 | n.d. | 10.0 | 24.9 | 12.3 | 588.7 | n.d. | 3341.15 |
| Z12 | 0.04 | 0.01 | 6.4 | 5.1 | n.d. | 9.7 | 21.1 | 12.5 | 12.8 | n.d. | 35.9285 |
| Z14 | 0.06 | 0.01 | 3.6 | 2.8 | n.d. | 9.6 | 5.9 | 11.1 | n.d. | n.d. | 208.8 |
| BV | 3.00 | 0.1 | n.d. | n.d. | 0.5 | 8.6 | n.d. | n.d. | n.d. | n.d. | n.d. |
| K0 | 2.60 | 0.1 | 482.0 | 0.4 | n.d. | 7.5 | 9.8 | 7.2 | 5.96 | 20.8 | 20.4 |
| K01 | 2.60 | 1 | 494.0 | 0.4 | n.d. | 7.6 | 9.7 | 7.1 | 7.09 | 22.5 | 21.5 |
| A | 2.10 | 0.05 | 0.5 | 0.4 | n.d. | 7.9 | 24.6 | 1.3 | 194.7 | n.d. | n.d. |
| B | 1.00 | 0.05 | 0.5 | 0.4 | n.d. | 8.0 | 24.7 | 1.9 | 115 | n.d. | n.d. |
| B0 | 1.30 | 0.5 | 2.6 | 2.1 | n.d. | 8.7 | 28.5 | 6.6 | 5.3 | n.d. | n.d. |
| C | 1.40 | 0.05 | 0.5 | 0.4 | n.d. | 7.7 | 25.5 | 2.8 | 50.4 | n.d. | n.d. |
| D | 2.10 | 0.05 | 0.5 | 0.4 | n.d. | 7.8 | 25.3 | 1.8 | 28.9 | n.d. | n.d. |
| Ds | 4.05 | 0.05 | 0.6 | 0.5 | n.d. | 8.6 | 28.1 | 7.0 | 70.7 | n.d. | n.d. |
| E | 6.00 | 0.05 | 0.6 | 0.5 | n.d. | 8.3 | 29.0 | 8.9 | 56.4 | n.d. | n.d. |
| F | 0.48 | 0.05 | 0.6 | 0.5 | n.d. | 8.5 | 29.3 | 9.1 | 54.7 | n.d. | n.d. |
| FDN | 0.65 | 0.05 | 0.6 | 0.5 | n.d. | 8.6 | 30.8 | 10.8 | 34.4 | n.d. | n.d. |
| FDO | 1.80 | 0.05 | 0.6 | 0.5 | n.d. | 8.6 | 30.5 | 10.9 | 34.1 | n.d. | n.d. |
| Ff | 4.50 | 0.05 | 0.8 | 0.6 | n.d. | 8.7 | 28.1 | 7.8 | 1.4 | n.d. | n.d. |
| K | 3.10 | 0.05 | 0.7 | 0.6 | n.d. | 8.6 | 27.3 | 7.4 | 6.1 | n.d. | n.d. |
| KH | 0.75 | 0.5 | 2.9 | 2.3 | n.d. | 8.7 | 28.9 | 6.4 | 16.7 | n.d. | n.d. |
| Kn | 3.50 | 0.05 | n.d. | n.d. | n.d. | 8.7 | 29.0 | 17.7 | 116.3 | n.d. | n.d. |
| LK | 0.72 | 0.05 | 0.4 | 0.3 | n.d. | 7.5 | 28.0 | 2.3 | 6.2 | n.d. | n.d. |
| LN | 0.25 | 0.05 | 0.4 | 0.3 | n.d. | 7.5 | 27.0 | 2.5 | 10.5 | n.d. | n.d. |
| NH | 1.00 | 0.5 | 2.8 | 2.2 | n.d. | 8.8 | 30.1 | 7.4 | 18.7 | n.d. | n.d. |
| R | 0.75 | 0.05 | 0.6 | 0.5 | n.d. | 9.7 | 31.3 | 14.5 | 140.2 | n.d. | n.d. |
| TLO | 1.60 | 0.05 | 0.7 | 0.5 | n.d. | 8.6 | 29.0 | 9.7 | 88.2 | n.d. | n.d. |
| TNB | 1.00 | 0.05 | 0.5 | 0.4 | n.d. | 7.4 | 28.2 | 2.8 | 17.2 | n.d. | n.d. |
| TO | 1.85 | 0.05 | 0.5 | 0.4 | n.d. | 8.2 | 27.5 | 7.1 | 31.1 | n.d. | n.d. |
| TS | 1.00 | 0.05 | 0.5 | 0.4 | n.d. | 7.8 | 28.4 | 4.2 | 22.3 | n.d. | n.d. |
| Z | 4.20 | 0.05 | 0.6 | 0.5 | n.d. | 7.6 | 24.3 | 0.3 | 3.4 | n.d. | n.d. |
| SS15 | n.d. | 0.01 | n.d. | n.d. | n.d. | n.d. | 0.5 | n.d. | n.d. | n.d. | n.d. |
| SS16 | n.d. | 0.01 | n.d. | n.d. | n.d. | n.d. | 0.0 | n.d. | n.d. | n.d. | n.d. |
| SS17 | n.d. | 0.01 | n.d. | n.d. | n.d. | n.d. | 8.5 | n.d. | n.d. | n.d. | n.d. |

| | | | | | | | | | | | |
|---|---|---|---|---|---|---|---|---|---|---|---|
| US15 | n.d. | 0.01 | n.d. | n.d. | n.d. | n.d. | 0.5 | n.d. | n.d. | n.d. | n.d. |
| US16 | n.d. | 0.01 | n.d. | n.d. | n.d. | n.d. | 0.0 | n.d. | n.d. | n.d. | n.d. |
| US17 | n.d. | 0.01 | n.d. | n.d. | n.d. | n.d. | 8.5 | n.d. | n.d. | n.d. | n.d. |
| ZL15 | n.d. | 0.01 | 12.9 | 10.3 | n.d. | 9.5 | 0.6 | n.d. | 193.9 | n.d. | n.d. |
| ZL16 | n.d. | 0.01 | n.d. | n.d. | n.d. | n.d. | 0.0 | n.d. | n.d. | n.d. | n.d. |
| ZL17 | n.d. | 0.01 | 9.0 | 7.2 | n.d. | 9.6 | 8.9 | n.d. | 299.2 | n.d. | n.d. |
| Sp17_01 | 0.04 | 0.02 | 6.7 | 5.4 | n.d. | 9.4 | 14.5 | 9.9 | 111.6 | n.d. | n.d. |
| Sp17_02 | 0.08 | 0.02 | 2.1 | 1.7 | n.d. | 8.7 | 13.1 | 10.4 | 174 | n.d. | n.d. |
| Sp17_04 | 0.05 | 0.02 | 2.2 | 1.8 | n.d. | 8.8 | 12.1 | 10.6 | 318.2 | n.d. | n.d. |
| Sp17_05 | 0.30 | 0.02 | 2.6 | 2.0 | n.d. | 9.1 | 15.6 | 20.1 | 108.7 | n.d. | n.d. |
| Sp17_07 | 0.12 | 0.02 | 6.7 | 5.4 | n.d. | 8.9 | 11.1 | 10.8 | 350.7 | n.d. | n.d. |
| Sp17_09 | 0.06 | 0.02 | 4.2 | 3.4 | n.d. | 9.3 | 9.5 | 11.6 | 318.6 | n.d. | n.d. |
| Sp17_10 | 0.02 | 0.02 | 4.5 | 3.6 | n.d. | 8.9 | 14.2 | 10.1 | 77 | n.d. | n.d. |
| Sp17_11 | 0.32 | 0.02 | 3.5 | 2.8 | n.d. | 9.2 | 20.0 | 9.0 | 177.2 | n.d. | n.d. |
| Sp17_12 | 0.03 | 0.02 | 3.0 | 2.4 | n.d. | 9.2 | 21.5 | 8.6 | 203.8 | n.d. | n.d. |
| Sp17_13 | 0.15 | 0.02 | 7.2 | 5.8 | n.d. | 9.5 | 16.2 | 10.1 | 38.4 | n.d. | n.d. |
| Sp17_14 | 0.09 | 0.02 | 4.1 | 3.3 | n.d. | 8.6 | 14.6 | 12.3 | 21.9 | n.d. | n.d. |
| Sp17_15 | 0.06 | 0.02 | 3.6 | 2.9 | n.d. | 9.1 | 13.7 | 10.1 | 90.4 | n.d. | n.d. |
| Sp17_16 | 0.06 | 0.02 | 8.5 | 6.8 | n.d. | 9.7 | 11.8 | 11.2 | 13.1 | n.d. | n.d. |
| Sp17_17 | 0.05 | 0.02 | 8.7 | 7.0 | n.d. | 9.6 | 17.5 | 10.8 | 168.3 | n.d. | n.d. |
| Sp17_18 | 0.06 | 0.02 | 4.2 | 3.3 | n.d. | 8.9 | 12.0 | 11.0 | 585.9 | n.d. | n.d. |
| Sp17_19 | 0.12 | 0.02 | 2.7 | 2.2 | n.d. | 8.8 | 12.7 | 10.3 | 351.1 | n.d. | n.d. |
| Sp17_20 | 0.22 | 0.02 | 3.0 | 2.4 | n.d. | 8.6 | 12.9 | 11.0 | 3.4 | n.d. | n.d. |
| Sp17_21 | 0.08 | 0.02 | 9.7 | 7.8 | n.d. | 9.6 | 15.8 | 10.6 | 39.2 | n.d. | n.d. |
| Sp17_22 | 0.05 | 0.02 | 8.2 | 6.6 | n.d. | 8.9 | 13.1 | 10.4 | 90 | n.d. | n.d. |
| Sp17_23 | 0.04 | 0.02 | 2.3 | 1.8 | n.d. | 8.9 | 13.4 | 11.2 | 182.3 | n.d. | n.d. |
| Sp17_24 | 0.12 | 0.02 | 11.0 | 8.8 | n.d. | 9.8 | 13.3 | 11.3 | 1 | n.d. | n.d. |
| Sp17_27 | 0.03 | 0.02 | 6.1 | 4.9 | n.d. | 9.0 | 12.3 | 10.7 | 190.9 | n.d. | n.d. |
| Sp17_28 | 0.25 | 0.02 | 3.7 | 2.9 | n.d. | 8.7 | 12.6 | 10.5 | 16.3 | n.d. | n.d. |
| Sp17_29 | 0.10 | 0.02 | 5.8 | 4.6 | n.d. | 9.5 | 13.6 | 10.9 | 43.6 | n.d. | n.d. |
| Sp17_31 | 0.30 | 0.02 | 2.6 | 2.0 | n.d. | 9.1 | 15.6 | 20.1 | 108.7 | n.d. | n.d. |
| Sp17_Neu2 | 0.05 | 0.02 | 4.4 | 3.5 | n.d. | 9.0 | 13.8 | 11.3 | 130.7 | n.d. | n.d. |
| Sp18_01 | 0.18 | 0.02 | 1.5 | 1.2 | n.d. | 9.0 | 20.3 | 12.8 | 20.5 | n.d. | n.d. |
| Sp18_04 | 0.15 | 0.02 | 0.7 | 0.5 | n.d. | 8.7 | 16.8 | 10.3 | 16.2 | n.d. | n.d. |
| Sp18_05 | 0.26 | 0.02 | 2.2 | 1.7 | n.d. | 9.0 | 11.6 | 8.9 | 52.2 | n.d. | n.d. |
| Sp18_07 | 0.22 | 0.02 | 1.3 | 1.0 | n.d. | 8.7 | 17.8 | 10.3 | 7.6 | n.d. | n.d. |
| Sp18_08 | 0.21 | 0.02 | 0.7 | 0.6 | n.d. | 8.8 | 11.0 | 10.7 | 44 | n.d. | n.d. |
| Sp18_10 | 0.12 | 0.02 | 0.8 | 0.7 | n.d. | 8.6 | 20.3 | 9.9 | 1.1 | n.d. | n.d. |
| Sp18_11 | 0.28 | 0.02 | 1.7 | 1.4 | n.d. | 8.7 | 14.2 | 10.9 | 16.6 | n.d. | n.d. |
| Sp18_12 | 0.17 | 0.02 | 0.7 | 0.6 | n.d. | 8.5 | 12.3 | 10.4 | 2.9 | n.d. | n.d. |
| Sp18_13 | 0.24 | 0.02 | 3.4 | 2.7 | n.d. | 9.2 | 11.9 | 10.6 | 131.3 | n.d. | n.d. |
| Sp18_15 | 0.36 | 0.02 | 2.1 | 1.7 | n.d. | 8.7 | 10.5 | 77.2 | 60.2 | n.d. | n.d. |
| Sp18_16 | 0.15 | 0.02 | 2.5 | 2.0 | n.d. | 9.1 | 14.5 | 96.0 | 102.2 | n.d. | n.d. |
| Sp18_17 | 0.14 | 0.02 | 3.4 | 2.7 | n.d. | 9.3 | 11.4 | 102.8 | 238.3 | n.d. | n.d. |
| Sp18_18 | 0.17 | 0.02 | 1.2 | 0.9 | n.d. | 8.9 | 13.6 | 101.8 | 14.6 | n.d. | n.d. |
| Sp18_19 | 0.17 | 0.02 | 0.6 | 0.5 | n.d. | 8.6 | 19.2 | 116.5 | 101.1 | n.d. | n.d. |
| Sp18_1_d | 0.23 | 0.02 | 0.8 | 0.7 | n.d. | 8.6 | 18.5 | 10.0 | 5.9 | n.d. | n.d. |
| Sp18_20 | 0.44 | 0.02 | 2.5 | 2.0 | n.d. | 8.8 | 15.1 | 11.2 | 2.7 | n.d. | n.d. |

| | | | | | | | | | | | |
|---|---|---|---|---|---|---|---|---|---|---|---|
| Sp18_21 | 0.30 | 0.02 | 3.9 | 3.1 | n.d. | 9.1 | 14.8 | 10.7 | 82.3 | n.d. | n.d. |
| Sp18_22 | 0.21 | 0.02 | 1.8 | 1.4 | n.d. | 8.9 | 17.0 | 11.2 | 56.2 | n.d. | n.d. |
| Sp18_23 | 0.12 | 0.02 | 0.6 | 0.4 | n.d. | 8.7 | 18.7 | 11.1 | 31.7 | n.d. | n.d. |
| Sp18_24 | 0.48 | 0.02 | 4.2 | 3.4 | n.d. | 9.3 | 18.7 | 12.0 | 11.6 | n.d. | n.d. |
| Sp18_27 | 0.18 | 0.02 | 1.2 | 1.0 | n.d. | 8.8 | 17.1 | 9.9 | 17.4 | n.d. | n.d. |
| Sp18_28 | 0.39 | 0.02 | 2.3 | 1.8 | n.d. | 8.7 | 13.8 | 10.4 | 5 | n.d. | n.d. |
| Sp18_29 | 0.26 | 0.02 | 2.1 | 1.7 | n.d. | 9.2 | 17.7 | 10.9 | 33.1 | n.d. | n.d. |
| Sp18_30 | 0.35 | 0.02 | 4.1 | 3.3 | n.d. | 9.1 | 13.6 | 10.6 | 37.9 | n.d. | n.d. |
| Sp18_31 | 0.31 | 0.02 | 9.1 | 7.3 | n.d. | 9.2 | 17.8 | 9.3 | 44.2 | n.d. | n.d. |
| Sp18_9 | 0.28 | 0.02 | 1.2 | 1.0 | n.d. | 8.7 | 11.1 | 9.9 | 74.7 | n.d. | n.d. |
| Sp18_Neu2 | 0.12 | 0.02 | 1.0 | 0.8 | n.d. | 9.1 | 22.0 | 9.5 | 25.3 | n.d. | n.d. |
| 01Boddi | 0.12 | 0.01 | 6.2 | 5.0 | n.d. | 9.2 | 19.9 | 9.2 | n.d. | n.d. | n.d. |
| 02Soser | 0.42 | 0.01 | 4.8 | 3.8 | n.d. | 8.7 | 20.5 | 4.3 | n.d. | n.d. | n.d. |
| 03KolonNy | 1.85 | 0.01 | 0.5 | 0.4 | n.d. | 7.9 | 21.2 | 6.3 | n.d. | n.d. | n.d. |
| 04KolonT18 | 1.60 | 0.01 | 0.3 | 0.2 | n.d. | 7.2 | 21.2 | 6.6 | n.d. | n.d. | n.d. |
| 05Szelid18 | 4.70 | 0.01 | 2.2 | 1.8 | n.d. | 8.7 | 20.0 | 6.9 | n.d. | n.d. | n.d. |
| 06Rusanda18 | 0.15 | 0.01 | 18.7 | 15.0 | n.d. | 9.6 | 23.9 | 8.1 | n.d. | n.d. | n.d. |
| 07SlanoKopovo18 | 0.20 | 0.01 | 4.9 | 3.9 | n.d. | 9.0 | 19.0 | 8.7 | n.d. | n.d. | n.d. |
| 08VBK18 | 0.42 | 0.01 | 4.3 | 3.4 | n.d. | 9.5 | 20.0 | 9.3 | n.d. | n.d. | n.d. |
| 09Kelemen18 | 0.17 | 0.01 | 2.6 | 2.1 | n.d. | 9.3 | 21.4 | 8.7 | n.d. | n.d. | n.d. |
| 10Zab18 | 0.38 | 0.01 | 3.7 | 3.0 | n.d. | 9.5 | 26.0 | 8.6 | n.d. | n.d. | n.d. |
| 12KH18 | 0.89 | 0.01 | 2.0 | 1.6 | n.d. | 8.4 | 23.7 | 5.9 | n.d. | n.d. | n.d. |
| 13FertoB018 | 1.55 | 0.01 | 1.8 | 1.4 | n.d. | 9.0 | 23.1 | 8.1 | n.d. | n.d. | n.d. |
| 14VelenceiKo18 | 2.23 | 0.01 | 3.2 | 2.6 | n.d. | 8.9 | 22.2 | 9.1 | n.d. | n.d. | n.d. |
| 15Dinnyesi18 | 0.71 | 0.01 | 3.0 | 2.4 | n.d. | 9.0 | 24.3 | 13.9 | n.d. | n.d. | n.d. |
| 16VelenceiNy18 | 1.72 | 0.01 | 2.5 | 2.0 | n.d. | 8.1 | 23.1 | 3.3 | n.d. | n.d. | n.d. |
| 17Zicklacke18 | 0.10 | 0.01 | 4.6 | 3.7 | n.d. | 9.7 | 28.9 | 14.1 | n.d. | n.d. | n.d. |
| 18ObererStinker18 | 0.08 | 0.01 | 9.4 | 7.5 | n.d. | 9.5 | 25.2 | 10.5 | n.d. | n.d. | n.d. |
| 19SudlicherSilbersee18 | 0.20 | 0.01 | 9.1 | 7.3 | n.d. | 9.6 | 27.3 | 14.8 | n.d. | n.d. | n.d. |
| 20LangeLacke18 | 0.11 | 0.01 | 4.2 | 3.4 | n.d. | 9.3 | 25.1 | 8.7 | n.d. | n.d. | n.d. |
| 21BalatonTihany18 | 4.20 | 0.01 | 0.8 | 0.6 | n.d. | 8.7 | 22.6 | 7.9 | n.d. | n.d. | n.d. |
| 22BalatonKeszthely18 | 3.20 | 0.01 | 0.7 | 0.6 | n.d. | 8.6 | 21.3 | 7.9 | n.d. | n.d. | n.d. |
| 23UntererStinker18 | 0.24 | 0.01 | 4.8 | 3.8 | n.d. | 10.0 | 24.8 | 10.3 | n.d. | n.d. | n.d. |
| 01T | 0.04 | 0.01 | 17.5 | n.d. | 12.3 | 9.3 | 18.7 | 10.4 | 5.5 | 42.7 | 35.9 |
| 02T | 0.06 | 0.01 | 12.7 | n.d. | 11.1 | 9.5 | 23.3 | 13.4 | 18.9 | 257.1 | 236.8 |
| 03T | 0.06 | 0.01 | 3.8 | n.d. | 3.1 | 9.4 | 18.6 | 10.6 | 23.1 | 68.7 | 37.3 |
| 04T | 0.23 | 0.01 | 2.1 | n.d. | 1.8 | 9.7 | 16.3 | 11.2 | 2.9 | 40.3 | 37.6 |
| 05T | 0.31 | 0.01 | 2.0 | n.d. | 1.6 | 9.1 | 16.8 | 15.8 | 102.8 | 51.3 | 41.2 |
| 06T | 0.13 | 0.01 | 3.7 | n.d. | 2.9 | 9.3 | 23.9 | 9.2 | 19.2 | 159.3 | 136.4 |
| 07T | 0.11 | 0.01 | 7.2 | n.d. | 5.5 | 9.6 | 23.8 | 9.4 | 136.1 | 71.8 | 39.6 |
| 08T | 0.11 | 0.01 | 6.2 | n.d. | 4.6 | 9.7 | 23.3 | 9.1 | 60.9 | 54 | 24.3 |
| 09T | 0.12 | 0.01 | 1.9 | n.d. | 1.5 | 9.5 | 21.6 | 9.0 | 25.5 | 90.6 | 14.6 |
| 10T | 0.05 | 0.01 | 4.2 | n.d. | 3.3 | 9.5 | 19.0 | 9.5 | 32.4 | 32.8 | 28.7 |
| 11T | 0.07 | 0.01 | 1.5 | n.d. | 1.5 | 9.5 | 14.1 | 10.1 | 27.9 | 61.2 | 18.6 |
| 12T | 0.11 | 0.01 | 1.7 | n.d. | 1.5 | 9.7 | 14.5 | 10.4 | 15.3 | 27.1 | 14.4 |
| 13T | 0.05 | 0.01 | 2.0 | n.d. | 1.8 | 9.6 | 17.0 | 9.9 | 20.9 | 56.9 | 42.5 |
| 14T | 0.15 | 0.01 | 5.6 | n.d. | 5.0 | 9.1 | 24.2 | 8.1 | 0 | 51.6 | 10 |
| 15T | 0.13 | 0.01 | 5.7 | n.d. | 5.1 | 9.0 | 26.3 | 8.2 | 0 | 66.6 | 49.8 |

| | | | | | | | | | | | |
|---|---|---|---|---|---|---|---|---|---|---|---|
| 16T | 0.15 | 0.01 | 0.5 | n.d. | 0.3 | 8.3 | 21.8 | 10.6 | 14 | 5.7 | 4 |
| 17T | 0.22 | 0.01 | 1.3 | n.d. | 1.2 | 8.2 | 23.4 | 4.4 | 34.4 | 65.9 | 46.4 |
| 18T | 0.24 | 0.01 | 0.8 | n.d. | 0.6 | 8.1 | 25.7 | 7.8 | 1.9 | 13.9 | 13.3 |
| 19T | 0.25 | 0.01 | 4.2 | n.d. | 3.6 | 9.0 | 21.2 | 8.7 | 5.1 | 41 | 37.7 |
| 20T | 0.05 | 0.01 | 8.5 | n.d. | 8.0 | 9.4 | 23.9 | 11.4 | 0 | 141 | 88 |
| 21T | 1.20 | 0.01 | 2.8 | n.d. | 2.3 | 8.7 | 24.6 | 9.1 | 4.7 | 36 | 30 |
| 22T | 1.48 | 0.01 | 2.8 | n.d. | 2.2 | 8.8 | 24.6 | 9.9 | 20.8 | 26.5 | 24.9 |
| 23T | 1.48 | 0.01 | 2.8 | n.d. | 2.3 | 8.4 | 25.6 | 9.5 | 16.8 | 24.8 | 21 |
| 24T | 1.03 | 0.01 | 2.8 | n.d. | 2.3 | 8.5 | 25.2 | 8.5 | 6.4 | 33.2 | 27.5 |
| 25T | 1.10 | 0.01 | 2.9 | n.d. | 2.5 | 8.9 | 24.7 | 9.9 | 21.3 | 30 | 23 |
| 26T | 4.50 | 0.01 | 1.6 | n.d. | 1.1 | 8.5 | 26.8 | 9.6 | 10.9 | 12.4 | 11.7 |
| 27T | 4.00 | 0.01 | 1.6 | n.d. | 1.2 | 8.5 | 27.8 | 9.5 | 12.6 | 14.8 | 12.1 |
| 28T | 1.75 | 0.01 | 4.7 | n.d. | 4.2 | 9.0 | 29.2 | 9.7 | 13.9 | 61.7 | 38.7 |
| 29T | 1.48 | 0.01 | 4.5 | n.d. | 4.1 | 8.9 | 28.8 | 8.5 | 12.3 | 41.3 | 39.4 |
| 30T | 1.12 | 0.01 | 4.4 | n.d. | 3.9 | 8.9 | 28.1 | 7.0 | 28.7 | 45.4 | 43.9 |
| 31T | 0.45 | 0.01 | 4.7 | n.d. | 4.2 | 9.3 | 29.5 | 11.9 | 407.1 | 79.8 | 24.6 |
| DL-M Day 0 | n.d. | 0.05 | 22.2 | n.d. | 22.9 | 10.1 | n.d. | n.d. | n.d. | n.d. | n.d. |
| DLM 2015 | n.d. | 0.05 | 22.2 | n.d. | 22.9 | 10.1 | n.d. | n.d. | n.d. | n.d. | n.d. |
| DLM 2017 | n.d. | 0.05 | 19.8 | n.d. | 16.7 | 10.4 | n.d. | n.d. | n.d. | n.d. | n.d. |
| GEL-M Day 0 | n.d. | 0.05 | 33.1 | n.d. | 36.1 | 10.1 | n.d. | n.d. | n.d. | n.d. | n.d. |
| GEM 2015 | n.d. | 0.05 | 33.1 | n.d. | 36.1 | 10.1 | n.d. | n.d. | n.d. | n.d. | n.d. |
| GEM 2016 | n.d. | 0.05 | 33.1 | n.d. | 36.1 | 10.1 | n.d. | n.d. | n.d. | n.d. | n.d. |
| GEM 2017 | n.d. | 0.05 | 16.4 | n.d. | 13.8 | 10.7 | n.d. | n.d. | n.d. | n.d. | n.d. |
| LCL-M Day 0 | n.d. | 0.05 | 44.2 | n.d. | 48.3 | 10.1 | n.d. | n.d. | n.d. | n.d. | n.d. |
| LCM 2015 | n.d. | 0.05 | 44.2 | n.d. | 48.3 | 10.1 | n.d. | n.d. | n.d. | n.d. | n.d. |
| LCM 2016 | n.d. | 0.05 | 44.2 | n.d. | 48.3 | 10.1 | n.d. | n.d. | n.d. | n.d. | n.d. |
| LCM 2017 | n.d. | 0.05 | 46.5 | n.d. | 45.4 | 10.3 | n.d. | n.d. | n.d. | n.d. | n.d. |
| PL-M Day 0 | n.d. | 0.05 | 22.9 | n.d. | 23.8 | 10.1 | n.d. | n.d. | n.d. | n.d. | n.d. |
| PLM 2015 | n.d. | 0.05 | 22.9 | n.d. | 23.8 | 10.1 | n.d. | n.d. | n.d. | n.d. | n.d. |
| PLM 2016 | n.d. | 0.05 | 22.9 | n.d. | 23.8 | 10.1 | n.d. | n.d. | n.d. | n.d. | n.d. |
| PLM 2017 | n.d. | 0.05 | 19.3 | n.d. | 16.7 | 10.4 | n.d. | n.d. | n.d. | n.d. | n.d. |
| ML16S_10m | 41.00 | 1 | n.d. | n.d. | 77.6 | 9.8 | 17.2 | 5.2 | n.d. | n.d. | n.d. |
| MLW_0517_00 | 41.00 | 0.01 | n.d. | n.d. | 85.0 | 9.8 | n.d. | n.d. | n.d. | n.d. | n.d. |
| MLW_0517_10 | 41.00 | 10 | n.d. | n.d. | 85.0 | 9.8 | n.d. | n.d. | n.d. | n.d. | n.d. |
| MLW_0617_00 | 41.00 | 0.01 | n.d. | n.d. | 85.0 | 9.8 | n.d. | n.d. | n.d. | n.d. | n.d. |
| MLW_0917_00_1;MLW_0917_00_2;MLW_0917_00_3 | 41.00 | 0.01 | n.d. | n.d. | 75.0 | 9.8 | n.d. | n.d. | n.d. | n.d. | n.d. |
| MLW_0917_05_1;MLW_0917_05_2;MLW_0917_05_3 | 41.00 | 5 | n.d. | n.d. | 86.0 | 9.8 | n.d. | n.d. | n.d. | n.d. | n.d. |
| MLW_0917_10_1;MLW_0917_10_2;MLW_0917_10_3 | 41.00 | 10 | n.d. | n.d. | 86.0 | 9.8 | n.d. | n.d. | n.d. | n.d. | n.d. |
| MLW_1018_00 | 41.00 | 0.01 | n.d. | n.d. | 80.0 | 9.8 | n.d. | n.d. | n.d. | n.d. | n.d. |
| MLW_1018_05 | 41.00 | 5 | n.d. | n.d. | 80.0 | 9.8 | n.d. | n.d. | n.d. | n.d. | n.d. |
| MLW_1018_12 | 41.00 | 12 | n.d. | n.d. | 80.0 | 9.8 | n.d. | n.d. | n.d. | n.d. | n.d. |

n.d. - not determined
*defined according to Boros & Kolpakova 2018

| Sample ID | Reference for environmental variables | Note |
|---|---|---|
| Bac-12S | Fazi et al., 2018; 2021 | |
| Bac-18E | Fazi et al., 2018;Lameck et al., 2023 | |
| Bac-6M | Fazi et al., 2018; Getenet et al., 2022 | |
| B1-W | Jirsa et al., 2013 | the two read set from Lake Bogoria merged together during the bioinformatic analysis |
| B2-W | Jirsa et al., 2013 | the two read set from Lake Bogoria merged together during the bioinformatic analysis |
| KZ02 | Boros et al., 2017 | |
| K03bact | Boros et al., 2017 | |
| KZ04 | Boros et al., 2017 | |
| KZ05 | this study | |
| K06bact | Boros et al., 2017 | |
| KZ08 | Boros et al., 2017 | |
| K10bact | Boros et al., 2017 | |
| KZ12 | Boros et al., 2017 | |
| K13bact | Boros et al., 2017 | |
| K15bact | Boros et al., 2017 | |
| K16bact | Boros et al., 2017 | |
| K18bact | this study | |
| KZ19 | Boros et al., 2017 | |
| K20bact | this study | |
| KT01 | this study | |
| KT02 | this study | |
| KT03 | this study | |
| KT04 | this study | |
| KT05 | this study | |
| KT06 | this study | |
| KT07 | this study | |
| KT08 | this study | |
| KT09 | this study | |
| KT10 | this study | |
| KT11 | this study | |
| KT12 | this study | |
| KT13 | this study | salinity values derived from conductivity based on the region specific equation ($R^2$ = 0.91): $salinity_{EC}$ = 1.1 * EC - 2.92 |
| KT14 | this study | salinity values derived from conductivity based on the region specific equation ($R^2$ = 0.91): $salinity_{EC}$ = 1.1 * EC - 2.92 |
| KT15 | this study | salinity values derived from conductivity based on the region specific equation ($R^2$ = 0.91): $salinity_{EC}$ = 1.1 * EC - 2.92 |
| KT16 | this study | |
| KT17 | this study | salinity values derived from conductivity based on the region specific equation ($R^2$ = 0.91): $salinity_{EC}$ = 1.1 * EC - 2.92 |
| KT18 | this study | |
| KT19 | this study | |
| KT20 | this study | |
| KT21 | this study | |
| KT22 | this study | |
| KT23 | this study | |
| KT24 | this study | |
| KT25 | this study | |
| KT26 | this study | |
| KT27 | this study | |
| 0S | this study | |
| 02S | this study | |
| 04S | this study | |
| 06S | this study | |
| 08S | this study | |
| 09S | this study | |
| 11S | this study | |

| | | |
|---|---|---|
| SHV | this study | |
| KA1 | this study | |
| KA2 | this study | |
| KA3 | this study | |
| KA4 | this study | |
| KA5 | this study | |
| KA6 | this study | |
| KA7 | this study | |
| KA8 | this study | |
| Gudzh_water | Lavrentyeva et al., 2020 | |
| NB30 | Matyugina et al., 2018 | |
| NB31 | Matyugina et al., 2018 | |
| NB32 | Matyugina et al., 2018 | |
| NB41 | Matyugina et al., 2018 | |
| NB42 | Matyugina et al., 2018 | |
| NB43 | Matyugina et al., 2018 | |
| Bangong Co | Ji et al., 2019, Yue et al., 2019 | |
| Bong Co | Ji et al., 2019, Yue et al., 2019 | |
| Co Ngoin (1) | Ji et al., 2019, Yue et al., 2019 | |
| Mapam Yumco | Ji et al., 2019, Yue et al., 2019 | |
| Urru Co | Ji et al., 2019, Yue et al., 2019 | |
| Bam Co | Ji et al., 2019, Yue et al., 2019 | |
| Bero Zeco | Ji et al., 2019, Yue et al., 2019 | |
| Co Ngoin (2) | Ji et al., 2019, Yue et al., 2019 | |
| Dawa Co | Ji et al., 2019, Yuan et al., 2011 | |
| Tangra Yumco | Ji et al., 2019, Yue et al., 2019 | |
| Dong Co | Ji et al., 2019, Günther et al., 2014 | |
| Kunggyu | Ji et al., 2019, Yue et al., 2019 | |
| Nam Co | Ji et al., 2019; Wang2010 | Salinity published by Ji et al. (2019) is slightly below our 1 g/L threshold, however other sources reported elevated salinities for Nam Co (1.1 - 2.9 g/L), therefore we considered this site as 'soda' |
| Pung Co | Ji et al., 2019, Yue et al., 2019 | |
| Selin Co | Ji et al., 2019, Yue et al., 2019 | |
| Zhari Namco | Ji et al., 2019, Yue et al., 2019 | |
| Zhaxi Co | Ji et al., 2019, Yue et al., 2019 | |
| Bangkog Co | Ji et al., 2019, Yue et al., 2019 | |
| Dangqiong Co | Ji et al., 2019 | |
| Nyer Co (Nieer Co) | Ji et al., 2019, Ye et al., 2016 | pH value is based on the average of 8 sample given in Ye et al., 2016 |
| BU1 | Szabo et al., 2017 | |
| SO1 | Szabo et al., 2017 | |
| ZA1 | Szabo et al., 2017 | |
| SO2 | Szabo et al., 2020 | |
| SO3 | Szabo et al., 2020 | |
| SO4 | Szabo et al., 2020 | |
| SO5 | Szabo et al., 2020 | |
| SO6 | Szabo et al., 2020 | |
| SO9 | Szabo et al., 2020 | |
| SO11 | Szabo et al., 2020 | |
| SO12 | Szabo et al., 2020 | |
| SO13 | Szabo et al., 2020 | |
| SO14 | Szabo et al., 2020 | |
| SO15 | Szabo et al., 2020 | |
| SO16 | Szabo et al., 2020 | |
| VBK | Korponai et al., 2019 | |
| ZA2 | Szabo et al., 2020 | |
| ZA3 | Szabo et al., 2020 | |

| | |
|---|---|
| ZA4 | Szabo et al., 2020 |
| ZA5 | Szabo et al., 2020 |
| ZA6 | Szabo et al., 2020 |
| ZA7 | Szabo et al., 2020 |
| ZA8 | Szabo et al., 2020 |
| ZA9 | Szabo et al., 2020 |
| ZA10 | Szabo et al., 2020 |
| ZA11 | Szabo et al., 2020 |
| ZA12 | Szabo et al., 2020 |
| ZA13 | Szabo et al., 2020 |
| ZA14 | Szabo et al., 2020 |
| ZA15 | Szabo et al., 2020 |
| ZA16 | Szabo et al., 2020 |
| ZA17 | Szabo et al., 2020 |
| M05 | this study |
| U05 | this study |
| U15 | this study |
| U28 | this study |
| B01w | Szuroczki et al., 2020 |
| B02w | this study |
| B03w | this study |
| B04w | this study |
| B05w | this study |
| B06w | this study |
| B07w | this study |
| B08w | Szuroczki et al., 2020 |
| B09w | this study |
| B010w | this study |
| B011w | this study |
| KH1w | Szuroczki et al., 2020 |
| KH2w | this study |
| KH3w | this study |
| KH4w | this study |
| KH5w | this study |
| KH6w | this study |
| KH7w | this study |
| KH8w | Szuroczki et al., 2020 |
| KH9w | this study |
| KH10w | this study |
| KH11w | this study |
| B01 | Márton et al., 2023 |
| B02 | Márton et al., 2023 |
| B03 | Márton et al., 2023 |
| B04 | Márton et al., 2023 |
| B05 | Márton et al., 2023 |
| B06 | Márton et al., 2023 |
| B07 | Márton et al., 2023 |
| B08 | Márton et al., 2023 |
| B09 | Márton et al., 2023 |
| B10 | Márton et al., 2023 |
| B11 | Márton et al., 2023 |
| B12 | Márton et al., 2023 |
| B13 | Márton et al., 2023 |
| B14 | Márton et al., 2023 |

| | |
|---|---|
| K01 | Márton et al., 2023 |
| K02 | Márton et al., 2023 |
| K03 | Márton et al., 2023 |
| K04 | Márton et al., 2023 |
| K05 | Márton et al., 2023 |
| K06 | Márton et al., 2023 |
| K09 | Márton et al., 2023 |
| K12 | Márton et al., 2023 |
| K14 | Márton et al., 2023 |
| S01 | Márton et al., 2023 |
| S02 | Márton et al., 2023 |
| S03 | Márton et al., 2023 |
| S04 | Márton et al., 2023 |
| S05 | Márton et al., 2023 |
| S06 | Márton et al., 2023 |
| S07 | Márton et al., 2023 |
| S08 | Márton et al., 2023 |
| S09 | Márton et al., 2023 |
| S10 | Márton et al., 2023 |
| S11 | Márton et al., 2023 |
| S12 | Márton et al., 2023 |
| S13 | Márton et al., 2023 |
| S14 | Márton et al., 2023 |
| V01 | Márton et al., 2023 |
| V02 | Márton et al., 2023 |
| V03 | Márton et al., 2023 |
| V04 | Márton et al., 2023 |
| V05 | Márton et al., 2023 |
| V06 | Márton et al., 2023 |
| V07 | Márton et al., 2023 |
| V08 | Márton et al., 2023 |
| V09 | Márton et al., 2023 |
| V10 | Márton et al., 2023 |
| V11 | Márton et al., 2023 |
| V12 | Márton et al., 2023 |
| V13 | Márton et al., 2023 |
| V14 | Márton et al., 2023 |
| Z01 | Márton et al., 2023 |
| Z02 | Márton et al., 2023 |
| Z03 | Márton et al., 2023 |
| Z04 | Márton et al., 2023 |
| Z05 | Márton et al., 2023 |
| Z06 | Márton et al., 2023 |
| Z07 | Márton et al., 2023 |
| Z08 | Márton et al., 2023 |
| Z09 | Márton et al., 2023 |
| Z10 | Márton et al., 2023 |
| Z12 | Márton et al., 2023 |
| Z14 | Márton et al., 2023 |
| BV | Szilágyi2005 |
| K0 | Mentes et al., 2018 |
| K01 | Mentes et al., 2018 |
| A | this study |
| B | this study |

| | | |
|---|---|---|
| B0 | this study | |
| C | this study | |
| D | this study | |
| Ds | this study | |
| E | this study | |
| F | this study | |
| FDN | this study | coordinates of the lake: 46.44978 N 18.85512 E |
| FDO | this study | coordinates of the lake: 46.44978 N 18.85512 E |
| Ff | this study | |
| K | this study | |
| KH | this study | |
| Kn | this study | |
| LK | this study | |
| LN | this study | |
| NH | this study | coordinates of the lake: 47.688386 N 16.713290 E |
| R | this study | |
| TLO | this study | coordinates of the lake: 46.42737 N 18.79895 E |
| TNB | this study | coordinates of the lake: 46.42737 N 18.79895 E |
| TO | this study | |
| TS | this study | |
| Z | this study | |
| SS15 | Siclair et al., 2015 | |
| SS16 | Siclair et al., 2015 | |
| SS17 | Siclair et al., 2015 | |
| US15 | Siclair et al., 2015 | |
| US16 | Siclair et al., 2015 | |
| US17 | Siclair et al., 2015 | |
| ZL15 | Siclair et al., 2015 | |
| ZL16 | Siclair et al., 2015 | |
| ZL17 | Siclair et al., 2015 | |
| Sp17_01 | Szabó et al., 2022 | |
| Sp17_02 | Szabó et al., 2022 | |
| Sp17_04 | Szabó et al., 2022 | |
| Sp17_05 | Szabó et al., 2022 | |
| Sp17_07 | Szabó et al., 2022 | |
| Sp17_09 | Márton et al., 2023b | |
| Sp17_10 | Szabó et al., 2022 | |
| Sp17_11 | Szabó et al., 2022 | |
| Sp17_12 | Szabó et al., 2022 | |
| Sp17_13 | Szabó et al., 2022 | |
| Sp17_14 | Márton et al., 2023b | |
| Sp17_15 | Szabó et al., 2022 | |
| Sp17_16 | Szabó et al., 2022 | |
| Sp17_17 | Szabó et al., 2022 | |
| Sp17_18 | Szabó et al., 2022 | |
| Sp17_19 | Szabó et al., 2022 | |
| Sp17_20 | Szabó et al., 2022 | |
| Sp17_21 | Szabó et al., 2022 | |
| Sp17_22 | Szabó et al., 2022 | |
| Sp17_23 | Szabó et al., 2022 | |
| Sp17_24 | Szabó et al., 2022 | |
| Sp17_27 | Szabó et al., 2022 | |
| Sp17_28 | Szabó et al., 2022 | |
| Sp17_29 | Szabó et al., 2022 | |

| | |
|---|---|
| Sp17_31 | Márton et al., 2023b |
| Sp17_Neu2 | Szabó et al., 2022 |
| Sp18_01 | Szabó et al., 2022 |
| Sp18_04 | Szabó et al., 2022 |
| Sp18_05 | Szabó et al., 2022 |
| Sp18_07 | Szabó et al., 2022 |
| Sp18_08 | Márton et al., 2023b |
| Sp18_10 | Szabó et al., 2022 |
| Sp18_11 | Szabó et al., 2022 |
| Sp18_12 | Szabó et al., 2022 |
| Sp18_13 | Szabó et al., 2022 |
| Sp18_15 | Szabó et al., 2022 |
| Sp18_16 | Szabó et al., 2022 |
| Sp18_17 | Szabó et al., 2022 |
| Sp18_18 | Szabó et al., 2022 |
| Sp18_19 | Szabó et al., 2022 |
| Sp18_1_d | Szabó et al., 2022 |
| Sp18_20 | Szabó et al., 2022 |
| Sp18_21 | Szabó et al., 2022 |
| Sp18_22 | Szabó et al., 2022 |
| Sp18_23 | Szabó et al., 2022 |
| Sp18_24 | Szabó et al., 2022 |
| Sp18_27 | Szabó et al., 2022 |
| Sp18_28 | Szabó et al., 2022 |
| Sp18_29 | Szabó et al., 2022 |
| Sp18_30 | Márton et al., 2023b |
| Sp18_31 | Márton et al., 2023b |
| Sp18_9 | Márton et al., 2023b |
| Sp18_Neu2 | Szabó et al., 2022 |
| 01Boddi | this study |
| 02Soser | this study |
| 03KolonNy | this study |
| 04KolonT18 | this study |
| 05Szelid18 | this study |
| 06Rusanda18 | this study |
| 07SlanoKopovo18 | this study |
| 08VBK18 | this study |
| 09Kelemen18 | this study |
| 10Zab18 | this study |
| 12KH18 | this study |
| 13FertoB018 | this study |
| 14VelenceiKo18 | this study |
| 15Dinnyesi18 | this study |
| 16VelenceiNy18 | this study |
| 17Zicklacke18 | this study |
| 18ObererStinker18 | this study |
| 19SudlicherSilbersee18 | this study |
| 20LangeLacke18 | this study |
| 21BalatonTihany18 | this study |
| 22BalatonKeszthely18 | this study |
| 23UntererStinker18 | this study |
| 01T | this study |
| 02T | this study |
| 03T | this study |

| | | |
|---|---|---|
| 04T | this study | |
| 05T | this study | |
| 06T | this study | |
| 07T | this study | |
| 08T | this study | |
| 09T | this study | |
| 10T | this study | |
| 11T | this study | |
| 12T | this study | |
| 13T | this study | |
| 14T | this study | |
| 15T | this study | |
| 16T | this study | |
| 17T | this study | |
| 18T | this study | |
| 19T | this study | |
| 20T | this study | |
| 21T | this study | |
| 22T | this study | |
| 23T | this study | |
| 24T | this study | |
| 25T | this study | |
| 26T | this study | |
| 27T | this study | |
| 28T | this study | |
| 29T | this study | |
| 30T | this study | |
| 31T | this study | |
| DL-M Day 0 | Zorz et al., 2019 | soda lake microbial mats; for instances where data were not determined, the measurements from 2015, as presented in Zorz et al., 2019, were used in the analysis |
| DLM 2015 | Zorz et al., 2019 | soda lake microbial mats |
| DLM 2017 | Zorz et al., 2019 | soda lake microbial mats |
| GEL-M Day 0 | Zorz et al., 2019 | soda lake microbial mats; for instances where data were not determined, the measurements from 2015, as presented in Zorz et al., 2019, were used |
| GEM 2015 | Zorz et al., 2019 | soda lake microbial mats |
| GEM 2016 | Zorz et al., 2019 | soda lake microbial mats; for instances where data were not determined, the measurements from 2015, as presented in Zorz et al., 2019, were used |
| GEM 2017 | Zorz et al., 2019 | soda lake microbial mats |
| LCL-M Day 0 | Zorz et al., 2019 | soda lake microbial mats; for instances where data were not determined, the measurements from 2015, as presented in Zorz et al., 2019, were used |
| LCM 2015 | Zorz et al., 2019 | soda lake microbial mats |
| LCM 2016 | Zorz et al., 2019 | soda lake microbial mats; for instances where data were not determined, the measurements from 2015, as presented in Zorz et al., 2019, were used |
| LCM 2017 | Zorz et al., 2019 | soda lake microbial mats |
| PL-M Day 0 | Zorz et al., 2019 | soda lake microbial mats; for instances where data were not determined, the measurements from 2015, as presented in Zorz et al., 2019, were used |
| PLM 2015 | Zorz et al., 2019 | soda lake microbial mats |
| PLM 2016 | Zorz et al., 2019 | soda lake microbial mats; for instances where data were not determined, the measurements from 2015, as presented in Zorz et al., 2019, were used |
| PLM 2017 | Zorz et al., 2019 | soda lake microbial mats |
| ML16S_10m | Edwardson & Hollibaugh, 2018 | |
| MLW_0517_00 | Philips et al., 2021 | |
| MLW_0517_10 | Philips et al., 2021 | |
| MLW_0617_00 | Philips et al., 2021 | |
| MLW_0917_00_1;MLW_0917_00_2;MLW_0917_00_3 | Philips et al., 2021 | |
| MLW_0917_05_1;MLW_0917_05_2;MLW_0917_05_3 | Philips et al., 2021 | |
| MLW_0917_10_1;MLW_0917_10_2;MLW_0917_10_3 | Philips et al., 2021 | |
| MLW_1018_00 | Philips et al., 2021 | |
| MLW_1018_05 | Philips et al., 2021 | |
| MLW_1018_12 | Philips et al., 2021 | |
